# Triptans reprogram Schwann cells to drive medication-overuse headache via β-glycan/TGF-β3 signaling

**DOI:** 10.64898/2026.04.24.720547

**Authors:** Martina Chieca, Matilde Marini, Marta Baragli, Elisa Bellantoni, Lorenzo Bonacchi, Francesco De Cesaris, Cristina Tassorelli, Roberto De Icco, Rosaria Greco, Lucrezia Timotei, Beatrice Pulli, Giuseppe Spinelli, Daniel Souza Monteiro de Araujo, Alice Papini, Gaetano De Siena, Irene Scuffi, Gabriele Ferroni, Giovanna Pivotto, Alberto Magi, Romina Nassini, Francesco De Logu

## Abstract

Medication-overuse headache (MOH) is one of the leading causes of chronic daily headache worldwide and arises from the repeated use of acute anti-migraine medications, including triptans. However, the cellular substrates and intracellular pathways driving this paradoxical chronification remain unknown. Here, using Schwann cell-selective silencing of the 5-HT_1B/D_ receptor, we observed that acute triptan administration counteracts CGRP-induced, endosome-confined cAMP accumulation, preventing the development of periorbital mechanical allodynia in mice. In contrast, prolonged 5-HT_1B/D_ activation in Schwann cells induces epigenetic and transcriptomic dysregulation associated with MOH in mice. Specifically, intronic hypermethylation-driven overexpression of *BETAGLYCAN* promotes activation of a non-canonical TGF-β-dependent signaling cascade. The resulting TGF-β3 upregulation establishes a feed-forward loop that sustains proalgesic paracrine communication between Schwann cells and primary sensory neurons. Analysis of plasma levels from patients with MOH confirmed elevated TGF-β3 levels specifically associated with triptan-dependent MOH, supporting the translational relevance of these findings. Together, our data identify Schwann cell 5-HT_1B/D_ signaling as a dual mediator of both the acute anti-migraine efficacy and the maladaptive mechanisms underlying MOH. These results provide a conceptual framework for strategies aimed at preserving the therapeutic benefits of triptans while limiting the adverse consequences of chronic administration.

**One Sentence Summary:** Schwann cell 5-HT1B/D receptor signaling mediates the dual effects of triptans by acutely inhibiting CGRP-driven nociceptive pathways while chronically promoting epigenetically driven TGF-β3-dependent proalgesic signaling that causes medication-overuse headache.

## Introduction

Migraine is a disabling and highly prevalent neurological disorder, affecting approximately 14-17% of the global population(*1*). Migraine primarily affects young women in their reproductive and most productive age. The International Headache Society defines chronic migraine as a condition characterized by 15 or more headache days per month, with at least 8 days with migraine features, for at least three months(*2*). Migraine is much more than pain, indeed the attack is accompanied by several associated symptoms, including sensitivity to light(*3*), sound(*4*), and odors(*5*), nausea and vomiting, mood changes, and fatigue(*6*). Currently, acute anti-migraine medications include specific molecules, including 5-HT_1_ receptors agonists (triptans), calcitonin gene related peptide (CGRP) receptor antagonists, as well as non-specific drugs, like cyclooxygenase (COX) activity inhibitors(*7*). However, these treatments are often associated with adverse effects and bear the risk of evolution toward medication overuse headache (MOH). MOH is a well-known complication of the repetitive intake of acute drugs. Triptans are associated with an exposure-related risk of MOH that is 5 to 10 times higher than in not exposed migraine individualsa(*8*). MOH is one of the most disabling headache disorders, with a marked female predominance and a prevalence ranging from 0.5% to 2.6% in the general population(*9, 10*). It develops in individuals with an underlying primary headache disorder who overuse acute medications for at least three months(*11, 12*). This overuse leads to an increase in both the intensity and frequency of attacks, creating a vicious cycle of medication consumption and escalating headache frequency. Although the mechanisms underlying MOH are not fully understood, several factors have been implicated, including central sensitization, metabolic alterations, abnormal pain processing, genetic susceptibility, and epigenetic modifications(*13–18*).

Recently, we observed in mice that the activation of the CGRP receptor in Schwann cells in mice increased the endosomal cAMP levels and was associated with a heightened periorbital mechanical allodynia (PMA) in mice(*19*). Given that triptans, as 5-HT1B/D receptor agonists, inhibit cAMP synthesis through modulation of adenylate cyclase, we hypothesized that their therapeutic effects, as well as their contribution to MOH development, may involve signaling pathways in Schwann cells(*20*).

In parallel, accumulating evidence indicates that altered gene regulation contributes to the pathophysiology of migraine and MOH(*21–23*). For instance, DNA hypomethylation at the intronic CpG site (cg14377273) of the *HDAC4* gene in MOH subjects is associated with a reduction in monthly headache days following the withdrawal of acute medication(*24*), and hypermethylation of three genes (*CORIN*, *CCKBR* and *CLDN9*) in specific cell types is associated with MOH(*18*).

Here we showed that sumatriptan-stimulated human Schwann cells exhibited a time-dependent epigenomic and transcriptomic alterations involving the TGF-β receptor 3 (β-glycan) and the TGF-β3. We speculated that sumatriptan induces an epigenetically driven overexpression of β-glycan linked to the autocrine TGF-β3 non-canonical signaling in Schwann cells, establishing a feedforward mechanism characterized by further overexpression of TGF-β3. Repeated administration of triptans (sumatriptan, naratriptan and eletriptan), but not the COX inhibitor indomethacin, in mice induced PMA and non-evoked pain responses (facial grimacing) that were prevented by Schwann cell selective silencing of *Betaglycan*, *Tgfb3* and the DNA methyltransferase. Human data from patients with triptans dependent MOH confirmed elevated circulating TGF-β3 levels compared to healthy controls. In contrast, both patients and mice with indomethacin-induced MOH did not show involvement of the β-Glycan/TGF-β3 pathway, supporting that the activation of Schwann cell 5-HT₁_B/D_ receptors represents a specific mechanism underlying the induction and maintenance of triptan-dependent MOH. Finally, we observed that TGF-β3 played a paracrine role in pain signaling through the activation of primary sensory neurons. These findings highlight the critical role of Schwann cells in both the abortive action of triptans in migraine attacks and in triptan-induced MOH, providing a foundation for targeted therapies reducing triptan-related off-target effects.

## Results

### Schwann cell 5-HT₁_B/D_ modulates evoked and non-evoked pain responses induced by CGRP

The paradigm that G protein-coupled receptors (GPCR) signaling is confined to the plasma membrane has been challenged by evidence showing that GPCRs can sustain intracellular signaling from endosomes(*25, 26*). Recently, the sustained activity of the Schwann cell CGRP receptor in endosome was associated to the persistence of migraine pain(*19*). To investigate the role of Schwann cells in sumatriptan MOH, we first asked whether Schwann cells play a role in sumatriptan as an abortive treatment for migraine attacks. Immunohistochemistry, qPCR and RNAscope confirmed the expression of sumatriptan receptors 5-HT_1B_ and 5-HT_1D_ (5-HT_1B/D_) proteins and mRNA in primary cultured of mouse and human Schwann cells, as well as in S100-positive cells in human and mouse sciatic nerve tissues (Fig 1a-d and Fig. S1a-d).

**Figure 1.**
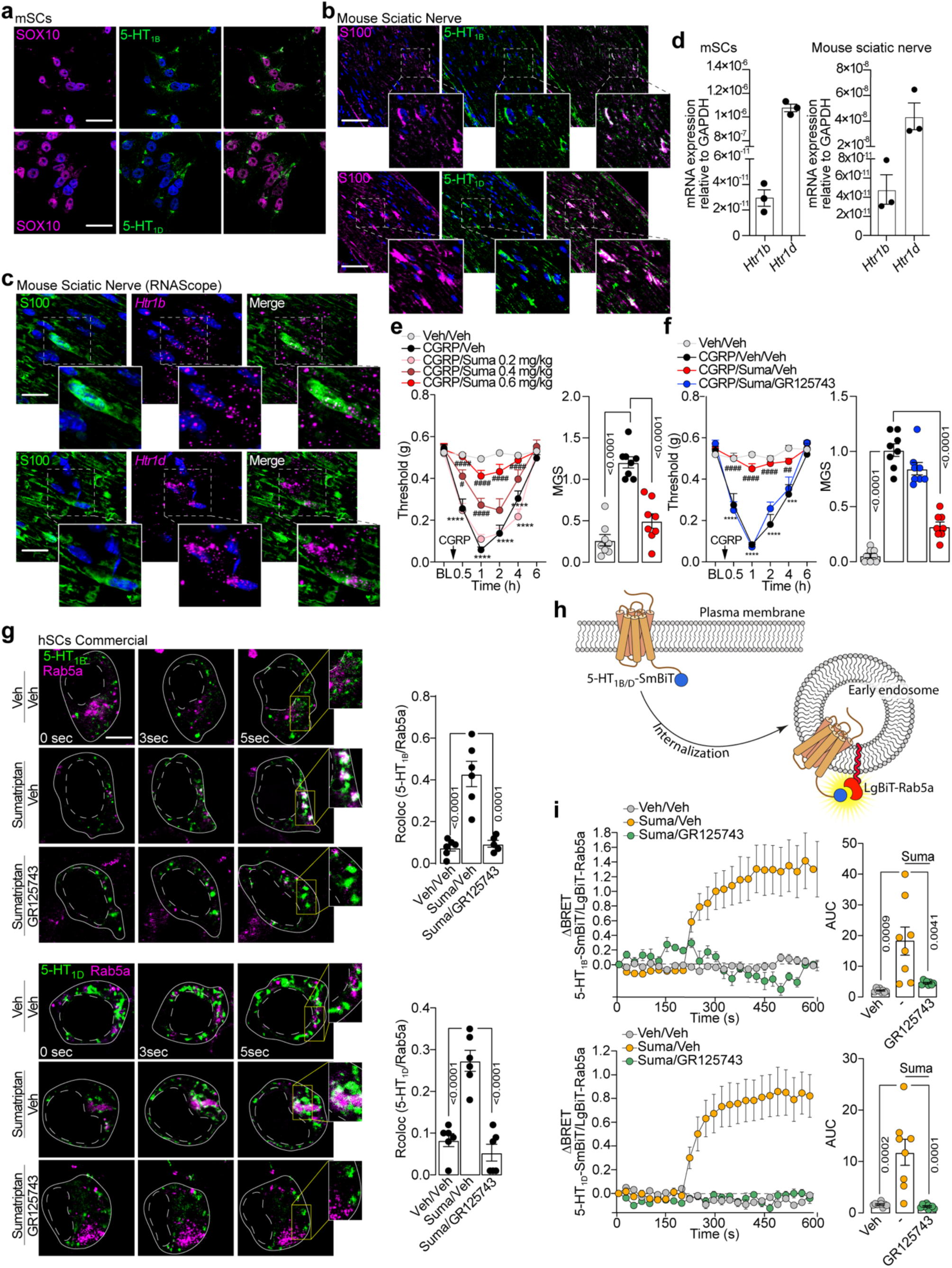
Schwann cells 5-HT_1B/1D_ control CGRP dependent pain. **(a)** Representative images of 5-HT_1B_ and 5-HT_1D_ expression in cultured primary mouse Schwann cells (mSCs, scale bar: 10 μm, n=6) and (**b)** in mouse sciatic nerve tissues (scale bar: 20μm, n=6). SOX10 and S100 are markers of SCs. **(c)** Representative RNAscope images of *Htr1b* and *Htr1d* mRNA and S100 protein expression in mouse sciatic nerve (scale bar: 5μm, n=6). **(d)** RT-qPCR for *Htr1b* and *Htr1d* mRNA in primary mSCs and mouse sciatic nerve tissues. **(e)** Dose dependent periorbital mechanical allodynia and cumulative data of facial grimacing (mouse grimace scale, MGS) in C57BL/6J mice treated with CGRP (0.1 mg/kg, i.p.) and pretreated with sumatriptan (Suma, 0.2-0.6 mg/kg) **(f)** and pretreated with GR125743 (0.5 mg/kg, i.p.) or vehicle (Veh). **(g)** Representative images and colocalization data (Rcoloc) of human SCs (hSCs) showing internalization of 5-HT_1B_-EGFP or 5-HT_1D_-EGFP in Ras-related protein (Rab5a)-RFP endosomes after sumatriptan exposure in presence of GR125743 or Veh (scale bar: 2μm n=6). **(h)** Schematic representation of 5-HT_1B/D_ trafficking to early endosomes using the NanoLuc Binary Technology (NanoBiT) assay. **(i)** Typical traces and cumulative data of 5-HT_1B_ and 5-HT_1D_ internalization after exposure to sumatriptan (10 µM) in presence of GR125743 (10 µM). (replicates: Veh=8, Suma=8, GR125743=8) Data are mean ± s.e.m. (n=8 mice/group). Student’s t-test, 1-way and 2-way ANOVA. ***p<0.001, ****p<0.0001 vs Veh/Veh; ^#^p<0.05, ^##^p<0.01, ^####^p<0.0001 vs CGRP/Veh or CGRP/Veh/Veh.

The role of Schwann cell 5-HT_1B/D_ receptors in mediating the analgesic effects of sumatriptan on CGRP-evoked pain was investigated *in vivo* in C57BL/6J (B6) mice. Specifically, we measured its impact on PMA and spontaneous non-evoked pain related behavior (assessed by the Grimace facial coding system) In male B6 mice, sumatriptan attenuated CGRP-mediated PMA in a dose-dependent manner (Fig. 1e). Sumatriptan higher dose (0.6 mg/kg) also reduced facial grimacing (Fig. 1e). The attenuation in the CGRP-evoked PMA and non-evoked response following sumatriptan was reversed by the pretreatment with the selective 5-HT_1B/D_ inhibitor, GR125743 (Fig. 1f).

The role of CGRP in long-lasting PMA in mice has been recently attributed to the ability of the CGRP receptor, the calcitonin-related polypeptide alpha/receptor activity modifying protein 1 (CALCA/RAMP1), to internalize in Schwann cell endosomes, along with adenylyl cyclase (AC), thus sustaining cAMP production over time(*19*). The 5-HT_1B/D_ receptors are GPCR coupled to G_i_ protein which inhibit AC decreasing intracellular cAMP levels(*27, 28*). Here, we investigated whether the analgesic effect of sumatriptan on CGRP-induced PMA and facial grimacing involves Schwann cell 5-HT_1B/D_ receptors. We first found that, sumatriptan stimulation of human Schwann cells promoted the internalization of EGFP-tagged 5-HT_1B/D_ receptors into RFP-marked early endosomes, which was prevented by the pretreatment with the 5-HT_1B/D_ receptors inhibitor, GR125743 (Fig. 1g). Receptor trafficking to early endosomes was further validated using the NanoLuc Binary Technology (NanoBiT) assay (Fig. 1h). Upon exposure to sumatriptan, Schwann cells displayed a marked increase in luminescence, which was abolished by pretreatment with GR125743 and absent in vehicle-treated cells (Fig. 1i).

To specifically monitor cAMP dynamics at early endosomes, we engineered a fusion construct of the early endosome marker Ras-related protein Rab5a and the circularly permuted GFP-based cAMP sensor G-Flamp2, generating G-Flamp2-Rab5a. CGRP stimulation of human Schwann cells induced a concentration dependent (3 nM-30 µM) and long lasting (approximately 300 seconds) endosomal cAMP increase which was inhibited by sumatriptan (100 nM-300 µM) (Fig. 2a-b). Pretreatment with GR125743 prevented the inhibitory effect of sumatriptan (Fig. 2c). These data were confirmed in primary Schwann cells isolated from human lingual, sublingual and inferior alveolar nerves biopsies and from mouse sciatic nerves (Fig. S1e-f). To minimize animal use in compliance with the 3Rs principle and given the rarity of obtaining human samples for Schwann cell isolation, we employed commercially available primary human Schwann cells for subsequent *in vitro* experiments.

**Figure 2.**
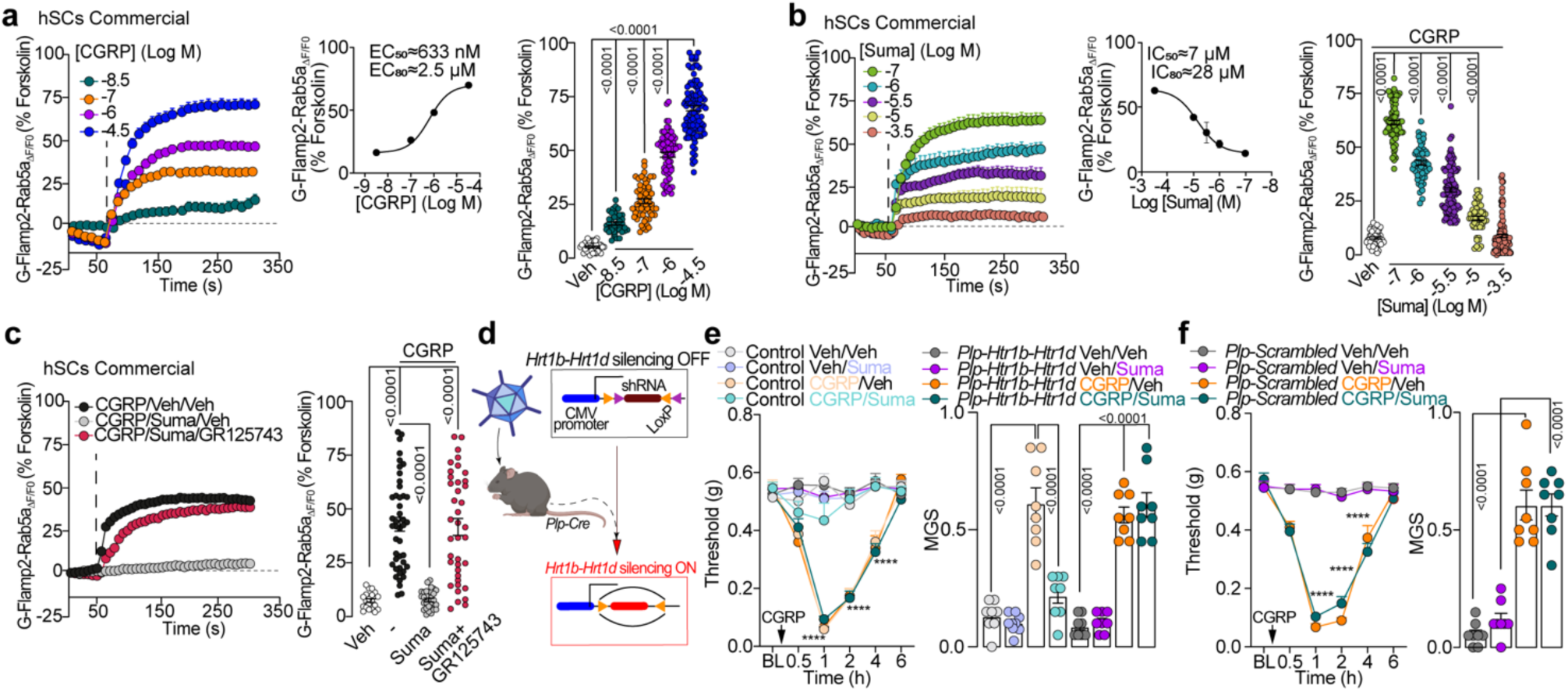
Schwann cell 5-HT_1B_/_1D_ mediate sumatriptan inhibition of CGRP signaling. **(a)** Typical traces, concentration response curve (CRC) and cumulative data showing the effects of graded concentrations of CGRP (3 nM-30 µM) or vehicle (Veh) on cAMP formation at early endosomes in human Schwann cells (hSCs) (cells number: 3 nM =38, 100 nM=54, 1 µM=70, 30 µM=57; EC_50_ CI=506-787 nM and EC_80_ CI=2-3.1 µM). Ras-related protein (Rab5a) is an early endosome marker. **(b)** Typical traces, CRC and cumulative data showing the effects of graded concentrations of sumatriptan (Suma, 100 nM-300 µM) or Veh on CGRP (3 µM)-evoked Rab5a associated cAMP formation in hSCs (cells number: 100 nM=75, 1 µM=55, 3 µM=108, 10 µM=53, 300 µM=109; IC_50_ CI=3.2-20 µM; IC_80_ CI=13-82 µM) and **(c)** in the presence of GR125743 (1 μM) or Veh (cells number: Veh/Veh/Veh=20, CGRP/Veh/Veh =44, CGRP/Suma/Veh =32, CGRP/Suma/GR125743 =39). **(d)** Schematic representation of Cre-dependent dicistronic adeno AAV strategy to selectively silence 5-HT_1B/D_ receptors in SCs in *Plp*^Cre^ mice. **(e)** Time dependent periorbital mechanical allodynia and facial grimacing (mouse grimace scale, MGS) induced by CGRP (0.1 mg/kg, i.p.) in Control, *Plp-Htr1b-Htr1d* and **(f)** *Plp-scrambled* mice pretreated with Suma (0.6 mg/kg, i.p.) or Veh. Data are mean ± s.e.m. (n=8 mice/group). 1-way and 2-way ANOVA, Bonferroni correction, ****p<0.0001 vs Control/Veh/Veh or *Plp-Scrambled*/Veh/Veh.

To investigate the role of Schwann cell 5-HT_1B/D_ in the modulation of CGRP-induced pain by sumatriptan we used a Cre-dependent dicistronic adeno associated viral vector (AAV)rh10 with *loxP*-flanked shRNA for Schwann cell selective 5-HT_1B/D_ silencing (shRNA-*Htr1b-Htr1d*) and a *Plp^Cre^* driver that functions as a lineage tracer to express shRNA-*Htr1b-Htr1d* in Schwann cells of *Plp^Cre^*mice (Fig. 2d). In *Plp-Htr1b-Htr1d* mice, sumatriptan did not prevent the CGRP dependent PMA and facial grimacing compared to Control mice or *Plp-Scrambled* mice (Fig. 2e-f). The efficiency of 5-HT_1B/D_ silencing in SCs in *Plp-Htr1b-Htr1d* mice was confirmed by immunohistochemistry (Fig. S1g).

### Sumatriptan drives MOH altering genes expression in Schwann cell

In humans, the chronic daily headache (15 or more headache days/month for more than 3 months) related to overuse of triptans (above 10 or more days) is defined MOH(*29*). To date the biological mechanism for MOH is not well understood. Here, we hypothesized that since Schwann cells play a role in the abortive mechanism of sumatriptan in CGRP-induced PMA, they might also be involved in triptan-induced MOH. First, we used a sumatriptan-induced MOH model in B6 mice(*30*). Repeated treatment with sumatriptan (0.6 mg/kg once daily over 7 days) in B6 mice induced time-dependent PMA and facial grimacing (Fig. 3a). Since female and male mice did not differ in PMA or non-evoked responses, only male mice were used for the MOH experiments (Fig. 3a). Both PMA and non-evoked responses induced by sumatriptan were blocked by pre-treatment with GR125743 (Fig. 3b). In addition, In *Plp-5-Htr1b-Htr1d* mice, PMA and facial grimacing were significantly reduced compared to Control or *Scrambled* mice (Fig. 3c), establishing a key role for Schwann cell 5-HT_1B/D_ receptors in sumatriptan-induced MOH in mice.

**Figure 3.**
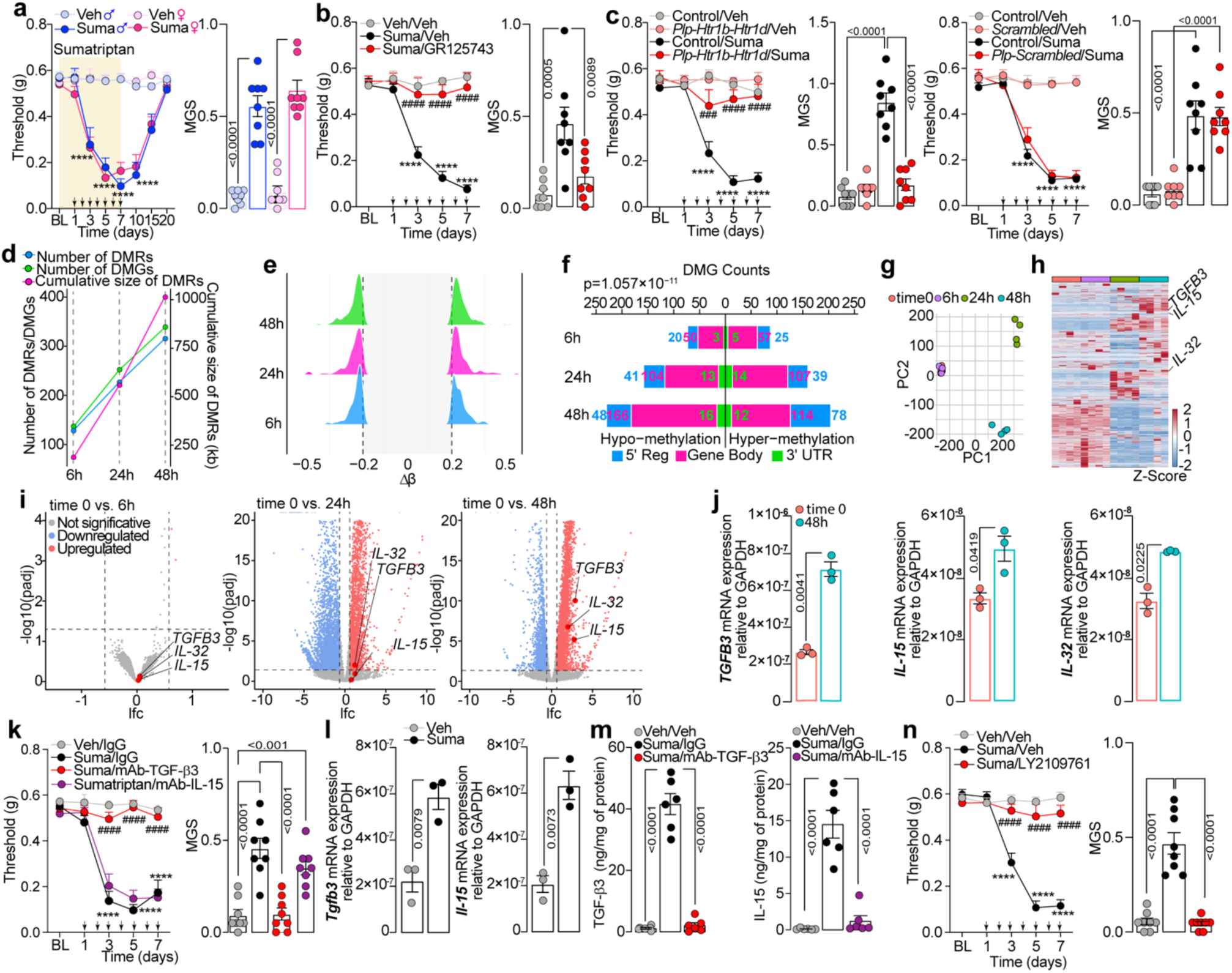
Repeated sumatriptan administration induces Schwann cell epigenetic and transcriptional dysregulation. **(a)**Time dependent periorbital mechanical allodynia (PMA) and facial grimacing (mouse grimace scale, MGS) in male and female C57BL/6J (B6) mice during/after repeated administration of sumatriptan (Suma, 0.6 mg/kg once daily over 7 consecutive days) or vehicle (Veh). **(b)** PMA and facial grimacing induced by Suma or Veh in B6 mice pretreated with GR125743 (0.5 mg/kg, i.p.) or Veh. **(c)** PMA and facial grimacing induced by Suma or Veh in *Plp-5-Htr1b-Htr1d,* Control or *Plp-Scrambled* mice. **(d)** Differentially methylated regions (DMRs), differentially methylated genes (DMGs) and the cumulative size of DMRs in human Schwann cells (hSCs) stimulated with Suma (10 µM) for 6h, 24h, and 48h. **(e)** Density plots of methylation changes distribution (Δβ) for all DMRs identified at 6h, 24h, and 48h. **(f)** Butterfly plot of the DMGs distribution across different genomic features Consistent with the global trend described in panel d (p=1.057×10^−11^). **(g)** Multidimensional scaling plot representing relationships between different time points after hSCs stimulation with Suma. Each point represents a single sample. **(h)** Heatmap of differentially expressed genes (DEGs) in hSCs across the indicated time points. Each column represents an individual sample. Conditions are ordered from left to right as time 0, 6h, 24h, and 48h. List of genes is available in Table S3 **(i)** Volcano plots showing the log2-fold change (lfc) and –log10(padj) for all DEGs between hSCs at time 0 vs. 6h, 24h and 48h of Suma stimulation. The top 200 differentially expressed genes are provided in the Table S4, while the complete list of genes is available in the Data file S1. **(j)** RT-qPCR for *TGFB3*, *IL-15* and *IL-32* mRNA in hSCs after Suma stimulation (48 h). (**k)** PMA and facial grimacing induced by Suma or Veh in B6 mice pretreated with mAb-TGF-β3, mAb-IL-15 (all, 200 μg/mice) or IgG. **(l)** RT-qPCR for *Tgfb3* and *Il-15* mRNA in mouse trigeminal nerve homogenates. **(m)** ELISA assay for TGF-β3 and IL-15 in trigeminal nerve tissue homogenates from B6 mice after Suma or Veh and pretreated with mAb-TGF-β3, mAb-IL-15 or IgG. **(n)** PMA and facial grimacing after Suma or Veh administration in B6 mice pretreated with LY2109761 (50 mg/kg, i.g.) or Veh. Black arrows indicate Suma administration. Data are mean ± s.e.m. (n=8 mice/group). Student’s t-test, 1-way and 2-way ANOVA, Bonferroni correction. ****p<0.0001 vs Veh♂, Veh♀, Veh/Veh, Control/Veh, Veh/IgG; ^###^p<0.001, ^####^p<0.0001 vs Suma/Veh, Control/Suma or Suma/IgG. Statistical significance was assessed **(d)** using the Cochran-Armitage test for trend. DMRs p value < 2.2 x 10^-16^, DMGs p value = 1.057 10^-11^, and cumulative DMR size p value < 2.2 x 10^-16^.

A differential DNA-methylation pattern was already observed in blood sample from MOH patients(*18*) and the activity of 5-HT_1_ receptors was found to be paralleled by oscillations in DNA methylation(*31, 32*). Using an in vitro chronic stimulation paradigm (sumatriptan 10 µM), we evaluated time-dependent (6, 12, 24, 48 h) alterations in DNA methylation patterns in human Schwann cells to examine the role of sumatriptan-related epigenetic regulation in these cells. Nanopore long-reads sequencing revealed a time-dependent relationship between sumatriptan exposure and differential DNA methylation in human Schwann cells, compared to vehicle-treated controls. The number of differentially methylated regions (DMRs) progressively increased with treatment duration, from 128 at 6 h to 226 at 24 h and 315 at 48 h, with a corresponding expansion in cumulative DMR size (∼219 kb, 595 kb, and 1054 kb, respectively) (Fig. 3d). These findings indicate that sumatriptan induces a progressive accumulation of epigenetic alterations over time. The Δβ distribution of DMRs was consistent across time points, with a slight increase in hypermethylation at 48 h (Fig. 3e), and an overall balanced representation of hypo- and hypermethylated regions.

To assess the genomic context, DMRs were annotated to genes, revealing 134, 250, and 337 differentially methylated genes (DMGs) at 6 h, 24 h, and 48 h, respectively (including 72, 154, and 202 protein-coding genes). Most DMRs are localized within gene bodies (internal exons and introns), with fewer mapped to the 5′ regulatory regions (promoter and/or first exon) and very few to the 3′ UTR. This pattern did not change over time or between genes with low and high levels of methylation (Fig. 3f). Genes that were both differentially methylated and differentially expressed (DM-DEGs) increased over time, from none at 6 hours to 56 at 24 hours and 85 at 48 hours, according to the integration of transcriptome and methylation data. Notably, among these were 6 and 14 transcription factor genes (at 24 h and 48 h, respectively). Network analysis using RegNetwork revealed that these transcription factors collectively regulate 324 DEGs at 24 h and 186 DEGs at 48 h, highlighting a robust regulatory cascade triggered by sumatriptan-induced DNA methylation. Schwann cell/neuronal paracrine signaling is a key component of various physiological and pathological processes, including axonal regeneration, chronic demyelination, and pain(*33*). Among differentially expressed genes (DEGs) in sumatriptan-stimulated human Schwann cells, we used the ProteINSIDE tool (www.proteinside.org) to identify those involved in intercellular signaling. The DEGs in Schwann cells after sumatriptan stimulation (48 h) associated with the Golgi/ER-dependent secretory pathway included transforming growth factor beta 3 (TGF-β3), interleukin (IL-)32, and IL-15 (Fig. 3g-i). RNAseq data were confirmed by RT-qPCR (Fig. 3j). To investigate the role of TGF-β3 and IL-15 in sumatriptan-dependent MOH in mice, we used monoclonal antibody (mAb) *in vivo* neutralization (IL-32 was excluded because it is not expressed in rodents)(*34*). B6 mice treated with mAb-TGF-β3 but not mAb-IL-15 or IgG were protected from PMA and facial grimacing induced by sumatriptan (Fig. 3k). RT-qPCR and ELISA assays in trigeminal nerve homogenates confirmed the increase of TGF-β3 and IL-15 in MOH mice compared to Control mice (Fig. 3l,m). Pharmacological inhibition of the TGF-β receptors (TGFBRs) with the selective inhibitor LY2109761 reduced PMA and facial grimacing induced by sumatriptan, confirming the role of TGF-β3/TGF-βR signaling in sumatriptan-dependent MOH in mice (Fig. 3n).

### Intronic hypermethylation of BETAGLYCAN (TGFB3 receptor) drives autocrine TGFB3 upregulation

Despite the correlation between epigenomic and transcriptomic modulation in human Schwann cells stimulated with sumatriptan, there were no differentially methylated regions of the *TGFB3* gene that could explain its overexpression. However, the TGF-β3 receptor gene (*TGFBR3*, also known as *BETAGLYCAN*, will be used throughout this text to prevent any confusion with *TGF-β3*) was also overexpressed (confirmed by RT-qPCR Fig. 4a) and was correlated with hypermethylation (Δβ = 0.206) of 19 consecutive CpGs located on the second intron of the *BETAGLYCAN* gene at 48 h compared to vehicle treated cells (Fig. 4b). Moreover, enrichment analysis performed on DEGs regulated by DM-DE-TFs at both 24 and 48 h revealed statistically significant enrichment of genes related to TGF-β signaling (Fig. 4c), indicating that the activation of this signaling pathway is strongly dependent on the transcriptional cascade induced by differential methylation following sumatriptan treatment(*35*). Non-canonical intracellular pathway activated by the TGFBRs include the activation of transcription factors *via* ERK1, p38, or JNK1, which can induce the expression of TGF-β(*36, 37*). β-glycan has been reported to act as a co-receptor of TGF-βR1 and TGF-βR2 augmenting signaling through the canonical and non-canonical TGF-β signaling pathways(*35*). We therefore hypothesized that the chronic sumatriptan stimulation promoted overexpression of β-glycan following intronic hypermethylation inducing increased of non-canonical TGF-βR intracellular signaling, leading to the upregulation of TGF-β3 expression and generating a feed-forward mechanism (Fig. 4d). This is consistent with the enrichment analysis, which shows a more general TGF-β pathway enrichment at 24 hours and increased involvement of SMAD and TGF-βR associated signaling at 48 h (Fig. 4c).To test the correlation between the increase in β-glycan and TGF-β3, we stimulated human Schwann cells with sumatriptan (48 h) and the TGF-βRs inhibitor or vehicle. RT-qPCR and ELISA assay data showed that the overexpression of TGF-β3 was prevented in Schwann cells treated with the TGF-βRs inhibitor LY2109761(Fig. 4e,f). However, β-glycan overexpression was independent from TGF-βRs inhibition but was prevented by the pretreatment with the DNA methyltransferase (Dnmt) inhibitor 5-aza-2′-deoxycytidine (5-AZA-CdR) (Fig. 4g), confirming the temporal sequence of events (Fig. 4d). To determine whether hypermethylation of the second intron of *Betaglycan* mediates PMA and facial grimacing induced by sumatriptan infusion, B6 MOH mice were treated with the DNMT inhibitor 5-AZA-CdR. Repeated administration of 5-AZA-CdR, reduced PMA and facial grimacing in B6 sumatriptan-induced MOH mice (Fig. 4h). This data together indicates that the hypermethylation of the intronic sequence of the *Betaglycan* gene and the subsequent overexpression is essential to start the sumatriptan pain pathway correlated to MOH.

**Figure 4.**
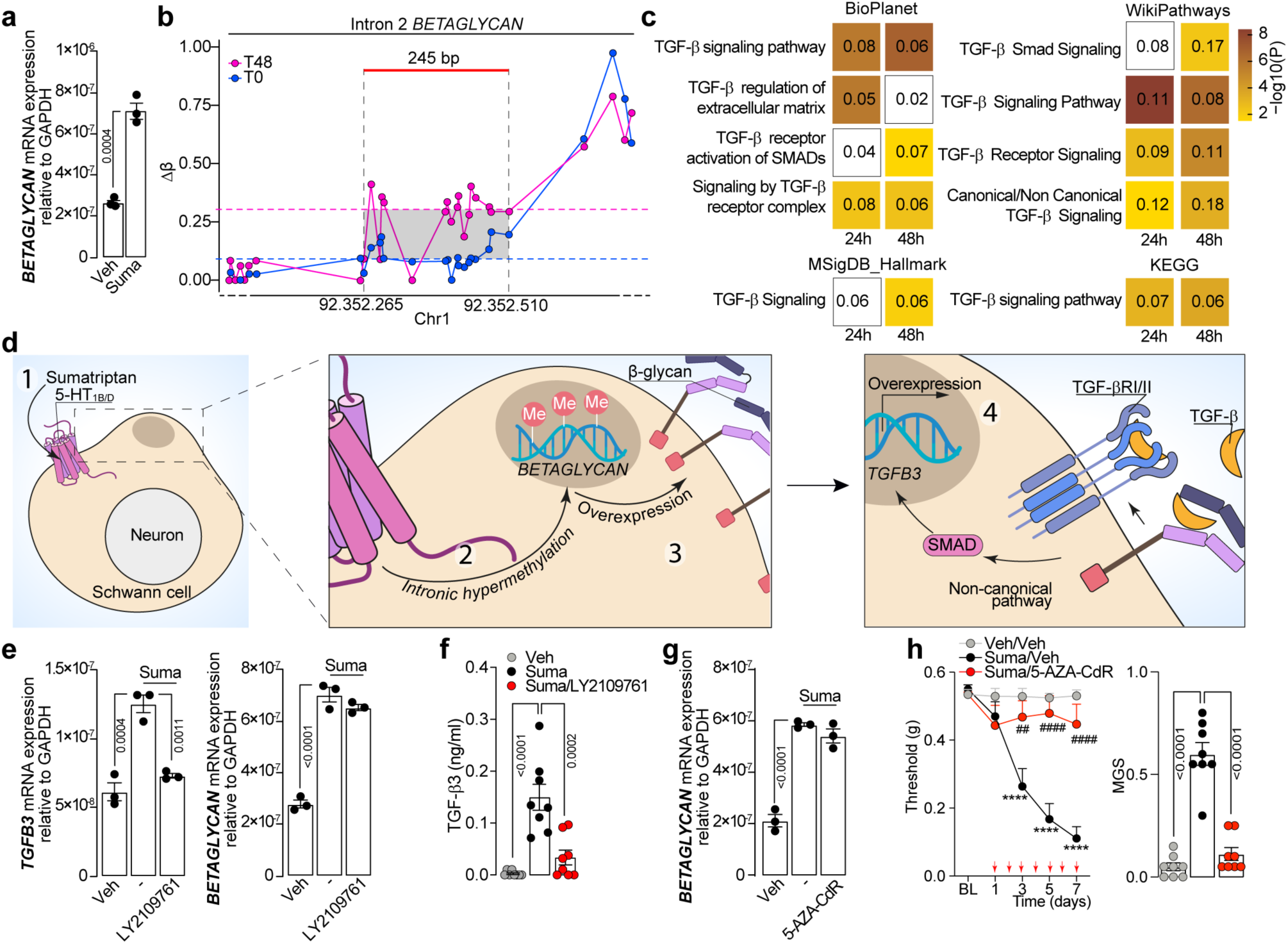
Betaglycan epigenetic regulation drives TGF-β3 signaling in Schwann cells. **(a)** RT-qPCR for *BETAGLYCAN* mRNA and **(b)** methylation changes distribution (Δβ) for CpG sites in the *BETAGLYCAN* intron 2 in human Schwann cells (hSCs) stimulated with sumatriptan (10 μM) (Suma). **(c)** Enrichment analysis on TGF-β pathways in hSCs treated for 24h and 48h with Suma. Numbers indicated the ratio of genes differentially expressed. **(d)** Schematic representation illustrating the sequence of events induced by chronic stimulation: (1) Suma activates SCs 5-HT_1B/D_ receptors, (2) promoting *BETAGLYCAN* intronic hypermethylation, (3) *BETAGLYCAN* over-expression, (4) BETAGLYCAN enhanced non-canonical TGF-βR intracellular signaling and a *TGFΒ3* upregulation establishing a feed-forward mechanism. **(e)** RT-qPCR for *TGFB3* and *BETAGLYCAN* mRNA and **(f)** ELISA assay for TGF-β3 in hSCs stimulated with Suma or vehicle (Veh) and pretreated with TGF-βRs inhibitor LY2109761 (10 μM) or Veh. **(g)** RT-qPCR for *BETAGLYCAN* mRNA in hSC stimulated with Suma or Veh and pretreated with 5-Aza-2′-deossicitidina (5-AZA-CdR, 1 μM) or Veh. **(h)** Periorbital mechanical allodynia and facial grimacing (mouse grimace scale, MGS) in C57BL/6J mice after repeated (once daily over 7 consecutives days) administration of suma (0.6 mg/kg), 5-AZA-CdR (4 mg/kg, i.p.) or Veh. Red arrows indicate Suma and 5-AZA-CdR administration. hSCs are stimulated with Suma in all the experiments for 48h. Data are mean ± s.e.m. (n=8 mice/group). Student’s t-test, 1-way and 2-way ANOVA, Bonferroni correction. ****p<0.0001 vs Veh/Veh, ^##^p<0.01, ^####^p<0.0001 vs Suma/Veh.

### Schwann cell-specific Dnmt, Betaglycan, or Tgfb3 silencing prevents sumatriptan-induced MOH

Next, we investigated whether the feed-forward mechanism identified *in vitro* was confirmed in Schwann cells *in vivo* in mice. To prevent the hypermethylation of the *Betaglycan* intron induced by repeated sumatriptan administration, we used a Cre-dependent AAVrh10 to selectively silence DNMT3A and DNMT3B expression (AAV-*Dnmt3a-Dnmt3b*) in Schwann cells. *Plp^Cre^* MOH mice infected with AAV-*Dnmt3a-Dnmt3b* (*Plp-Dnmt3a-Dnmt3b*) showed reduced PMA and non-evoked pain responses compared to Control or *Plp-Scrambled* mice (Fig. 5a,b). Furthermore, RT-qPCR in trigeminal nerve homogenates showed no overexpression of *Betaglycan* and *Tgfb3* in *Plp-Dnmt3a-Dnmt3b* mice compared to Control or *Plp-Scrambled* mice (Fig. 5c). DNMT silencing was confirmed by immunohistochemistry (Fig. S2a-d). These data indicate that the hypermethylation of the second intron of the *Betaglycan* in Schwann cells in mice after repeated sumatriptan administration was the initial step in the activation of a feed-forward mechanism that led to the development of PMA and spontaneous pain in mice.

**Figure 5.**
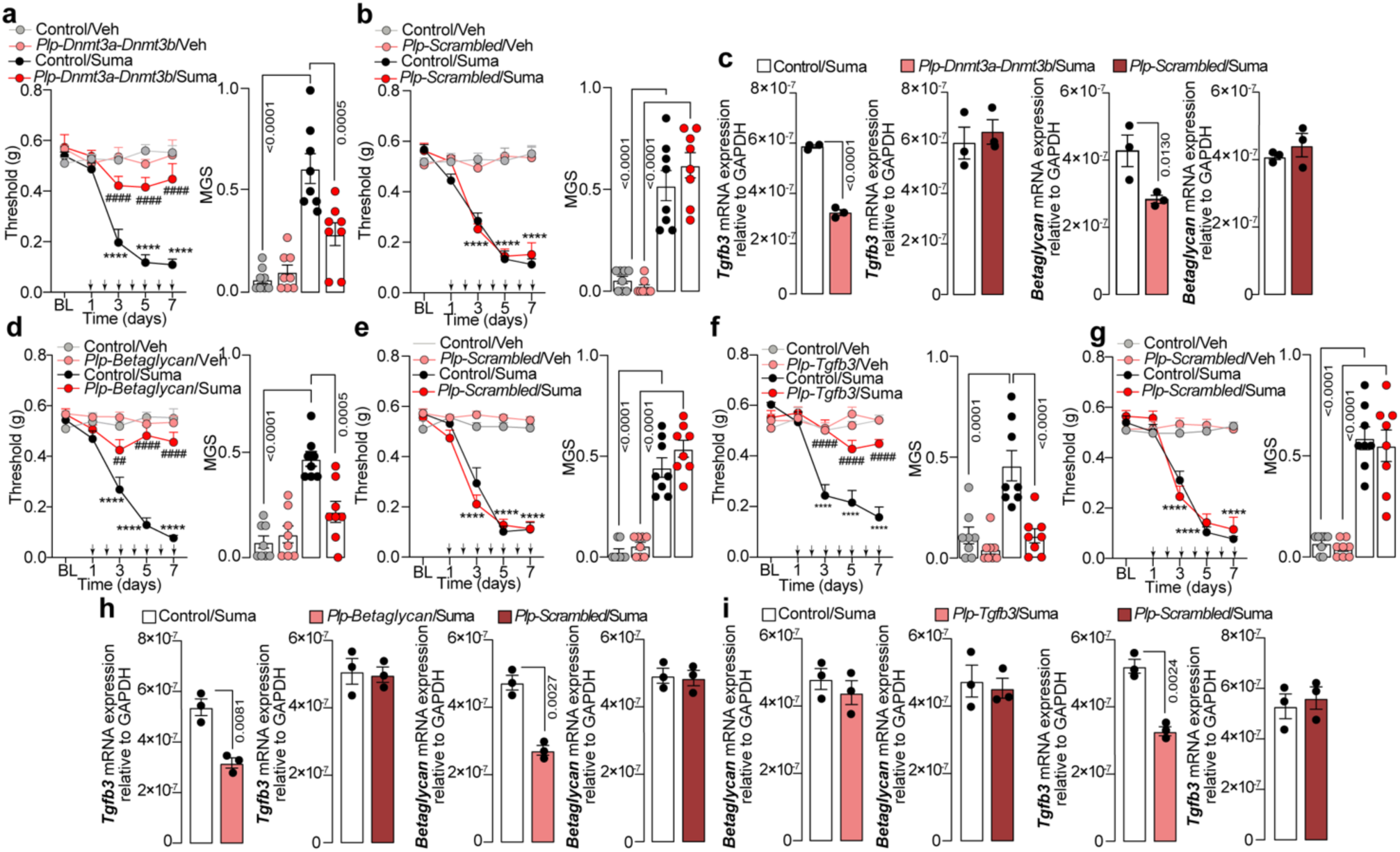
Schwann cell β-glycan/TGF-β3 signaling regulate periorbital mechanical allodynia and grimace behavior in MOH mice. Time dependent periorbital mechanical allodynia (PMA) and facial grimacing (mouse grimace scale, MGS) in Control, **(a)** *Plp-Dnmt3a-Dnmt3b,* **(d)** *Plp-Betaglycan,* **(f)** *Plp-Tgfb3* and **(b,e,g)** *Plp-Scrambled* mice during repeated administration of sumatriptan (Suma, 0.6 mg/kg once daily over 7 consecutive days) or vehicle (Veh). RT-qPCR for *Tgfb3* and *Betaglycan* in Control, *Plp-Scrambled* and **(c)** *Plp-Dnmt3a-Dnmt3b,* **(h)** *Plp-Betaglycan* and **(i)** *Plp-Tgfb3* mice treated with Suma. Black arrows indicate Suma administration. Data are mean ± s.e.m. (n=8 mice/group). Student’s t-test, 1-way and 2-way ANOVA, Bonferroni correction. ****p<0.0001 vs Control/Veh, ^##^p<0.01, ^####^p<0.0001 vs *Plp-Dnmt3a-Dnmt3b*/Veh, *Plp-Betaglycan*/Veh or *Plp-Tgfb3*/Veh.

To substantiate these observations, we used the AAV approach for the Schwann cell selective silencing of the β-glycan and TGF-β3. *Plp-Betaglycan* and *Plp-Tgfb3* MOH mice showed reduced PMA and non-evoked pain responses compared to Control or *Scrambled* mice (Fig. 5d-g). In *Plp-Betaglycan* MOH mice we confirmed a reduction in *Betaglycan* and *Tgfb3* mRNA expression in trigeminal tissues homogenates compared to Control and *Scrambled* mice (Fig. 5h). However, in *Plp-Tgfb3* MOH mice, the *Tgfb3* but not the *Betaglycan* mRNA was downregulated (Fig. 5i). TGF-β3 and β-glycan silencing was confirmed by immunohistochemistry and RNA scope (Fig. S2e-h). All together these findings confirmed that the development of sumatriptan induced MOH is highly regulated by multiple sequential steps that started with the overexpression of β-glycan induced by epigenetic modulation and culminate with TGF-β3 upregulation.

### Schwann cell-derived TGF-β3 activates nociceptors *via* paracrine signaling

Schwann cells interact with neurons *via* paracrine signaling, releasing various mediators that affect neuronal activity and pain sensitivity including TNF-𝛼, NGF, BDNF and hydrogen peroxide(*38–41*). In addition, the excess of TGF-β in pancreas was associated with sensory neuron hyperexcitability, SMAD3 activation, and increased nociception(*42*).

To identify top differentially expressed genes in nociceptive neurons shared across distinct chronic pain conditions in B6 mice, we first analyzed the iPAIN Atlas available on the CELLxGENE browser. This dataset allowed us to pinpoint a conserved nociceptive transcriptional signature (*Atf3, Onecut2, Sox11, Fols2, Klf6, Arid5a,* and *Atf4*) (Fig. 6a)(*43, 44*) Subsequently, to dissect *in vitro* the functional interplay between sumatriptan-stimulated primary mouse Schwann cells and nociceptors, we employed two-chamber co-culture devices connected by a microfluidic system (Fig. 6b). Culture medium of primary mouse Schwann cells stimulated for 48 hours with sumatriptan was able to induce pain-associated genes upregulation in mouse primary sensory neurons compared to vehicle (Fig.6c). Pre-treatment of Schwann cells with the 5-HT_1B/D_ inhibitor GR125743, the TGF-βRs inhibitor LY2109761, the TGF-β3 neutralizing antibody mAb-TGF-β3 but not IgG, prevented the pain-associated genes dysregulation in primary sensory neurons (Fig.6c). Stimulation of DRG neurons with sumatriptan excludes the direct effect in pain associated genes upregulation (Fig.6c). This data confirmed that the chronic *in vitro* stimulation of Schwann cells induced, through a paracrine stimulation, the molecular pain signature activation in primary sensory neuron.

**Figure 6.**
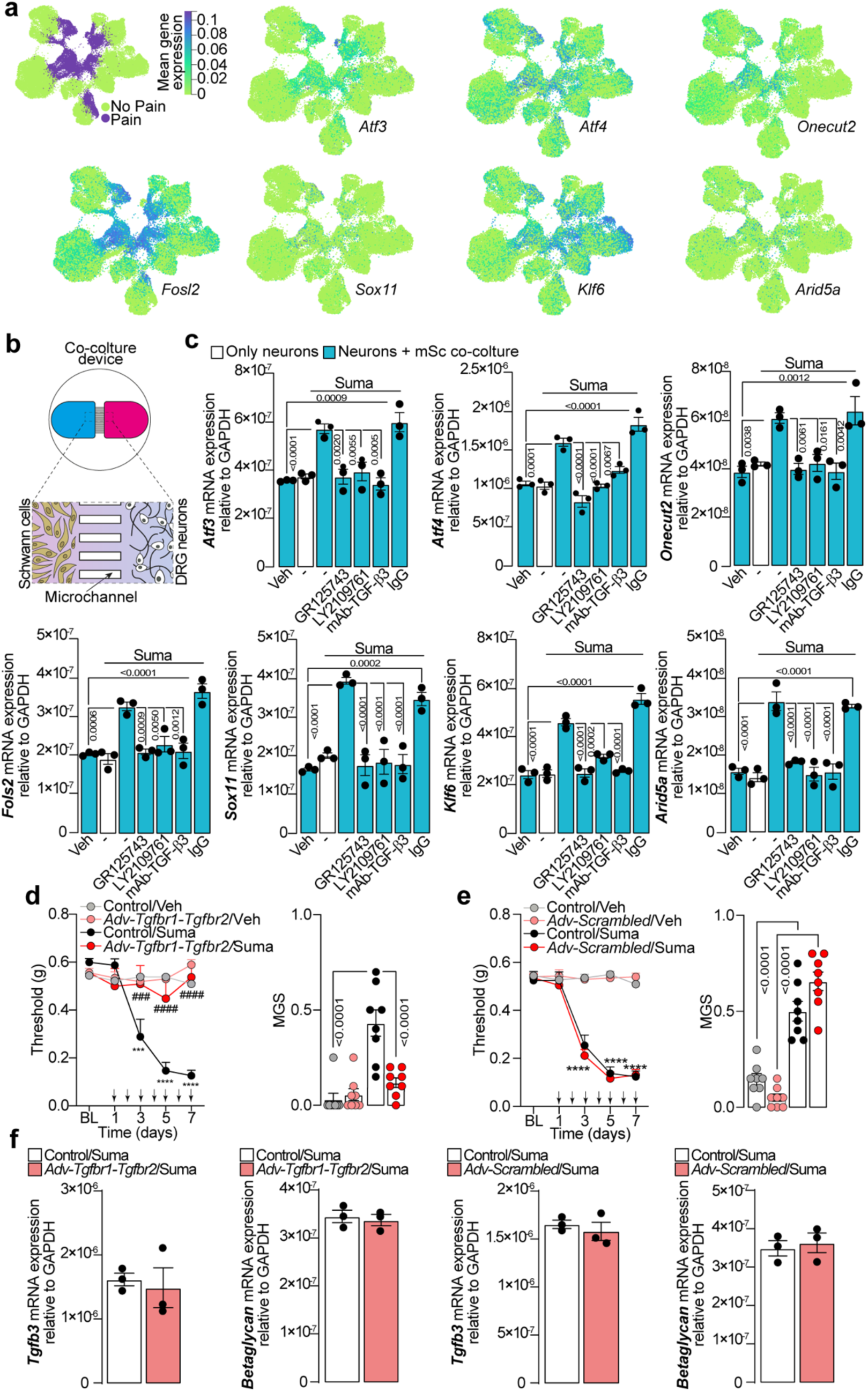
Schwann cell-neuron crosstalk via TGF-β signaling promotes pain in MOH mice. **(a**) UMAP plot of the nociceptive lineage cells highlighted by the pain state identified by aggregation of Leiden clusters. **(b)** Schematic representation of dorsal root ganglia (DRG) neurons and mouse Schwann cells (mSCs) in co-culture device. **(c)** RT-qPCR for *Atf3, Atf4, Onecut2, Fols2, Sox11, Klf6 and Arid5a* in DRG neurons alone or co-cultured with mSCs and stimulated with sumatriptan (Suma) or vehicle (Veh) and pretreated with GR125743 (1 μM), LY2109761 (10 μM), mAb-TGF-β3 (5 μg/ml), IgG (5 μg/ml) or Veh. Time dependent periorbital mechanical allodynia (PMA) and facial grimacing (mouse grimace scale, MGS) in Control, and **(d)** *Adv-Tgfbr1-Tgfbr2 and* **(e)** *Adv-Scrambled* during repeated administration of Suma (0.6 mg/kg once daily over 7 consecutive days) or vehicle (Veh). **(f)** RT-qPCR for *Tgfb3 and Betaglycan* in *Control, Adv-Tgfbr1-Tgfbr2* and *Adv-Scrambled* mice treated with Suma or Veh. Black arrows indicate Suma administration Data are mean ± s.e.m. (n=8 mice/group). Student’s t-test, 1-way and 2-way ANOVA, Bonferroni correction. ***p<0.001, ****p<0.0001 vs Veh/Veh; ^###^p<0.001, ^####^p<0.0001 vs Control/Suma

To assess the nociceptors activation by Schwann cells released TGF-β3, *in vivo*, we used a Cre-dependent bicistronic AAV-PHP.S with *loxP*-flanked shRNA for sensory neurons selective TGF-β receptor 1 and TGF-β receptor 2 (TGF-βR1 and TGF-βR2) silencing (shRNA- *Tgfbr1-Tgfbr2*) and a *Adv^Cre^* driver that functions as a lineage tracer to express shRNA-*Tgfbr1-Tgfbr2* only in sensory neurons of *Adv^Cre^* mice. TGF-βR1 and TGF-βR2 silencing was confirmed by immunohistochemistry (Fig. S3a-d). In *Adv-Tgfbr1-Tgfbr2* mice PMA and facial grimacing induced by sumatriptan infusion was reduced respect to Control mice or *Adv-Scrambled* mice (Fig. 6d,e), however the *Tgfb3* and the *Betaglycan* mRNA was not down-regulated (Fig. 6f). Together, our *in vivo*, *in vitro* and *ex vivo* experiments showed that after the feed forward mechanism activated in SCs the sensory neurons TGF-BRs are the last target of the sumatriptan induced MOH pain pathway.

### Triptans but not NSAID-induced MOH is mediated by Schwann cell 5-HT₁_B/D_-TGFB3 signaling

Since the approval of sumatriptan in 1991, other triptans(*45*), including naratriptan and eletriptan, have been approved for acute migraine treatment(*46–48*). Although their pharmacodynamic profiles are slightly different, they all share agonism for 5-HT_1B/D_ receptors, and despite differences in their formulations and pharmacokinetic properties, these medications have retained the ability to induce MOH(*29, 49*).

We reasoned that sustained activation of Schwann cell 5-HT_1B/D_ receptors could represent a common trigger among triptans in the induction of MOH. Therefore, we administered naratriptan and eletriptan to B6 mice evaluating the development of PMA and facial grimacing (Fig. 7a,b). We then tested whether the pain pathway initiated by the overexpression of β-glycan and TGF-β3 in Schwann cells and culminating in the activation of TGF-β receptors in primary sensory neurons, was also responsible for naratriptan- and eletriptan-induced MOH. We observed that the stimulation with naratriptan and eletriptan induced an upregulation of β-GLYCAN and TGF-β3 in Schwann cells compared to vehicle (Fig. 7c). Consistently, treatment with naratriptan and eletriptan resulted in increased TGF-β3 levels in trigeminal nerve homogenates (Fig.7d). Moreover, repeated administration of naratriptan and eletriptan-induced PMA and facial grimacing was markedly reduced in *Plp-Htr1b-Htr1d*, *Plp-Tgfb3, Plp*-*Betaglycan*, and *Adv-Tgfbr1-Tgfbr2* mice (Fig. 7e-h).

**Figure 7.**
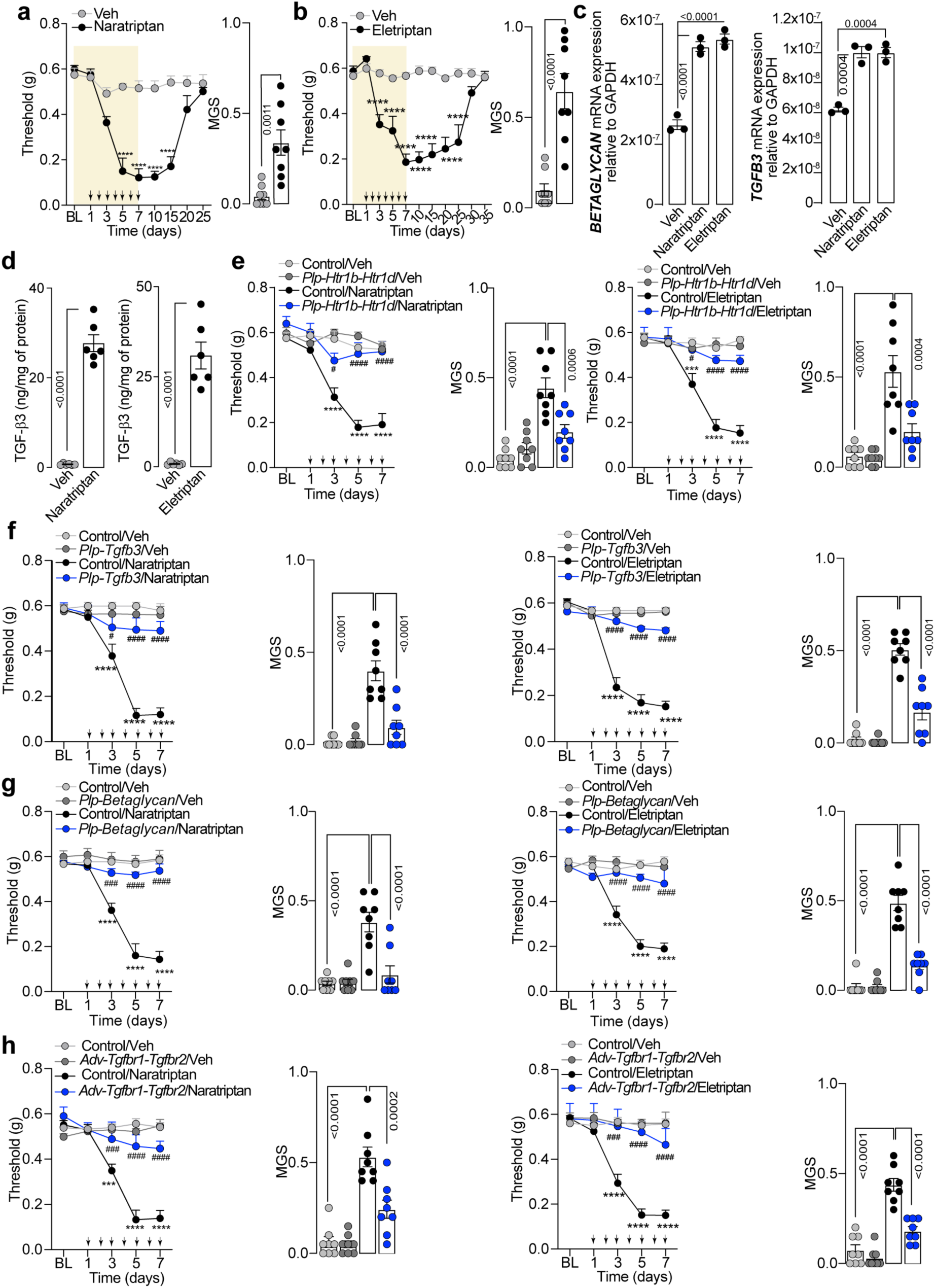
Naratriptan and eletriptan induced periorbital mechanical allodynia and grimace behavior is mediated by Schwann cell 5-HT_1B_/_1D_ -TGF-β3 signaling (a,b) Time dependent periorbital mechanical allodynia (PMA) and facial grimacing (mouse grimace scale, MGS) in C57BL/6J (B6) mice during/after repeated (once daily over 7 consecutives days) administration of naratriptan (0.2 mg/kg i.p.), eletriptan (1 mg/kg, i.p.) or vehicle (Veh). **(c)** RT-qPCR for *BETAGLYCAN and TGFB3* in human Schwann cells (hSCs) stimulated with naratriptan, eletriptan or Veh. **(d)** ELISA assay for TGF-β3 in trigeminal nerve homogenates from naratriptan, eletriptan and Veh treated B6 mice. PMA and facial grimacing induced by repeated naratriptan, eletriptan or Veh in Control and **(e)** *Plp-Htr1b-Htr1d,* **(f)** *Plp-Tgfb3,* **(g)** *Plp-Betaglycan*, **(h)** *Adv-Tgfbr1-Tgfbr2* mice. Black arrows indicate drugs administration. Data are mean ± s.e.m. (n=8 mice/group). Student’s t test, 1-way and 2-way ANOVA, Bonferroni correction. ***p<0.001, ****p<0.0001 vs Veh/Veh; ^#^p<0.05, ^###^p<0.001, ^####^p<0.0001 vs Control/naratriptan/eletriptan.

Although triptans are the most widely recognized medications associated with medication-overuse headache, they are not the only drugs capable of triggering MOH. Prolonged use of common analgesics, including non-steroidal anti-inflammatory drugs (NSAIDs) such as naproxen, diclofenac, and indomethacin for ≥15 days per month, can also lead to rebound headaches. Here, we tested if the mechanism underlying MOH in our sumatriptan-overuse model was shared by other MOH-inducing agents with different pharmacological targets. To address this hypothesis, we used a mouse model of indomethacin-induced MOH. Repeated administration of indomethacin produced a time-dependent increase in PMA and facial grimacing in B6 mice (Fig. 8a), but this response was unchanged in *Plp*-*Htr1b-Htr1d*, *Plp*-*Betaglycan*, *Plp*-*Tgfb3* and *Adv*-*Tgfbr1-Tgfbr2* mice (Fig. 8b-e). These findings showed that the role of Schwann cell 5-HT_1B/D_ receptors is critical for triptan-induced MOH through ligand-receptor interaction and is not a universal mechanism for all MOH-inducing drugs.

**Figure 8.**
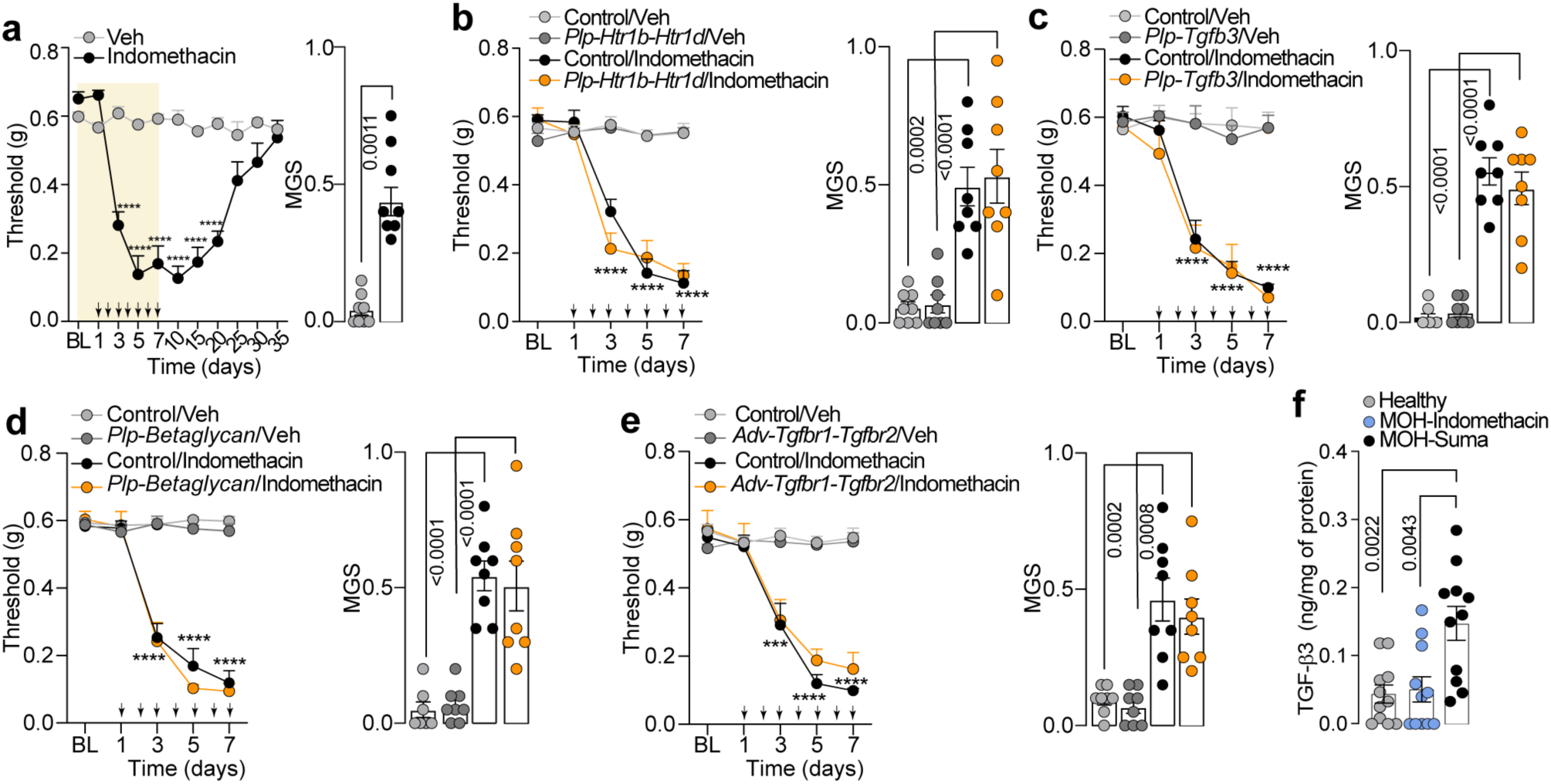
Indomethacin dependent periorbital mechanical allodynia and grimace behavior are not mediated by Schwann cell 5-HT_1B_/_1D_ -TGF-β3 signaling. **(a)** Time dependent periorbital mechanical allodynia (PMA) and facial grimacing (mouse grimace scale, MGS) in C57BL/6J (B6) mice during repeated (once daily over 7 consecutive days) administration of indomethacin (0.2 mg/kg i.g.) or vehicle (Veh). **(b-e)** PMA and facial grimacing induced by repeated indomethacin or Veh in *Plp-Htr1b-Htr1d, Plp-Tgfb3, Plp-Betaglycan*, *Adv-Tgfbr1-Tgfbr2* or Control mice. **(f)** TGF-β3 plasma levels in patients with suma-induced MOH compared with both control subjects and patients with indomethacin-induced MOH. Black arrows indicate drugs administration. Data are mean ± s.e.m. (n=8 mice/group). Student’s t test, 1-way and 2-way ANOVA, Bonferroni correction. ***p<0.001, ****p<0.0001 vs Veh/Veh.

Finally, we evaluated human relevance by analyzing serum samples from patients with MOH due to triptan or indomethacin overuse. ELISA assays confirmed that patients with triptan-induced MOH displayed significantly higher circulating TGF-β levels compared with both control subjects and patients with indomethacin-induced MOH (Fig. 8f). Taken together, these data support a key and specific role of Schwann cell-derived TGF-β as a mediator of triptan-induced MOH, distinguishing it from MOH triggered by analgesics with other pharmacological targets.

## Discussion

Here, we provide evidence that, in the present mouse model of triptan-related medication-overuse headache (MOH), triptans induce a time-dependent hypermethylation of the intronic sequence of the Schwann cell β-glycan gene. This epigenetic modification activates a proalgesic signaling pathway involved in neuronal pain transmission. We reveal that Schwann cells express the 5-HT₁_B/D_ receptors, which acutely mediate the antimigraine effects of sumatriptan by inhibiting CGRP-induced endosomal cAMP accumulation. However, chronic activation of these receptors leads to epigenetic/transcriptomic dysregulations in Schwann cells associated with the development of periorbital mechanical allodynia. Another unexpected finding of the present study highlights a dual role for Schwann cell-derived TGF-β3. Autocrinally, TGF-β3 exploits the overexpression of β-glycan to enhance its own transcription, while paracrinally, activates primary sensory neuron TGF-β receptors, contributing to the transmission of nociceptive signals.

The cAMP signaling pathway is critically regulated by the modulation of GPCRs and plays a central role in a wide range of cellular and physiological processes, including immune function(*50, 51*), metabolism(*52, 53*), and pain(*54, 55*). Our recent findings indicate that increases in intracellular cAMP levels within Schwann cells, compartmentalized in distinct subcellular nanodomains, can drive the emergence of different pain phenotypes in murine models. Specifically, EP2 receptor dependent cAMP elevation at the plasma membrane promotes TRPA1 sensitization through a transphosphorylation mechanism, thereby facilitating inflammatory pain(*56*). Conversely, the internalization of the CALCRL/RAMP1 receptor complex leads to an endosomal localized cAMP increase, which in turn activates endothelial nitric oxide synthase (eNOS), resulting in NO production and pain signaling(*19*). Here, we observed that 5-HT₁_B/D_ receptors internalize in endosomal nanodomains after sumatriptan stimulation, suggesting a spatially confined mechanism that counteracts CGRP-induced cAMP elevation. Supporting this hypothesis, the abortive effect of sumatriptan in CGRP-induced periorbital mechanical allodynia was absent in mice with 5-HT₁_B/D_ selective silencing in Schwann cells.

The cAMP-mediated phenotypic alterations are not restricted to the canonical activation of PKA and subsequent phosphorylation of effector protein. Increasing evidence demonstrates that cAMP regulates gene expression through epigenetic mechanisms(*57*). Such mechanisms have been implicated in several biological contexts, including alterations in DNA methylation in oocytes from obese mothers(*58*), early differentiation of pluripotent stem cells(*59*), and macrophage maturation driven by CSF1R signaling(*60*). In the peripheral nervous system, cAMP is also essential for Schwann cell differentiation(*61*) regulating transcriptional activity by enhancing DNA hydroxymethylation(*62*). Epigenetic regulation is now recognized as a key contributor to both the development and persistence of pain states. Inflammatory and neuropathic pain conditions are associated with HDAC-mediated histone deacetylation and repression of *Gad2* expression(*63*), differential DNA methylation of the *TRPA1* promoter has been correlated with inter-individual variability in pain sensitivity(*64*), while nerve injury-induced methylation of the *Kcna2* promoter enhances excitability in DRG neurons(*65*). In the spinal dorsal horn, HDAC4 was shown to regulate *OAT1*, contributing to nociceptive hypersensitivity under inflammatory conditions(*66*). Furthermore, pharmacological inhibition of HDAC1/2 alleviates oxycodone withdrawal-associated mechanical allodynia(*67*), and HDAC9 facilitates chronic neuropathic pain through upregulation of miR-203-3p in models of nerve injury(*68*).Gene-wide DNA methylation studies(*69, 70*) and investigations in both rodents and humans revealed that epigenetic modifications in key genes, including *Rimbp2* and *Grip*(*71*), as well as *Crcp*, *Calcrl*, and *Calca*(*72–74*), are associated with migraine-related pain. To date, only two studies have investigated DNA methylation in the context of MOH, highlighting associations between the methylation status of *HDAC4*(*24*)*, CORIN*, *CCKBR* and *CLDN9*(*18*) and headache chronification, based on analyses of peripheral blood cells from patients.

In our study, prolonged exposure of human Schwann cells to sumatriptan induced epigenetic modifications of *Betaglycan*, leading to transcriptional upregulation. Under basal conditions, Schwann cell-derived TGF-β3 exploits the increased expression of *β-glycan* to further enhance its own transcription *via* a positive feedback loop, thereby sustaining the proalgesic signaling cascade. A limitation of this study is that we did not assess DNA methylation of the *β-glycan* gene directly in Schwann cells in the peripheral nerve tissue of MOH mice. Nevertheless, selective silencing of Schwann cell DNMTs significantly attenuated mechanical allodynia in MOH mice, supporting the critical role of Schwann cell-specific epigenetic modulation in the pathogenesis of MOH.

The role of TGF-β in pain modulation appears to be context-dependent, displaying both protective and pro-nociceptive effects in different conditions. For instance, in rat models of intervertebral disc degeneration(*75*), nerve injury(*76*), and chronic constriction injury(*77*), exogenous TGF-β administration has been shown to prevent the development of pain. Conversely, in rat models of chronic pancreatitis(*42, 75, 78*) and bone cancer(*79*), TGF-β signaling contributes to the facilitation of nociceptive transmission. In this context, TGF-β3 not only sustains its own transcription through an autocrine mechanism but also activates primary sensory neurons in a paracrine manner, contributing to nociceptive signaling. The most plausible explanation for the pro-nociceptive role of TGF-β observed in the MOH mouse model, contrasting with its analgesic effects reported in other chronic pain conditions, may lie in the highly localized Schwann cell-neuron paracrine interaction. In other models, where TGF-β levels are systemically elevated, its effects may be mediated indirectly through the modulation of non-neuronal cell populations, rather than through direct action on sensory neurons. Future studies should determine whether Schwann cell-specific epigenetic remodeling also occurs in patients with triptan-induced MOH, enabling the identification of predictive biomarkers and stratification strategies for personalized therapies. In addition, targeting Schwann cell-neuron paracrine signaling pathways may represent a novel therapeutic approach to preserve the acute efficacy of triptans while preventing the transition to MOH.

## Material and Methods

### Study design

The study combined mechanistic preclinical experiments with translational validation in human subjects to investigate the role of Schwann cell 5-HT1B/D receptor signaling in MOH. In mice, we employed Schwann cell-selective genetic silencing of 5-HT1B/D receptors to dissect cell-specific contributions to both the acute and chronic effects of triptans. Acute paradigms assessed the ability of triptan administration to modulate CGRP-induced periorbital mechanical allodynia. Chronic paradigms modeled MOH through repeated administration of triptans, followed by behavioral assessment of evoked and non-evoked pain responses and molecular analyses of peripheral nerves. To investigate causality within the identified pathway, we performed Schwann cell-selective silencing of DNA methyltransferases, β-glycan (TGFBR3), and TGF-β3, as well as pharmacological and antibody-mediated inhibition of TGF-β signaling. Complementary in vitro studies in primary human and mouse Schwann cells, including co-culture systems with sensory neurons, were used to define epigenetic, transcriptomic, and paracrine mechanisms. DNA methylation profiling and RNA sequencing identified a time-dependent epigenetic reprogramming characterized by intronic hypermethylation-driven β-glycan overexpression and activation of a non-canonical TGF-β3 signaling cascade. To establish specificity, parallel experiments were conducted using multiple triptans and a non-steroidal anti-inflammatory drug (indomethacin). Translational relevance was assessed by measuring circulating TGF-β3 levels in well-characterized patient cohorts with MOH, stratified by medication type. Together, this integrative approach enabled the identification of cellular and molecular mechanisms linking triptan exposure to both therapeutic and pathological outcomes. All work was performed in compliance with the institutional and international and national regulations. Animal experiments were carried out in accordance with European Union (EU) guidelines for animal care procedures and the Italian legislation (DLgs 26/2014) application of the EU Directive 2010/63/EU. The study was approved by the National Committee for the Protection of Animals used for Scientific Purposes of the Italian Ministry of Health (research permits #622/2025-PR). The group size of n=8 mice for behavioral experiments was determined by sample size estimation using G Power [v3.1](*80*) to detect the size effect in a *post-hoc* test with type 1 and 2 error rates of 5% and 20%, respectively. Allocation concealment of mice into the vehicle(s) or treatment groups was performed using a randomization procedure (http://www.randomizer.org/). The assessors were blinded to the identity of the animals (genetic background) or allocation to treatment groups. None of the animals were excluded from the study. Mice were housed in a temperature- and humidity-controlled *vivarium* (12 h dark/light cycle, free access to food and water, 5 animals per cage). At least 1 h before behavioral experiments, mice were acclimatized to the experimental room and behavior was evaluated between 9:00 am and 5:00 pm. Behavioral studies followed Animal Research: Reporting of *In Vivo* Experiments (ARRIVE) guidelines (*81*). Animals were anesthetized with a mixture of ketamine and xylazine (90 mg/kg and 3 mg/kg, respectively, i.p.).

The use of human tissue samples was approved by the local ethics committees of the Careggi University Hospital, Florence (protocol #29447_bio). The use of human plasma samples from patients with MOH due to triptan overuse and from those with MOH caused by indomethacin or healthy controls was approved by the local ethics committees of the IRCCS Mondino Foundation (Number 20170023686). The blood plasma samples derived from thirty-three subjects: triptan-induced MOH [female 9/male 2, median age 48 years (33-63], MOH caused by indomethacin [female 10/male 1, median age 41 years (24–60)] and healthy individuals [female 7/ male 4, median age 39 years (29–57)]. Blood samples were collected within ethylenediamine tetra-acetic acid (EDTA) containing tubes. Considering the exploratory nature of the blood sampling analyses, the sample size was not determined based on statistical considerations but enrolling all consecutive outpatients and healthy controls. Written informed consent was obtained from all participants.

### Materials

All reagents, unless otherwise indicated, were from Merk Life Science S.r.l. Detailed of other materials and suppliers are provided in the specific sections.

### Animals

Male and female mice C57BL/6J (Charles River, RRID: IMSR_JAX:000664) (25-30 g, 6-8 weeks old) were used throughout. Hemizygous B6.Cg-Tg(Plp1-CreERT)3Pop/J mice (Plp-Cre^ERT^, RRID: IMSR_JAX:005975 Jackson Laboratory) expressing a tamoxifen-inducible Cre in Schwann cells (Plp1, proteolipid protein myelin 1)(*41*) were used. The progeny was genotyped using PCR for *Plp-Cre^ERT^.* Mice that were negative for *Plp-Cre^ERT^* (Plp-Cre^-^) were used as control. Some *Plp-Cre^ERT^*^+^ or *Plp-Cre^ERT^*^-^ were treated with intraperitoneal (i.p.) 4-hydroxytamoxifen (4-OHT, 1 mg/100 μL in corn oil, once a day consecutively for 3 days) before the infection with AAV for selective silencing of the different genes in Schwann cells. Hemizygous Advillin-Cre mice (Adv-Cre)(*82*) positive (Adv-Cre^+^) or negative (Adv-Cre^-^, Control) for Cre were used. Adv-Cre^+^ or their respective control were used for the infection with AAV for selective silencing of the different genes in Schwann cells or in primary sensory neurons, respectively.

### Behavioral assays

#### Periorbital mechanical allodynia

Periorbital mechanical allodynia (PMA) was assessed using the up-down paradigm. Briefly, mice were placed in a restraint apparatus designed for the evaluation of periorbital mechanical thresholds(*83*). Mechanical threshold was evaluated in the periorbital region (rostral eye area) before and after treatments. On the test day, following 20 min acclimation, von Frey filaments (0.02–1.0 g) were applied perpendicularly to the skin for ∼5 s. A positive response was defined as face stroking, head withdrawal, or head shaking. Six measurements were collected per mouse or until four consecutive identical responses occurred. The 50% withdrawal threshold was calculated using a δ value of 0.205.

#### Facial grimace score

Spontaneous pain was tested using the Mouse Grimace Scale(*84*). Briefly, each mouse was placed in an individual chamber (9 × 5 × 5 cm) having transparent Plexiglas walls to allow experimental observation. Mice were habituated for 30 min before behavioral testing. Cameras directed at the front of the cubicle recorded 30 min of facial expressions. One clear facial image was taken for every 3 min interval, scrambled and scored blindly for facial grimacing (*84*). The scorers assessed facial expressions such as orbital tightening (closing or narrowing of the eyelid and orbital area), ear position (outward or backwards rotation of the ears), scoring either a 0 (not present), 1 (moderately visible) or 2 (severe) depending on the magnitude of the expression. Nose and cheek bulging, and whisker change were not score because they were difficult to distinguish against the dark fur of the mice.

#### Treatment protocols

Mice received intraperitoneal (i.p. 10 ml/kg) injection of CGRP (0.1 mg/kg) or vehicle (0.9% NaCl). Sumatriptan (0.6 mg/kg, i.p.) or vehicle (0.9% NaCl), GR125743 (0.5 mg/kg, i.p., #HY-121392, DBA) or vehicle (0.9% NaCl) were administered 30 minutes before CGRP. In another set of experiments, sumatriptan (0.6 mg/kg, i.p.), eletriptan (1 mg/kg, i.p.), naratriptan (0.2 mg/kg i.p.) or their vehicle (0.9% NaCl), indomethacin (2 mg/kg, intragastric, i.g.) or vehicle (1% carboxymethylcellulose in H_2_O) were administered once a day from day 0 to day 7. GR125743 (0.5 mg/kg, i.p.), LY2109761 (50 mg/kg, i.g., DBA), 5-Aza-2’-deossicitidina (5-AZA-CdR, 4 mg/kg i.p.), or vehicle (0.9% NaCl), monoclonal antibody anti-TGF-β3 (200 μg/mouse, i.p., #BE0057, Bioxcell) or anti-IL-15 (200 μg/mouse #BE0315, Bioxcell) isotype control (IgG) were administered just before sumatriptan once a day from day 0 to day 7. *Plp-Cre^+^* and *Adv-Cre^+^* and their control mice were infected with an intravenous (i.v., 10 µL, 1×10^12^ vg/ml) injection of different AAV and used 3 weeks after infection. Sciatic nerve and DRGs were harvested for evaluating AAVs infection.

### Plasmid Constructs

#### In vivo study

pAAV[FLEXon]-CMV- (EGFP-5’ miR-30E-BfuAI-ORF-BfuAI-3’-miR-30E)-WPRE obtained from Vectorbuilder was modified adding a Multi Cloning Site coding for AscI-NheI-AflII-BamHI (5’-ggCGCGCCGCTAGCatcatcatCTTAAGGgatcc-3’) between AscI and BamHI. The new plasmid pAAV[FLEXon]-CMV-rev(EGFP-5’-miR-30E-BfuAI-ORF-BfuAI-3’-miR-30E-NheI/AflII)-WPRE was used as backbone for a short hairpin RNA targeting gene of interest or Scrambled shRNA as negative control. shRNAs targeting the gene of interest or Scrambled shRNA were cloned by replacing the ORF with preannealed and phosphorylated oligonucleotides by using BfuAI compatible bases. shRNAs and Scrambled shRNA oligos are annealed as follow: P1/P2 Htr1b, P3/P4 Htr1d, P5/P6 Betaglycan, P7/P8 Tgfb3, P9/P10 Dnmt3a, P11/P12 Dnmt3b, P13/P14 Tgfbr1, P15/P16 Tgfbr2, P17/P18 Scrambled Htr1b, P19/P20 Scrambled Htr1d, P21/P22 Scrambled Betaglycan, P23/P24 Scrambled Tgfb3, P25/P26 Scrambled Dnmt3a, P27/P28 Scrambled Dnmt3b, P29/P30 Scrambled Tgfbr1, P31/P32 Scrambled Tgfbr2. Multicistronic plasmids were obtained by cutting the donor plasmid with AvrII/AflII and the receiving plasmid with NheI/AflII. Using the compatible ends of NheI and AvrII the plasmid pAAV[FLEXon]-CMV-(EGFP-5’-miR-30E-shRNA1-3’-miR-30E-5’-miR-30E-shRNA2-3’-miR-30E-NheI/AflII)-WPRE was obtained. All constructions were confirmed by Sanger sequencing. Primer sequences are listed in Table S1.

#### In vitro study

From pCAG-G-FLAMP2 (#192782, RRID:Addgene_192782, Addgene), G-FLAMP2 was cloned (EcoRI/XbaI) into pEG backbone (P33/P34). pEG G-FLAMP2-Rab5a was generated by amplifying with overlap-PCR G-FLAMP2 (P35/P36) and Rab5a from mCherry-Rab5 (#49201, RRID:Addgene_49201, Addgene) (P37/38) and cloning onto pEG backbone with EcoRI/SalI. Human 5-HT_1B_ cDNA (NP_000854) and Human 5-HT_1D_ cDNA (NP_000855) have been purchased from R&D Systems™ (respectively #RDC0284, #RDC0285). pEGFP 5-HT_1B_-EGFP and pEGFP 5-HT_1D_-EGFP were generated by cloning 5-HT_1B_ (P39/P40) and 5-HT_1D_ (P41/P42) into pEGFP-C1 (Clontech) using EcoRI/KpnI. Using NanoBiT® MCS Starter System (#N2014, Promega), LgBiT and SmBiT fusions proteins were generated. Using a Gibson assembly strategy, pBit2.1-C-5-HT_1B_-SmBIT (P43/P44 for 5-HT_1B_ fragment and P45/P46 for acceptor vector) and pBit2.1-C-5-HT_1D_-SmBIT (P47/P48 for 5-HT_1D_ fragment and P49/P50 for acceptor vector) were cloned. pBit1.1-N-LgBit-Rab5a was cloned with EcoRI/NheI into pBit1.1-N-LgBit (P51/P52). Plasmids were cloned using Gibson Assembly Cloning Kit (#E5510, New England Biolabs) or with restriction enzymes. All constructs were confirmed by Sanger sequencing. Primer sequences are listed in TableS1.

#### AAV generation

Recombinant AAV production, cell lysis, AAV purification and AAV titration were described in Supplementary Material.

#### Cell lines HEK293T cell line

Human kidney epithelial (HEK293T) (#CRL-3216™, RRID:CVCL_0063, ATCC) were cultured in Dulbecco’s Modified Medium (DMEM) supplemented with heat inactivated fetal bovine serum (FBS, 10%), and L-glutamine (2 mM) at 37°C in 5% CO_2_ and 95% O_2_ in a humidified atmosphere.

#### AAVpro293T cell line

AAVpro 293T cells (#632273, Takara), were maintained in DMEM high glucose supplemented with heat inactivated FBS (10%), L-glutamine (4 mM), penicillin/streptomycin (1 mM) and sodium pyruvate (1 mM) at 37 °C in 5% CO_2_ and 95% O_2_. The day before transfection, cells were plated in DMEM supplemented with 2% FBS.

#### Primary human Schwann cells

Commercial human primary Schwann cells (hSCs) (#P10351, Innoprot) were grown and maintained in Schwann cell medium (#P60123, Innoprot) at 37 °C in 5% CO_2_ and 95% O_2_. Cells were discarded and replaced after 12 passages. In a set of experiments hSCs were treated with sumatriptan (10 µM) for 48 hours in FBS- starved medium. In selected experimental groups, sumatriptan-treated cells were simultaneously exposed to LY2109761 (10 µM), 5-AZA-CdR (1 µM), or vehicle (0.001 % DMSO). All compounds were added at the beginning of the treatment and maintained for the entire 48-hour period. In parallel experiments, hSCs were treated with naratriptan (10 µM), eletriptan (10 µM) or vehicle (0.001 % DMSO) for 48 hours under the same serum-starved conditions. At the end of the treatment period, cells and conditioned media were collected for further analyses.

Isolated hSCs were obtained from lingual, sublingual and inferior alveolar nerves biopsies. After resection, tissues kept in sterile saline on ice were washed with sterile PBS and transferred to DMEM/F12(#21331020; Waltham, Massachusetts, USA), containing penicillin (100 U/mL), streptomycin (100 mg/mL) and L-glutamine (2mM). The *epinerium* was removed under a dissecting microscope and nerves were cut into 1 mm segments. Tissues were then incubated for 24 at 37° C with 95% O_2_ and 5% CO_2_ levels in an enzymatic mixture containing hyaluronidase Type I-S (250 U/mL) and collagenase Type I (160 U/mL) in DMEM/F12 medium. After 24 hours, the cell culture-containing medium was passed through an 18-gauge needle and then centrifugated at 1000 g for 5 min at room temperature (RT). The pellet was resuspended in supplemented DMEM/F12 medium containing FBS (10 %) and plated on poly-L-ornithine (0.1 mg/mL) and laminin (1µg/cm^2^) pre-coated 12-well dishes. Fresh culture medium was added after 24 hours and then every three days for 20 days prior to the experiments.

#### Primary mouse Schwann cells

Primary mouse Schwann cells (mSCs) were obtained from sciatic nerves of B6 mice (n=10 mice). Briefly, after removal of the *epineurium*, nerve explants were cut into1 mm segments and dissociated in Hank’s Balanced Salt Solution (HBSS) containing hyaluronidase (0.1%) and collagenase (0.05%) for 2 h, at 37 °C in agitation. After centrifugation (150 x g, 10 min, RT), the pellet was resuspended and cultured in DMEM containing streptomycin (100 mg/ml), fetal calf serum (10%), forskolin (2 μM), L-glutamine (2 mM), penicillin (100 U/ml) and neuregulin (10 nM). Cytosine arabinoside (Ara-C, 10 mM) was added after three days, to remove fibroblasts. Cells were cultured at 37 °C in 5% CO_2_ and 95% O_2_ for 15 days before experiments by replacing the culture medium every 3 days.

#### Primary mouse dorsal root ganglion (DRG) neurons

Mouse DRGs (combined cervical, thoracic, and lumbar) were bilaterally removed from B6 mice (n=8 mice) under a dissection microscope and then digested in HBSS for 35 min at 37 °C containing of collagenase type 1A (2 mg/mL) and papain (1 mg/mL). Ganglia were disrupted through several passages utilizing a series of syringe needles (23–25G). Mouse DRG neurons were centrifuged at 180 x g for 5 min at RT and then resuspended in DMEM/F12, containing: FBS (10%), penicillin (100 U/ mL), streptomycin (0.1 mg/mL), L-glutamine (2 mM) added with nerve growth factor (100 ng/mL), and cytosine-b-D-arabino-furanoside free base (2.5 mM). Cells were maintained at 37 °C in 5% CO_2_ and 95% O_2_ for 3 days before being used.

#### Double co-culture

Co-culture experiments were performed using OMEGAACE devices (#eN-o96-001; eNUVIO Inc.). Every device contained 2 chambers of identical dimensions (chamber #1 and #2) directly connected *via* a series of microfluidic channels (number of microchannels/pair: 70, microchannel length: 640 µm, microchannel width: 10 μm). The chambers were separated by a 250 μm-wide and 500 μm-high channel. To mimic the *in vivo* mechanism, experiments were performed using mouse primary Schwann cells and DRG neurons. Briefly, Schwann cells were plated in chamber #1 and DRG primary sensory neurons in chamber #2. To confirm the role of Schwann cells in one condition the chamber #1 was empty. Cells were seeded at a density of 3 × 10^5^ cells/cm^2^. The direction of fluid flow across the high-resistance microchannels (from Schwann cells to DRG neurons) was determined by asymmetrical volume loading contained within each of the chambers, and the volumes were as follows: 250 μL in chamber #1 and 200 μL in chamber #2. Schwann cells were treated with sumatriptan (30 μM) or vehicle (0.001% DMSO) in FBS- starved medium before being placed in contact with neurons for 48 hours prior to further analysis. In a set of experiments, GR125743 (#HY-121392, DBA, Italy), LY2109761 (10 μM, #HY-12075, DBA), mAb-TGF-β3 (#BE0057 bioxcell, RRID:AB_1107757, 5 μg/ml), IgG (#BE0122, RRID:AB_10951292 5 μg/ml), or vehicle (0.001% DMSO) were added to chamber #1. Cells were incubated at 37 °C in 5% CO_2_ and 95% O_2_ for 24 h. To minimize evaporation from the chambers during incubation steps, ∼500 μL of sterile water was added to the circular track of the evaporation minimizer.

#### cAMP in vitro imaging

To measure cAMP signaling in real time, primary commercial and isolated hSCs and primary mSCs were plated in 96-well poly-L-lysine-coated (8.3 μM) white clear bottom plates (5 ×10^5^ cells/well; #6005181, PerkinElmer) and transfected with GFlamp2-Rab5a (120 ng DNA) with jetOPTIMUS® DNA transfection reagent (#55-250, Polyplus). The fluorescent signal was recorded using a fluorescent microscope (Axio Observer 7; with a fast filter wheel and Digi-4 lens to record excitations and Ultra-fast Sutter Lambda DG4 Xenon excitation source (range 300-700 nM) (Zeiss) with the following settings: ex/em 506/517 nm (filter set: FT 495, ex BP 470/40, em BP 525/50, interval 1 s) HSCs were stimulated with graded concentrations of calcitonin gene related peptide (CGRP, 3 nM-30 µM). The response to CGRP (3 µM) was evaluated in the presence of sumatriptan (100 nM-300 µM) or vehicle (0.001% DMSO). In another set of experiments, cells were incubated with sumatriptan (30 μM) also in combination with GR125743 (1 μM, #HY-121392, DBA). Signals were recorded for approximately 300 s, was calculated for each experiment and the results were expressed as the percentage increase in the ΔF/F0 ratio over baseline normalized to the maximum effect induced by forskolin (10 μM) added at the end of each experiment.

#### Nanobit complementation assay

To evaluate the dynamics of 5-HT_1B_ and 5-HT_1D_ proximity in hSCs, a time-resolve luciferase re-complementation assay was used. hSCs were plated in 96-well poly-L-lysine-coated (8.3 μM) white clear bottom plates (5 ×10^5^ cells/well; #6005181, PerkinElmer) the day before transfection. pBit2.1-C-5-HT_1B_-SmBIT or pBit2.1-C-5-HT1_1D_-SmBIT were co-transfected with pBit1.1-N-LgBit-Rab5a (120 ng DNA in total) with jetOPTIMUS® DNA transfection reagent (#55-250, Polyplus). 48 h after the transfection cells were treated for 10 min with Nano-Glo® Live Cell Reagent, according to manufacture instructions (#N2011, Nano-Glo^®^ Live Cell Assay System, Promega). Then, hSCs were stimulated with sumatriptan (10 µM) or vehicle (0.001% DMSO). The response to 10 µM sumatriptan was evaluated in the presence of 5-HT_1B/D_ inhibitor GR12574335 (10 μM, #HY-121392, DBA) or its vehicle (0.001% DMSO). Luminescence was measured for 6 min using a luminescence plate reader (FlexStation3; Molecular Devices) with SoftMax^®^ Pro7 software (Molecular Devices). The results are expressed as arbitrary units (AU) The ΔF/F0 ratio was calculated for each experiment, and the results were expressed as the area under the curve (AUC).

#### Reverse transcription-quantitative and real time PCR (RT-qPCR)

Tissues were collected from mice and stored in RNA later at −80°C until processing. Total RNA was extracted from hSCs, primary mSCs using the RNeasy Mini Kit (Qiagen SpA), according to the manufacturer’s protocol. RNA from DRGs and sciatic nerves was extracted with RNeasy lipid tissue Mini Kit (Qiagen SpA). The quantity and quality of the RNA were assessed using an assessed spectrophotometrically by measuring the absorbance at 260 and 280 nm with QIAxpert System. RNA was reverse transcribed using SuperScript™ IV VILO™ Master Mix (Invitrogen) following the manufacturer’s protocol. For relative mRNA quantification, RT-PCR was performed using Rotor Gene® Q software (v.2.3.1.49; Qiagen SpA) and KAPA SYBR® FAST. The relative abundance of mRNA transcripts was calculated using the ΔCT method and normalized to glyceraldehyde-3-phosphate dehydrogenase (GAPDH) levels. The sets of primers are reported in Table S2.

#### Nanopore DNA sequencing

Genomic DNA was extracted from hSC after stimulation with sumatriptan (10µM) for 0, 6, 24 and 48 hours. Cells were harvested washed with DBPS twice and lysed using DNeasy Blood and Tissue kit (#69506 Qiagen SpA) according to manufacture instructions. DNA libraries from Schwann cells at each treatment timepoint were prepared using the SQK-LSK109 ligation kit (Oxford Nanopore Technology) and sequenced on a PromethION P2 Solo instrument using FLO-PRO002 r9.4.1 flow cells. Raw sequencing data were basecalled using Guppy (v6.5.7) with the sup_5hmc_5mc_CG model; alignment to the GRCh37 reference genome was performed using the integrated version of minimap2. Read-level 5mC and 5hmC probabilities were extracted from basecalled data using the modkit extract function (v0.1.4). DMRs were identified using PoreMeth2 by comparing methylation probabilities at 6, 24, and 48 hours against vehicle-treated cells, selecting regions with a p value ≤ 0.05 and |𝛥𝛽| ≥ 0.2. Finally, DMRs were annotated to genic (promoters, introns, exons) and regulatory elements (CpG islands, transcription factor binding sites, DNase hypersensitivity sites, enhancers) using the PoreMeth2 annotation module.

#### Nanopore RNA sequencing

Total RNA was extracted from hSCs after stimulation with sumatriptan (10µM) for 0, 6, 24 and 48 hours. Cells were harvested washed with DBPS twice and lysed using RNeasy Mini Kit (Qiagen SpA), according to the manufacturer’s protocol. RNA libraries were prepared in quadruplicate for each treatment timepoint using the SQK-PCB111-24 kit (Oxford Nanopore Technology) and sequenced on a PromethION P2 Solo instrument (FLO-PRO002 r9.4.1 flow cells). Raw data were basecalled and demultiplexed with Guppy (v6.5.7) and aligned to the GRCh37 reference genome via minimap2. Gene expression was quantified using the featureCounts function from the Rsubread package. Differential expression analysis was subsequently performed using DESeq2(*85*) Genes were filtered based on an adjusted p-value < 0.05 and an absolute log2 fold change (|log2FC|) ≥ 0.5. Heatmap and Volcano plots were made using the R packages ggplot2 (version 4.0.2) and pheatmap (version 1.0.13).

#### Immunofluorescence, Immunocytochemistry and RNAScope fluorescent in situ hybridization

The detailed methods were described in supplementary material.

#### Live-cell imaging 5-HTR internalization

hSCs were plated in 96-well black wall clear bottom plates (#165305, Thermo Fisher Scientific) (5 × 10^5^ cells/well) and maintained at 37 °C in 5% CO_2_ and 95% O_2_ for 24 h. hSCs were then transfected with 150 ng of plasmid DNA expressing EGFP-tagged 5-HT_1B_ or EGFP-tagged 5-HT_1D_ for 48h using jetOPTIMUS® DNA transfection reagent (#55-250; Polyplus,). 16h before imaging, hSC were infected with CellLight Early Endosomes-RFP, BacMam 2.0 (#C10587, Thermo Fisher Scientific) according to manufacturer’s instruction. hSCs were washed twice in HBSS/HEPES. Cells were preincubated with GR125743 (10µM, #HY-121392, DBA) or vehicle for 30 min and stimulated with sumatriptan (30 µM) or vehicle directly under the microscope. 5-HTR dynamics was imaged using Leica Stellaris 5 (Leica) confocal microscope, LAS X imaging software. Images were captured every 1 s. Processing and Rcoloc calculation were performed using ImageJ software.

### TGF-β3 and IL-15 ELISA kit assay

TGF-β3 and IL-15 levels were assayed in the mouse trigeminal nerve tissue homogenates at day 7 after treatments using a single-analyte enzyme-linked immunosorbent assay kit (#MBS824950 MyBioSource and #AC-A75519-96 Antibodies.com). TGF-β3 was also assayed human plasma samples (#ab272203, Abcam) according to the manufacturer’s protocol. The samples were assayed in triplicate. Data are expressed as pg/mg of protein.

### Data and statistical analysis

The results are expressed as the mean ± SEM. For multiple comparisons, a one-way ANOVA followed by a post-hoc Bonferroni’s test was used. The two groups were compared using Student’s t-test. For behavioral experiments with repeated measures, a two-way mixed-model ANOVA followed by a post-hoc Bonferroni’s test was used. Statistical analyses were performed on raw data using GraphPad Prism 8 (GraphPad Software Inc.). P-values less than 0.05 (P < 0.05) were considered significant. EC_50_, EC_80_, IC_50_ and IC_80_ values were determined from non-linear regression models using Graph Pad Prism 8. Raw count data from both sequencing runs were analyzed using R and associated packages. A principal component analysis (PCA) plot was generated to confirm appropriate clustering of the experimental conditions. Differential expression analysis was performed using the DESeq2 package (*85*). Genes were considered differentially expressed if they had a log2 fold change > 0.58 or < −0.58 (corresponding to approximately a 1.5-fold change) and an adjusted p-value <0.05. The statistical tests used and sample size for each analysis are shown in the Figure legends.

## List of Supplementary Materials

Supplementary methods Fig S1 to S3

Tables S1 to S4 Data file S1

## Author contributions

Conceptualization: AM, RN, FDL

Methodology: MC, MM, MB, EB, LB, LT, BP, DSMdA, AP, GDS, IS, GF, GP

Investigation: MC, MM, MB, EB, LB, LT, FDC, RDI, RG, BP, DSMdA, AP, GDS, IS, GF, GP

Funding acquisition: AM, RN, FDL Project administration: AM, RN, FDL Supervision: AM, RN, FDL, GS, RDI, RG

Writing – original draft: AM, RN, FDL, GS, MC, MM, MB, Writing – review & editing: AM, RN, FDL, GS, MC, MM, MB

## Data and materials availability

All data of this manuscript are openly available and are provided in the Main Text/Supplementary Information/ Data file and from Francesco De Logu and Romina Nassini.

## Supplementary Materials

### Supplementary Methods

#### AAV production and cell lysis

AAVPro-HEK293T cells (#632273, RRID:CVCL_B0XW, Takara) were cultured in DMEM with heat-inactivated FBS (10 %), penicillin/streptomycin (1%), sodium pyruvate (1%), and L-glutamine (2%). Cells were plated in a CellBIND Polystyrene CellSTACK 2 Chamber (#3310, Corning) for 48 h to reach 80% confluence. Cells were washed with PBS, detached with trypsin-EDTA 0.05% (EuroClone). Cells (250 x 10^6^) were seeded in a CellBIND Polystyrene CellSTACK 5 Chamber (#3311; Corning) for 24 h, until 80% of confluence. Cells were transfected with 2.5 mg of DNA containing the three plasmids (packaging, helper and gene of interest (GOI)) in a 1:1:1 molar ratio. To produce rAAV that infects with high efficiency Schwann cells, Rep/Cap 2/rh10 was used (pAAV2/rh10 #112866, RRID:Addgene_112866 Addgene). To mainly infect primary sensor neurons, AAV2 rep-AAV-PHP.S were used (pUCmini-iCAP-PHP.S #103006, RRID:Addgene_103006, Addgene). Total DNA was diluted in 176 ml of OptiMEM (Thermo Fisher Scientific) and combined with PEI (1:3 DNA to PEI ratio). After 15 min of incubation, 350 mL of DMEM supplemented with FBS (2%) was added to the OptiMEM/DNA/PEI mix and used to replace the complete medium in each chamber. Cells were maintained at 37 °C in 5% CO_2_ and 95% O_2_ for 72 h, before AAV particles were started to be collected. Cells were harvested and transferred into 50 mL conical tubes and centrifuged at 1000 × g for 10 min at 4°C. The supernatant was filtered through 0.45-μm PES membranes and then 25 mL of PEG solution (400 g of 40% polyethylene glycol + 24 g of NaCl in ddH2O to a final volume of 1.000 mL) was added to every 100 mL of collected supernatant. The total solution was slowly stirred at 4 °C for 1 h and then, kept for 3 h without stirring at 4°C to allow full precipitation of particles. The solution was then centrifuged at 2.800 × g for 15 min at 4°C, the supernatant was discarded, and the virus was resuspended in a10 ml of PBS/pluronic F68 (0.001%)/NaCl (200 mM) solution. Virus producing cell pellet was directly resuspended in 10 mL of PBS/Pluronic F68 (0.001%)/NaCl (200 mM) solution and cells were lysed by 4 cycles of freezing/thaw. Each cycle included a 30min step at −80°C, followed by a 10 min thaw at 37°C, interspersed with vortexing. Sample was then centrifuged at 3.200 × g for 15 min at 4°C and supernatant, containing the AAV particles was collected, while cell debris were discarded. Samples obtained by medium treatment and cell lyses, containing the AAV particles, were finally mixed, incubated with benzonase (50 UI/mL) at 37°C for 45 min to digest residual plasmids and residual genomic DNA / cellular RNA. Then, sample was centrifuged at 2.400 × g for 10 min at 4°C. The clarified supernatant was transferred to new tubes and was kept overnight at 4°C before purification. Purification using a gradient of iodixanol was performed. Starting with a 60% iodixanol solution (OptiPrep; STEMCELL Technologies), a iodixanol gradient was prepared with a 15% solution [4.5 mL of iodixanol (60%) + 13.5 mL of NaCl/PBS-MK buffer (1M)], a 25% solution [5 mL of iodixanol (60%) + 7 mL of PBS-MK buffer (1×) + 30 μL of phenol red], a 40% solution [6.7 mL of iodixanol (60%) + 3.3 mL of PBS-MK buffer (1×)], and a 60% solution [10 mL of iodixanol (60%) + 45 μL of phenol red]. Each solution was added into a 39-mL Quick-Seal tube (Beckman Coulter) using 18 G needle syringe in the following order: 8 mL of the 15% iodixanol solution, 6 mL of the 25% iodixanol solution, 5 mL of the 40% iodixanol solution, and 5 mL of the 60% iodixanol solution. Finally, tubes were filled with the sample, sealed and centrifuged in a Type 70 Ti rotor (Beckman Coulter, California, USA) at 350.000 x g at 10°C for 90 min and then pierced with a 16 G needle on top and an 18 G needle at the interface between the 60% and 40% iodixanol gradients. Viral particles contained in the 40% iodixanol layer were fractioned in 1.5 mL microcentrifuge tubes and concentrated using Amicon Ultra-15 centrifugal filter units (molecular weight cut-off, 100 kDa). Before the concentration step, membranes were activated with 15 mL of 0.1% Pluronic F68 in PBS solution that were discarded and replaced with 15 mL of 0.01% Pluronic F68 in PBS solution. The tubes were centrifuged at 3.000 rpm for 5 min at 4 °C. The supernatant was discarded, and 15 mL of 0.001% Pluronic F68 in PBS + NaCl 200mM solution was added and centrifuged at 3.000 rpm for 5 min at 4°C. The sample was then added and centrifuged at 3.500 rpm for 8 min at 4°C and flowthrough was discarded. During concentration process, formulation buffer (0.001% Pluronic F68 in PBS) was also added to the sample after few centrifugation steps to replace iodixanol and to avoid toxicity in animals after AAV injection. The viral title was quantified using RT-qPCR.

#### Immunofluorescence

Trigeminal nerve tissues and DRG were collected mice deeply anesthetized and transcardially perfused with PBS, were fixed for 24-48 h in 10% formalin, paraffin-embedded and cut with microtome at 5 µm. Tissues were incubated with different primary antibodies: anti-5HT_1D_ (#OSS00185W, Rabbit, 1:1000, Thermo Fisher), anti-5HT_1B_ (#ASR-022, RRID:AB_10561260, Rabbit, 1:100, Alomone labs), anti-DNMT3A (#ab188470, RRID:AB_3073896, Rabbit, 1:500, Abcam), anti-DNMT3B (#ab2851, RRID:AB_303356, Rabbit, 1:500, Abcam), anti-TGF-β3 (#ab15537, RRID:AB_2202305, Rabbit, 1:100 Abcam), anti-TGF-βR1 (#30117-1-AP, RRID:AB_3086235,Rabbit, 1:200, Proteintech), anti-TGF-βR2 (#PA5-35076, RRID:AB_2552386, Rabbit, 1:100, Invitrogen), S100 (#MA1-26621, RRID:AB_2552386Mouse, 1:50, Thermo Fisher) diluted in fresh blocking solution (PBS1X pH 7.4, 5% normal goat serum (NGS) or normal donkey serum (NDS); or PBS1X pH 7.4, 3% Bovine Serum Albumin (BSA) overnight at 4°C. The primary antibody was used after antibody retrieval performed with tri-sodium citrate buffer (pH 6) for TGF-β3, TGF-βR1, TGF-βR2 and for DNMT3B; tris-EDTA buffer (pH 8) for 5HT_1B_ and tris-EDTA (pH 9) for 5HT_1D_ and DNMT3A and following a 1 h incubation with the specific blocking solution. Sections were washed in PBS and then incubated with the appropriate fluorescent polyclonal secondary antibodies Alexa Fluor® 488 (#A32731, goat polyclonal anti-rabbit, 1:600, Invitrogen,), Alexa Fluor® 488 (#A32790, RRID:AB_2633280, donkey polyclonal anti-rabbit, 1:600, Invitrogen), Alexa Fluor® 647 (#A31571, RRID:AB_162542, donkey polyclonal anti-mouse, 1:600, Invitrogen, Carlsbad, California), Alexa Fluor® 647 (#A21236, RRID:AB_2535805,goat polyclonal anti-mouse, 1:600, Invitrogen), Alexa Fluor® 594 (#A11005, RRID:AB_141372,goat polyclonal anti-mouse, 1:600, Invitrogen) for two hours RT. At the end tissues were covered using the mounting medium with DAPI (#ab104139, Abcam) and stored at 4°C utile imaging. Human sciatic nerve (#0062-HP-261, Gentaur) were incubated with primary antibodies: anti-5HT_1D_ (#OSS00185W, Rabbit, 1:1000, Thermo Fisher), anti-5HT_1B_ (#ASR-022, RRID:AB_2756794, Rabbit, 1:100, Alomone labs,) followed by the same steps of mice tissues. All slides were visualized and analyzed using a Zeiss Axio Imager 2 microscope with Z-stacks in the Apotome mode (Zeiss) or Leica Stellaris 5 confocal microscope (Leica). Pearson correlation (Rcoloc) values in the colocalization analysis were calculated using the colocalization Plugin of the ImageJ (v.1.54 f; National Institutes of Health, Bethesda). We underline, as a limitation of the colocalization method, that detection of the biomarkers localized in two or more different intracellular compartments of the same cell may be affected by the slice orientation that may result in incomplete colocalization.

#### Immunocytochemistry

Human, primary and mouse Schwann cells were fixed in 10% neutral buffered formalin for 10 minutes, rinsed briefly in PBS and permeabilized with 0,5% Triton X-100 for 10 minutes. Cells were treated with a blocking solution 2% BSA in PBS for 1h and were incubated with different primary antibodies: anti-5HT_1D_ (#OSS00185W, Rabbit, 1:100, Thermo Fisher), anti-5HT_1B_ (#ASR-022, RRID:AB_10561260, Rabbit, 1:100, Alomone labs), anti-TGF-β3 (#ab15537, RRID:AB_2202305, Rabbit, 1:100, Abcam), and SOX10 (#AF-2864, RRID:AB_442208, Goat, 1:20, Biotechne) for 1h RT. All incubations were performed on a shaker to enhance the penetration of the solutions. Then cells were incubated with the fluorescent polyclonal secondary antibodies Alexa Fluor® 488 (#A32790, donkey polyclonal anti-rabbit, 1:600, Invitrogen), Alexa Fluor® 594 (#A11058, donkey polyclonal anti-goat, 1:600, Invitrogen) for two hours. At the end cells were covered using the mounting medium with DAPI (#ab104139, Abcam). Cells were imaged on a Leica Stellaris 5 confocal microscope (Leica), and images were processed with ImageJ.

#### RNAScope fluorescent in situ hybridization and immunofluorescence

Fluorescent *in situ* hybridization was performed using the RNAscope™ Multiplex Fluorescent V2 Assay (#323100, ACDbio), according to the manufacturer’s protocol. Trigeminal nerves were collected from control and treated animals, fixed for 24–48 h in 10% formalin, paraffin-embedded, and sectioned at 7 µm using a microtome. Slides were pretreated by incubation at 60°C for 1 h on a Thermobrite system, followed by rinsing in xylene and 100% ethanol. RNAscope hydrogen peroxide (#322335; ACDbio) was then applied for 10 min RT to inactivate endogenous peroxidases. Sections were briefly rinsed in distilled water and subjected to antigen retrieval in an oil bath at 99°C for 15 min. After washing in distilled water and 100% ethanol, slides were left to dry overnight RT. The following day, slides were treated with 2–4 drops of Protease Plus (#322331; ACDbio) and incubated at 40°C in a HybEZ II Oven (ACDbio) for 30 min.

Sections were hybridized for 2 h with probes targeting *HTR1B, HTR1D*, and *BETAGLYCAN* in human (#449831, #1113191, #441031; Bio-Techne) and *Htr1b*, *Htr1d, Tgfb3*, and *Betaglycan* in mouse (#315861, #315871, #406211, #406221; Bio-Techne) trigeminal nerves. Signal amplification was performed by sequential incubation with AMP1, AMP2, and AMP3 (#323110; ACDbio) at 40°C. Samples were subsequently incubated with HRP-C1 and the fluorophore TSA-Vivid 650 (#7536; ACDbio) diluted 1:1500 in TSA Buffer (#322810; ACDbio) for 30 min at 40°C in the HybEZ II Oven, followed by incubation with HRP Blocker for 15 min. After the RNAscope procedure, immunohistochemistry was performed. Slides were incubated with 5% normal goat serum (NGS) in 1× PBS containing 0.1% Tween-20 for 1 h RT, followed by incubation with the primary antibody S100 (#MA1-26621, RRID:AB_795376, mouse; 1:50 Thermo Fisher) or NeuN (#MAB377, RRID:AB_2298772 mouse; 1:250) diluted in blocking solution overnight at 4°C. Sections were washed in PBS and incubated with Alexa Fluor® 488 (#A32723, RRID:AB_2633275, goat polyclonal anti-mouse, 1:600, Invitrogen) for 2 h RT. Slides were mounted with DAPI-containing mounting medium (#ab104139, Abcam). Fluorescent images were acquired using a Leica Stellaris 5 (Leica).

## Supplementary Figures

**Figure S1.**
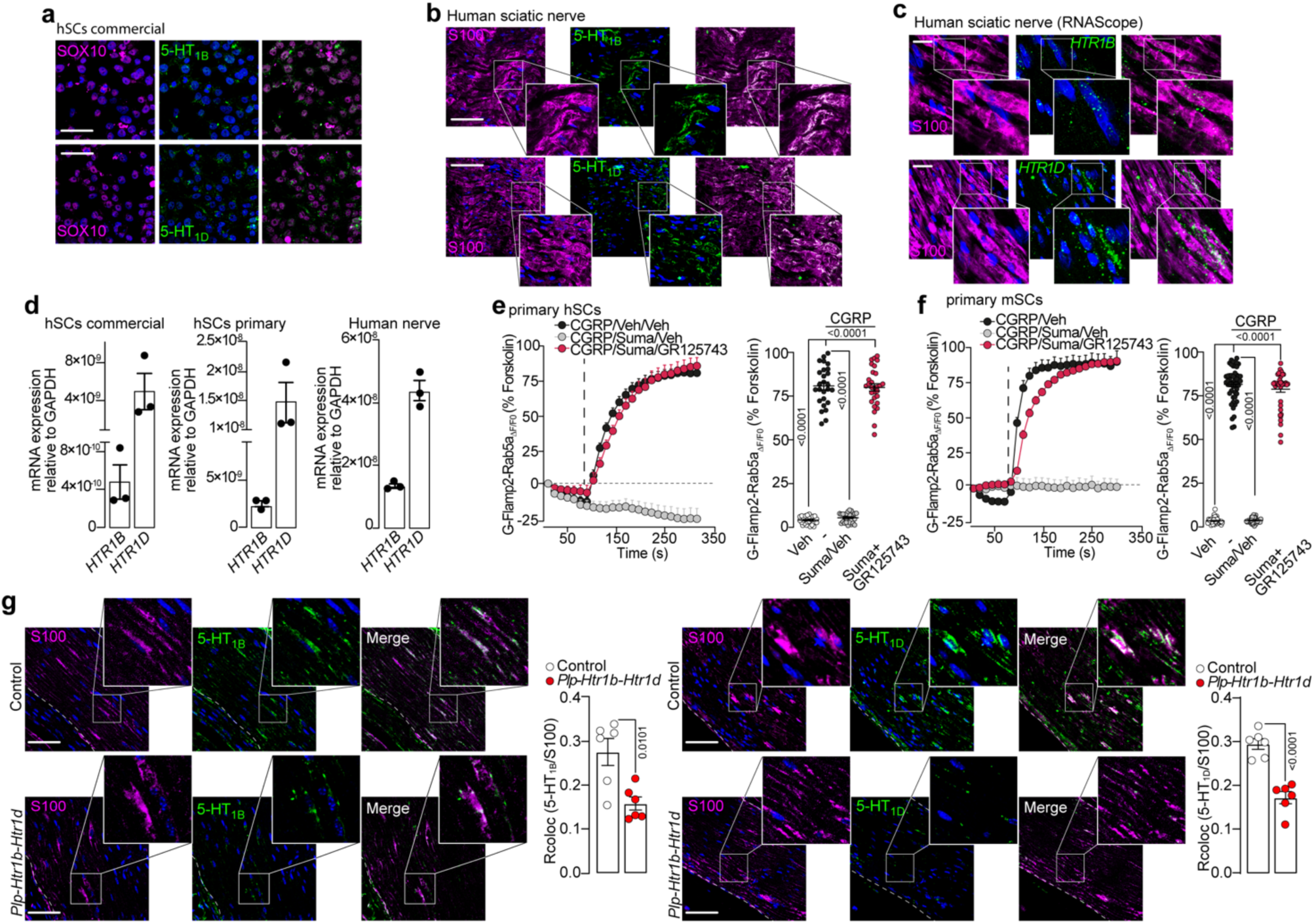
(**a**) Representative images of 5-HT_1B_ e 5-HT_1D_ expression in human Schwann cells (hSCs, scale bar: 10 μm, n=6) and (**b)** in human sciatic nerve tissues (scale bar: 20 μm, n=6). SOX10 and S100 are markers of SCs. **(c)** Representative RNAscope images of *HTR1B* and *HTR1D* mRNA and S100 protein expression in human sciatic nerve (scale bar: 20 μm, n=6). **(d)** RT-qPCR for *HTR1B* and *HTR1D* mRNA in commercial and primary (isolated from lingual, sublingual and inferior alveolar nerves biopsies) hSCs and in human nerve tissues. **(e)** Typical traces and cumulative data of the effects of sumatriptan (Suma, 30 μM) on CGRP (3 μM)-evoked Rab5a cAMP formation in the presence of GR125743 (1 μM) or vehicle (Veh) in primary isolated hSCs (cells number: Veh/Veh/Veh=29, CGRP/Veh/Veh =26, CGRP/Suma/Veh =26, CGRP/Suma/GR125743 =29) and in **(f)** primary mouse SCs (mSCs) (cells number: Veh/Veh/Veh=40, CGRP/Veh/Veh =50, CGRP/Suma/Veh =37, CGRP/Suma/GR125743 =41). **(g)** Representative images colocalization data (Rcoloc) of 5-HT_1B_ and 5-HT_1D_ silencing in SCs in *Plp-Htr1b-Htr1d* and Control mice. Data are mean ± s.e.m. Student’s t-test and 1-way ANOVA, Bonferroni correction.

**Figure S2.**
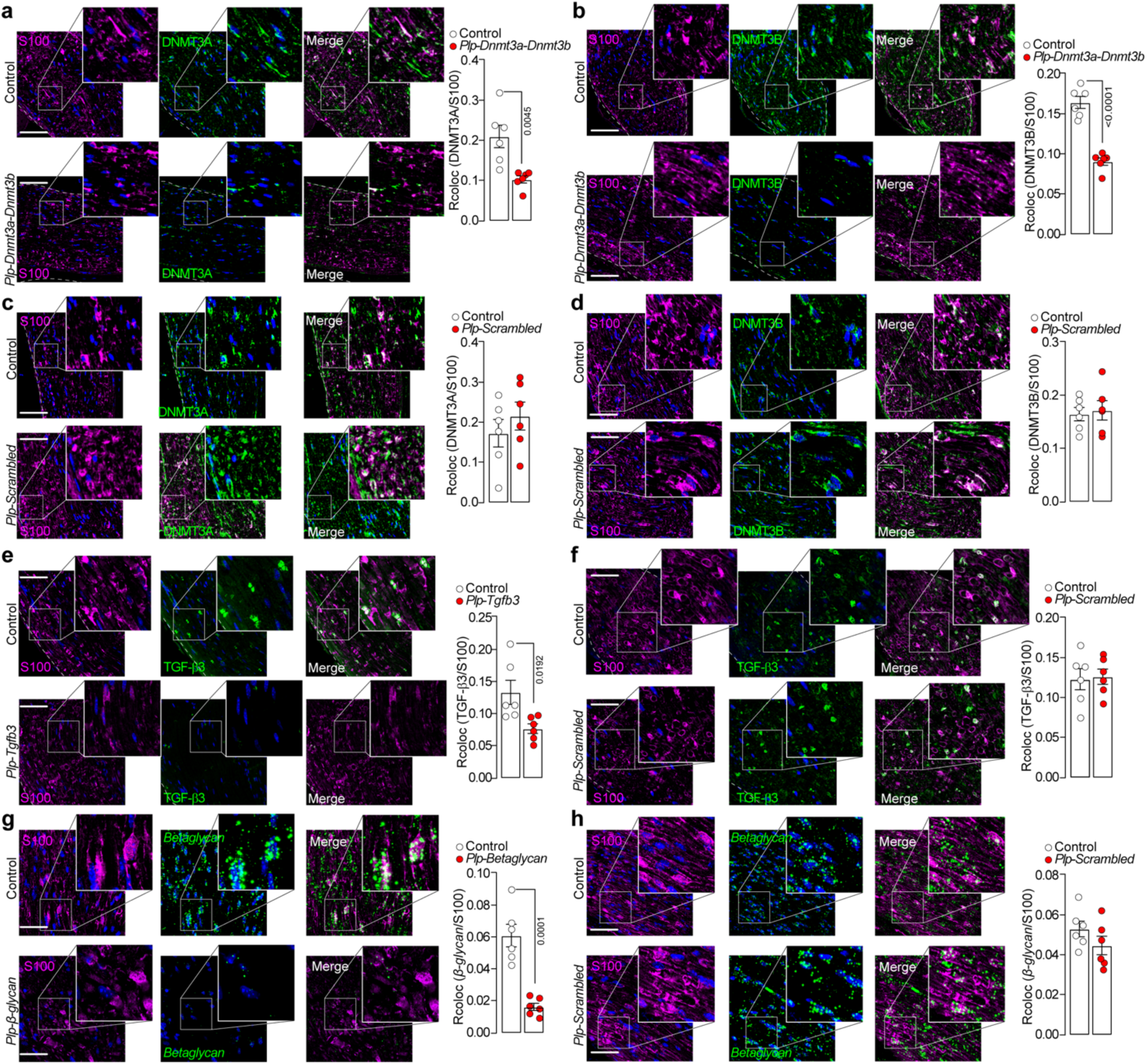
(**a-d**) Representative images of DNMT3A or DNMT3B and S100 protein expression and colocalization data (Rcoloc) in trigeminal nerve from *Plp-Dnmt3a-Dnmt3b*, *Plp-Scrambled* or Control mice (scale bar: 20 μm, n=6). **(e,f)** Representative images of TGF*-*β3 and S100 and Rcoloc in trigeminal nerve from *Plp-Tgfb3*, in *Plp-Scrambled* or Control mice (scale bar: 20 μm, n=6). **(g,h)** RNAscope images of *Betaglycan* and S100 and Rcoloc in trigeminal nerve from *Plp-Betaglycan, Plp-Scrambled* or control mice (scale bar: 20 μm, n=6). Data are mean ± s.e.m. (n=6 mice/group). Student’s t-test.

**Figure S3.**
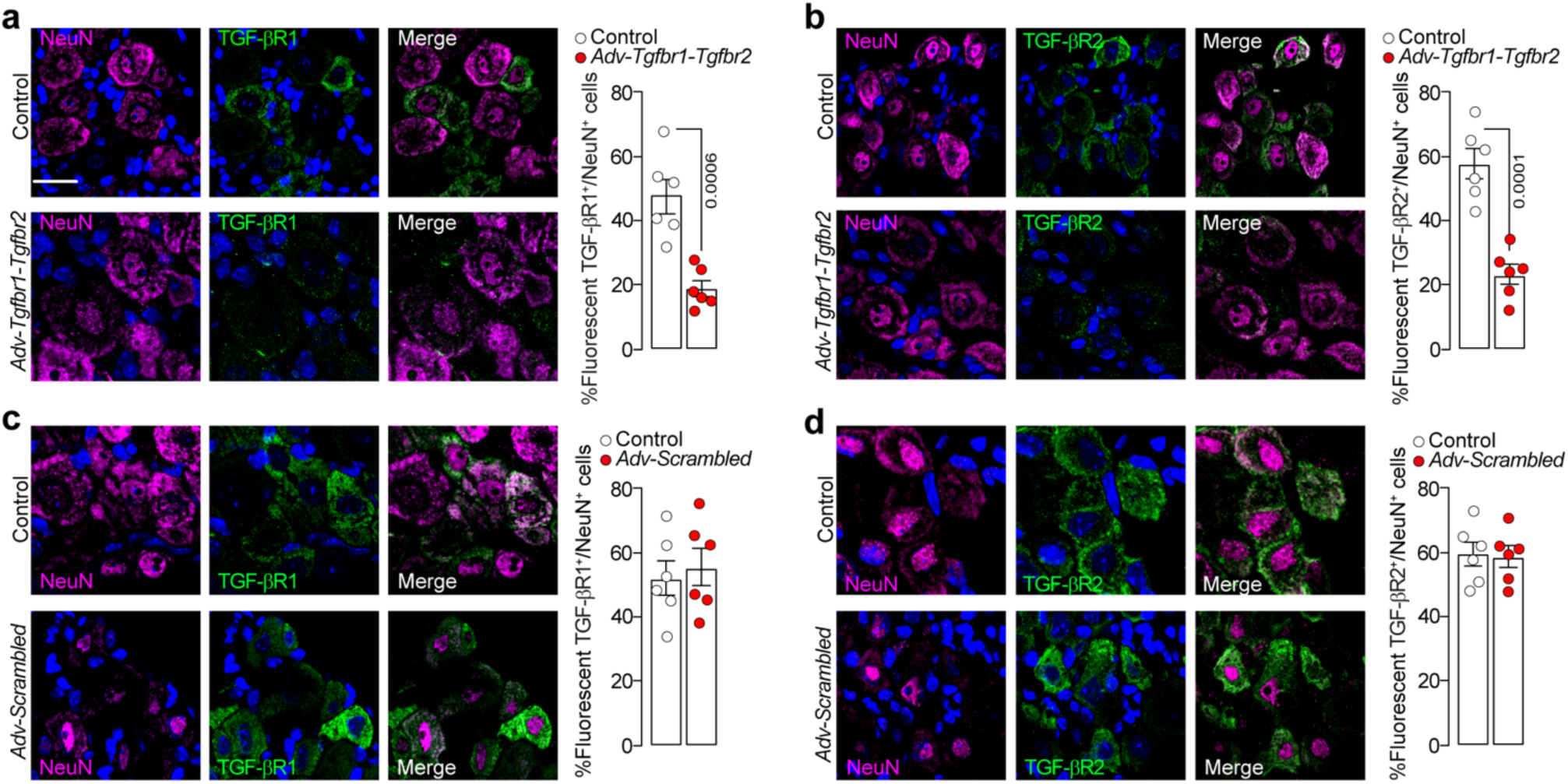
Representative images of TGF-βR1 or TGF-βR2 and NeuN protein expression in dorsal root ganglia (DRG) from Control mice and **(a,b)** *Adv-Tgfbr1-Tgfbr2* and **(c,d)** *Adv-Scrambled* mice (scale bar: 20 μm, n=6). Data are mean ± s.e.m. (n=6 mice/group). Student’s t-test.

## Supplementary Tables

**Table S1:**
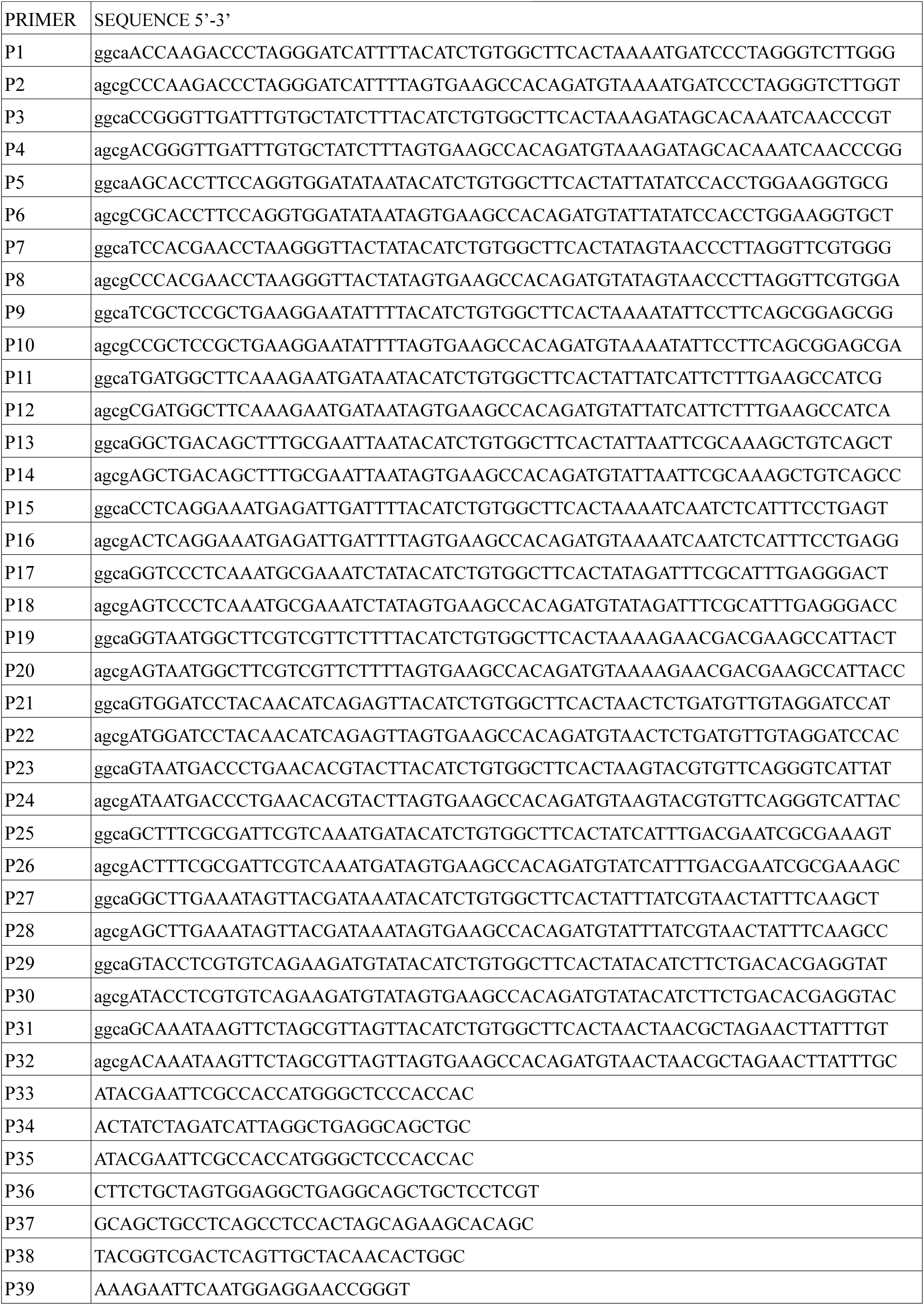
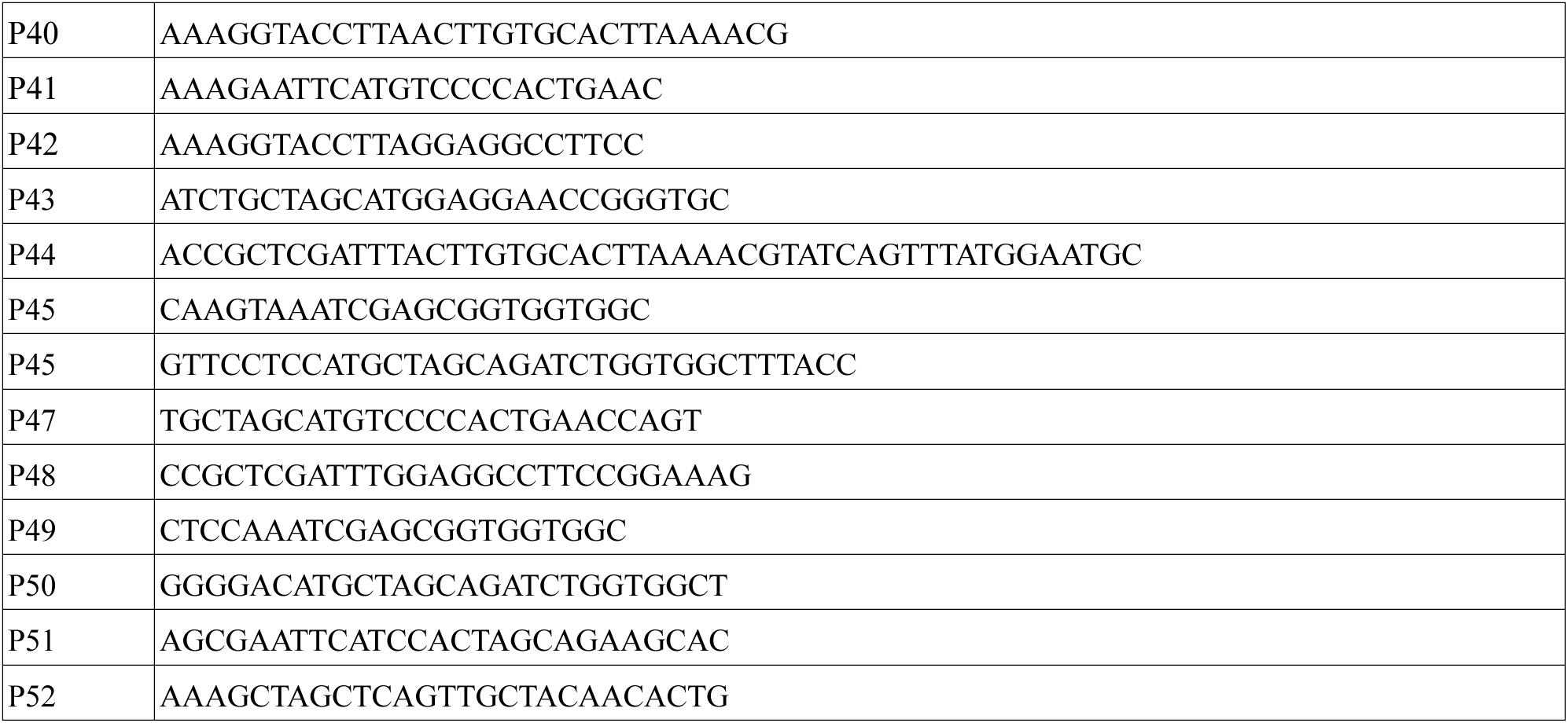
Primers for plasmids construction.

**Table S2:**
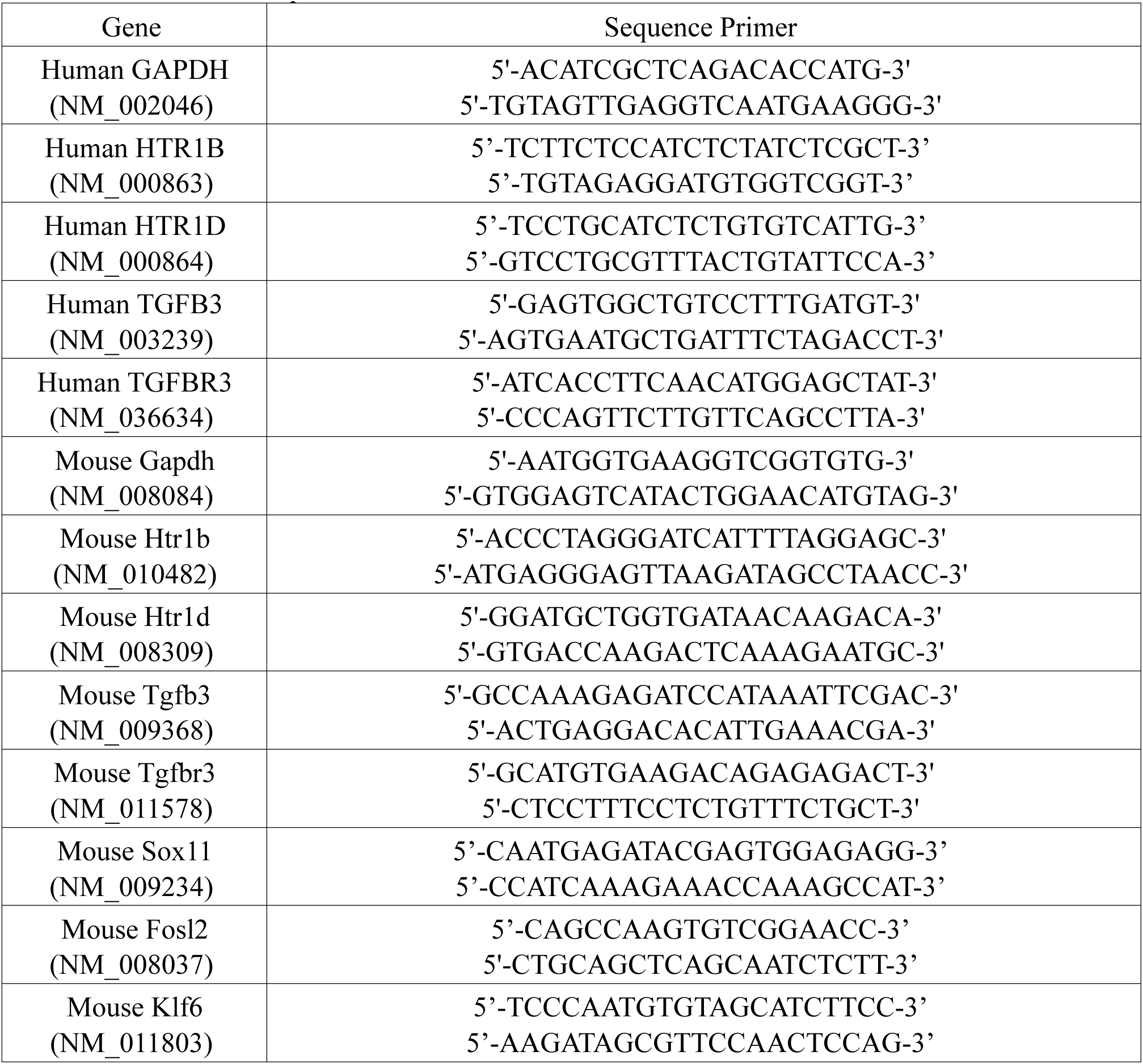
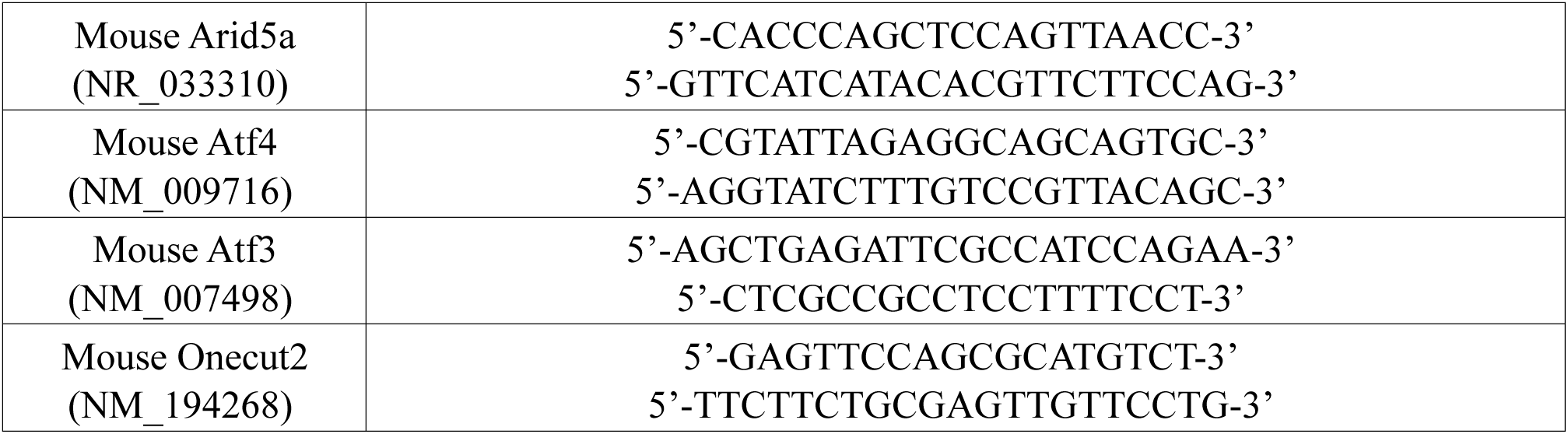
Primer for RT-qPCR.

**Table S3:**
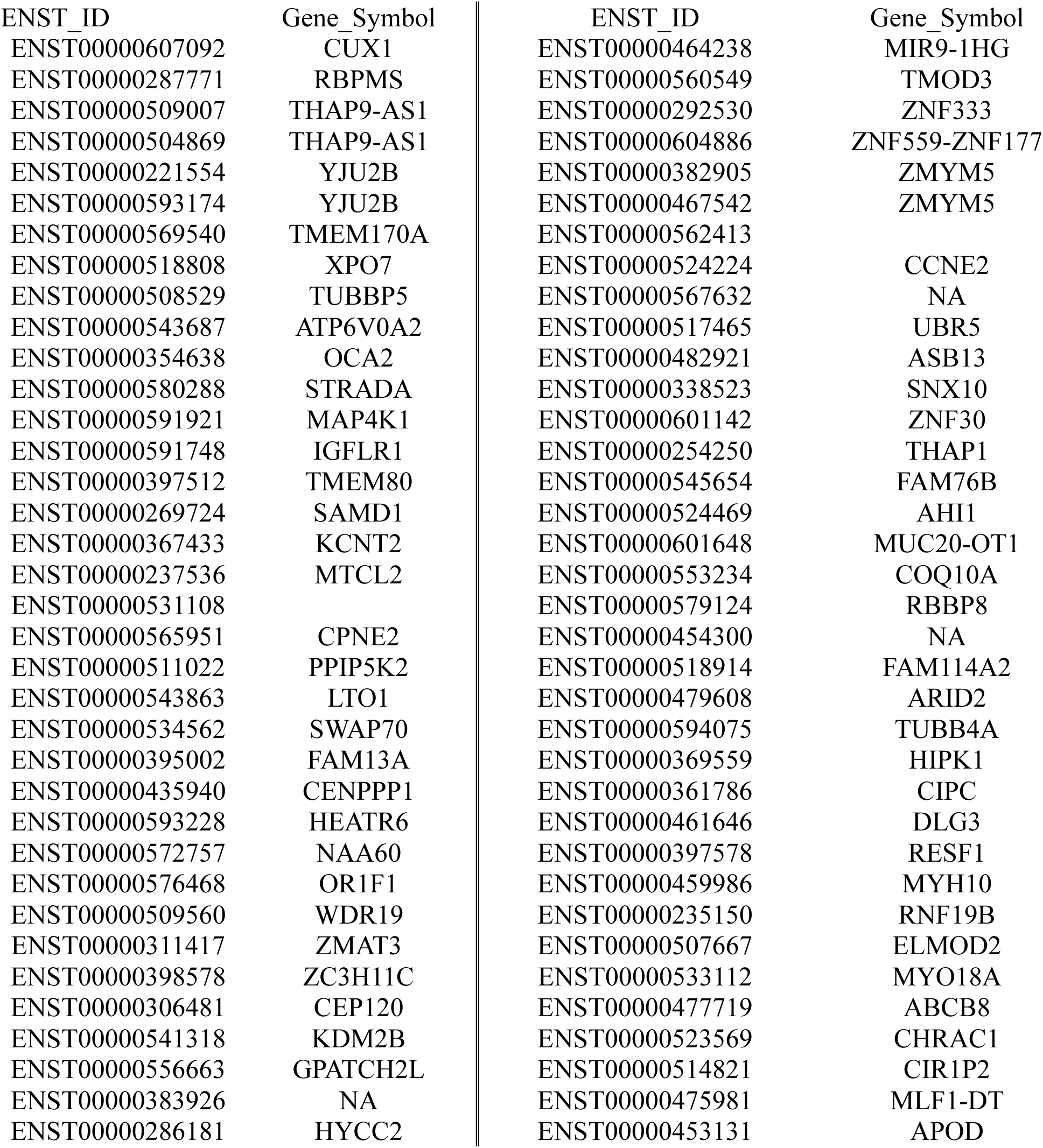
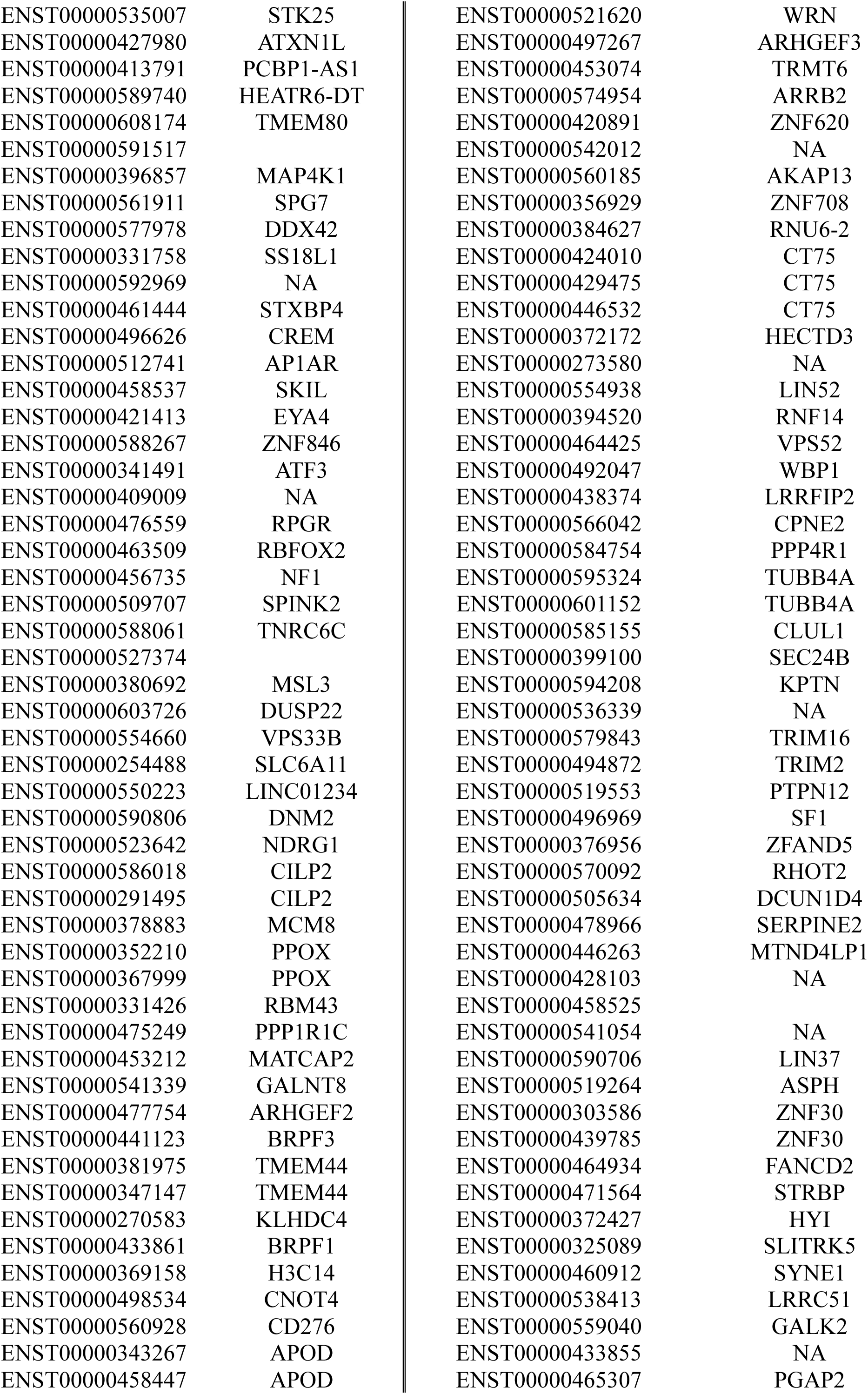
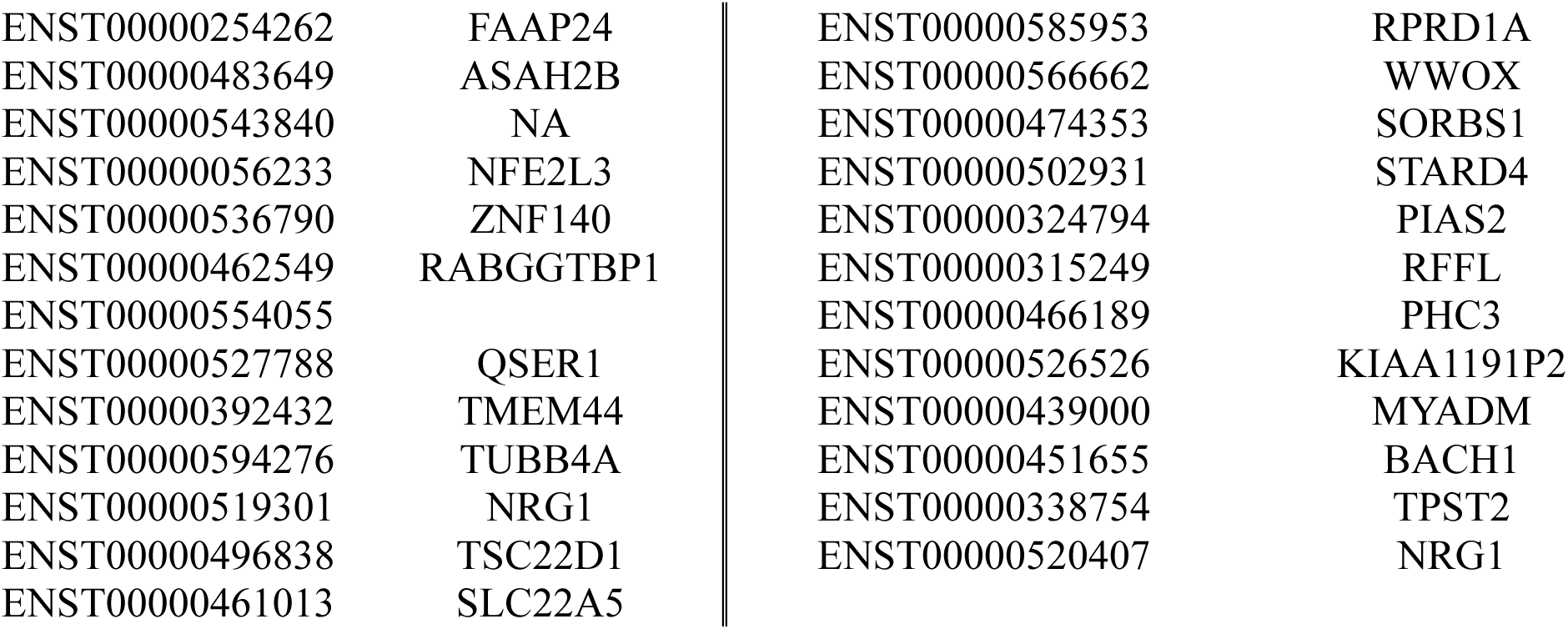
Heatmap of differentially expressed genes DEGs in hSCs.

**Table S4:**
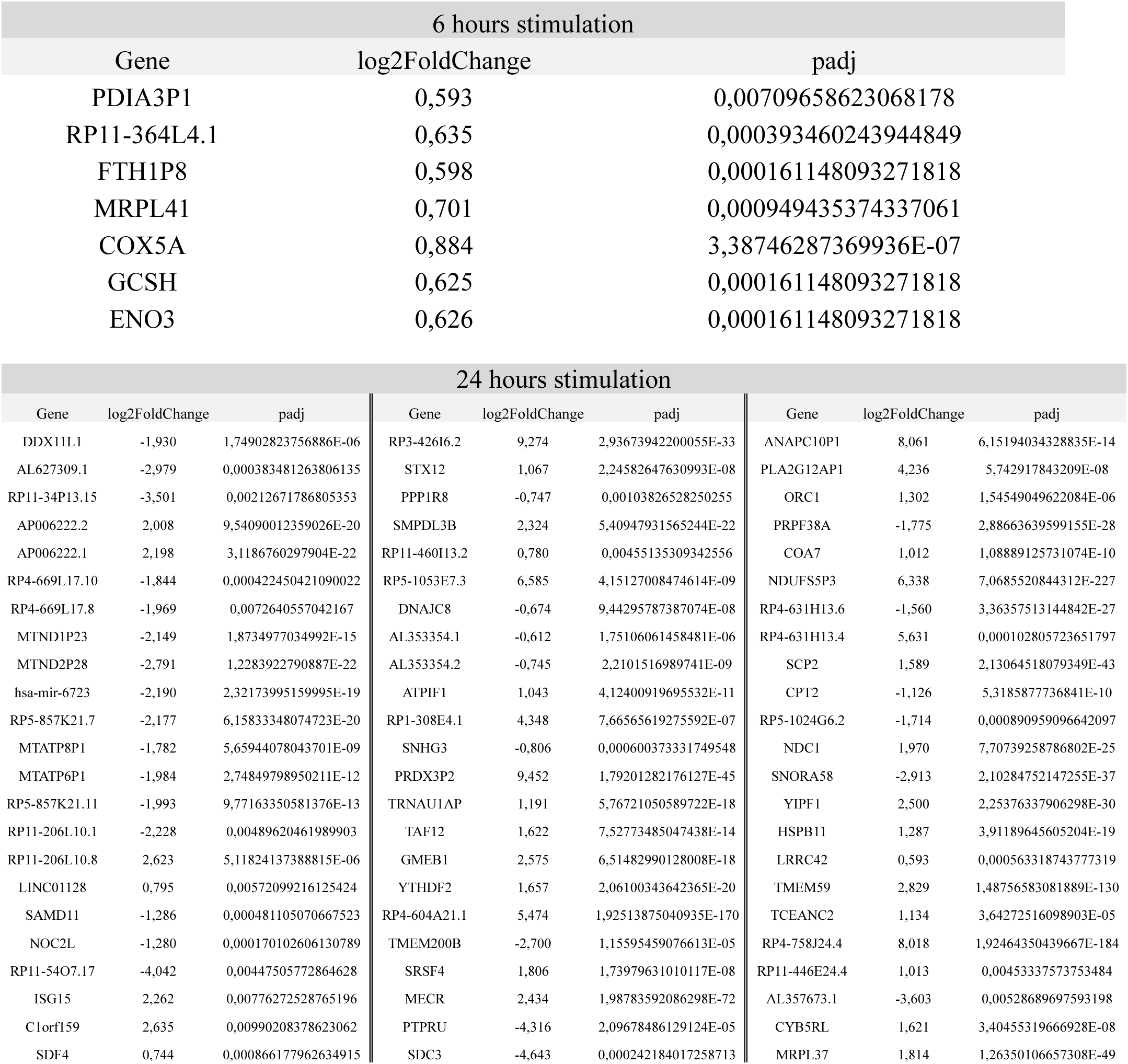
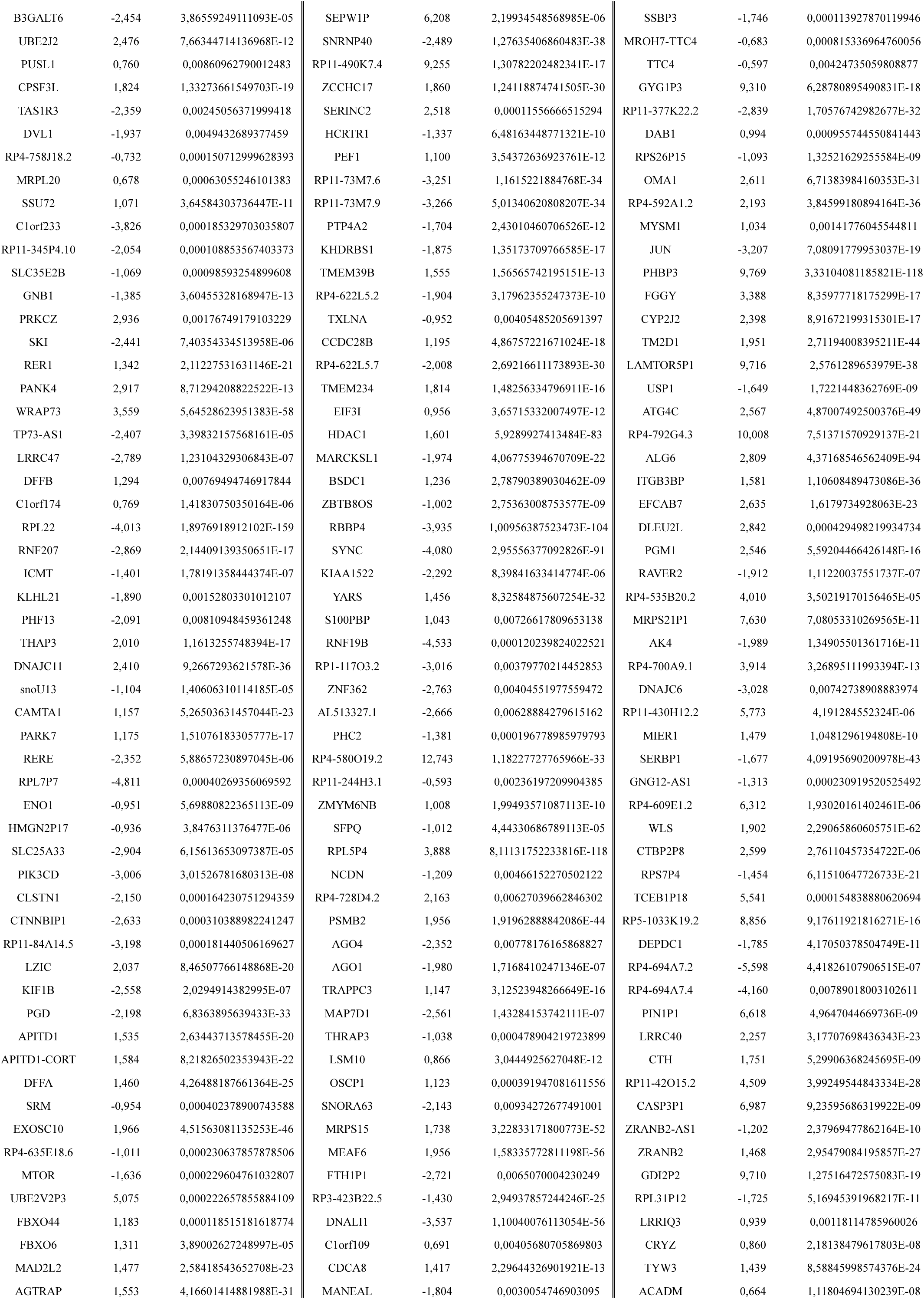
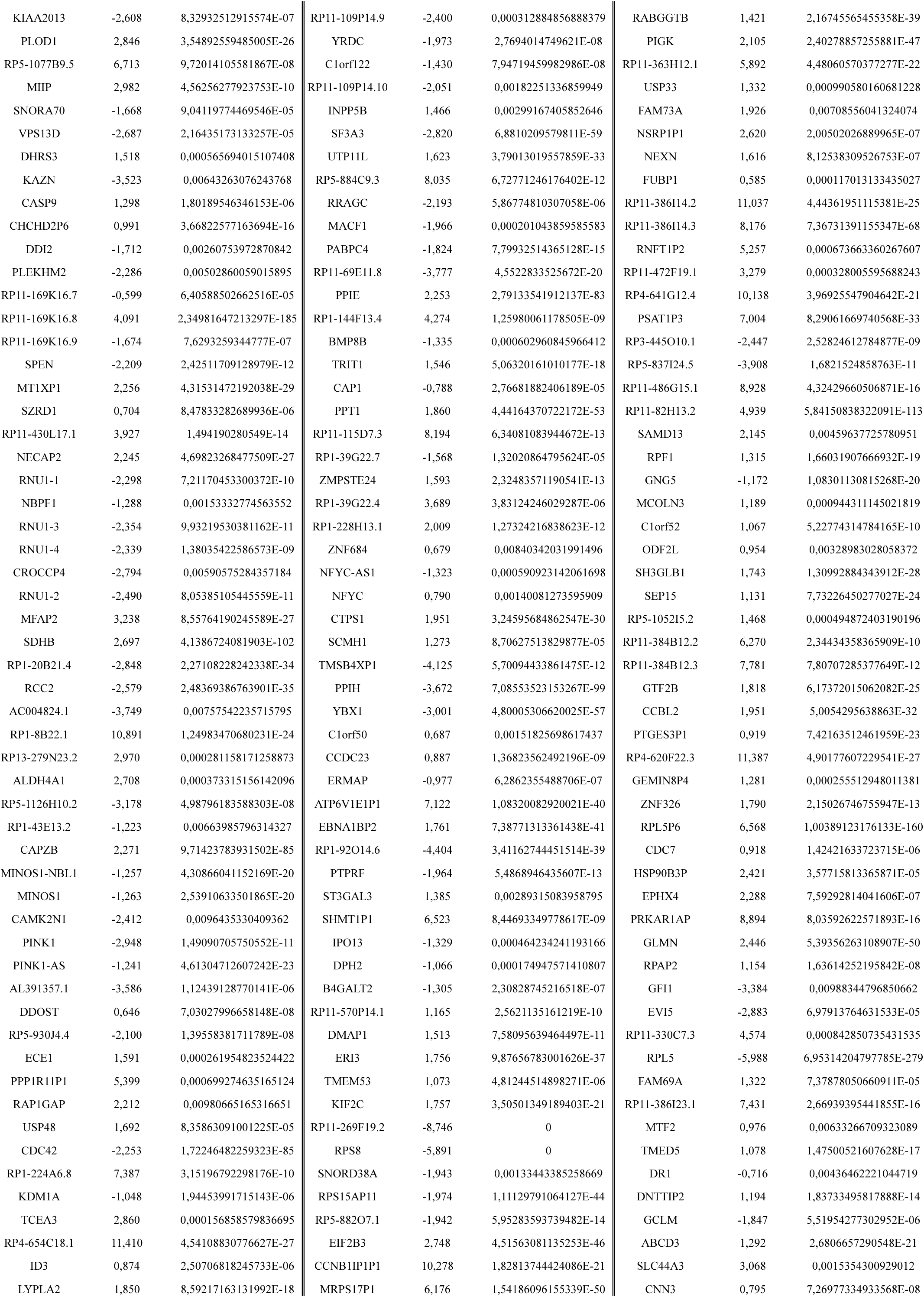
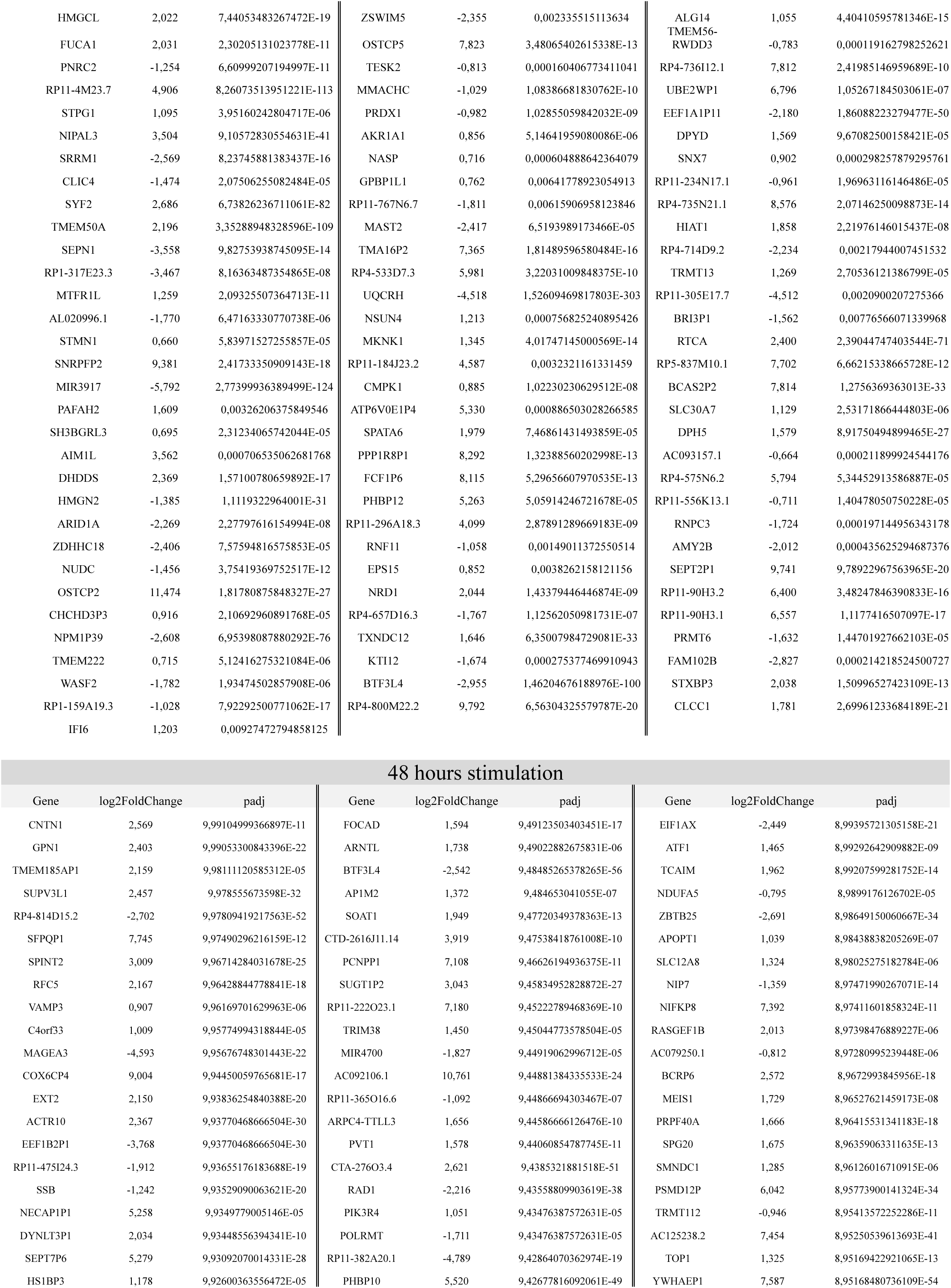
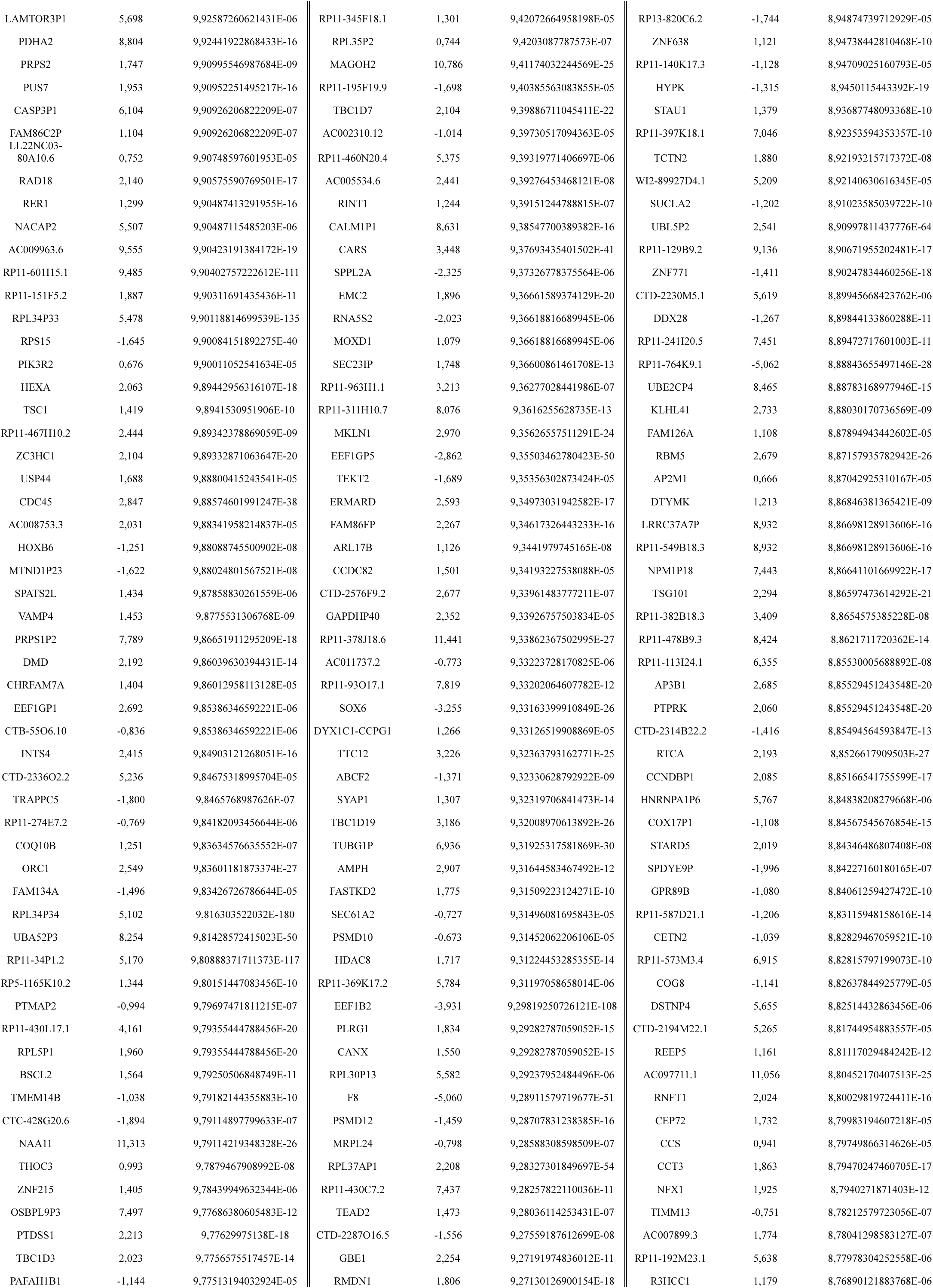
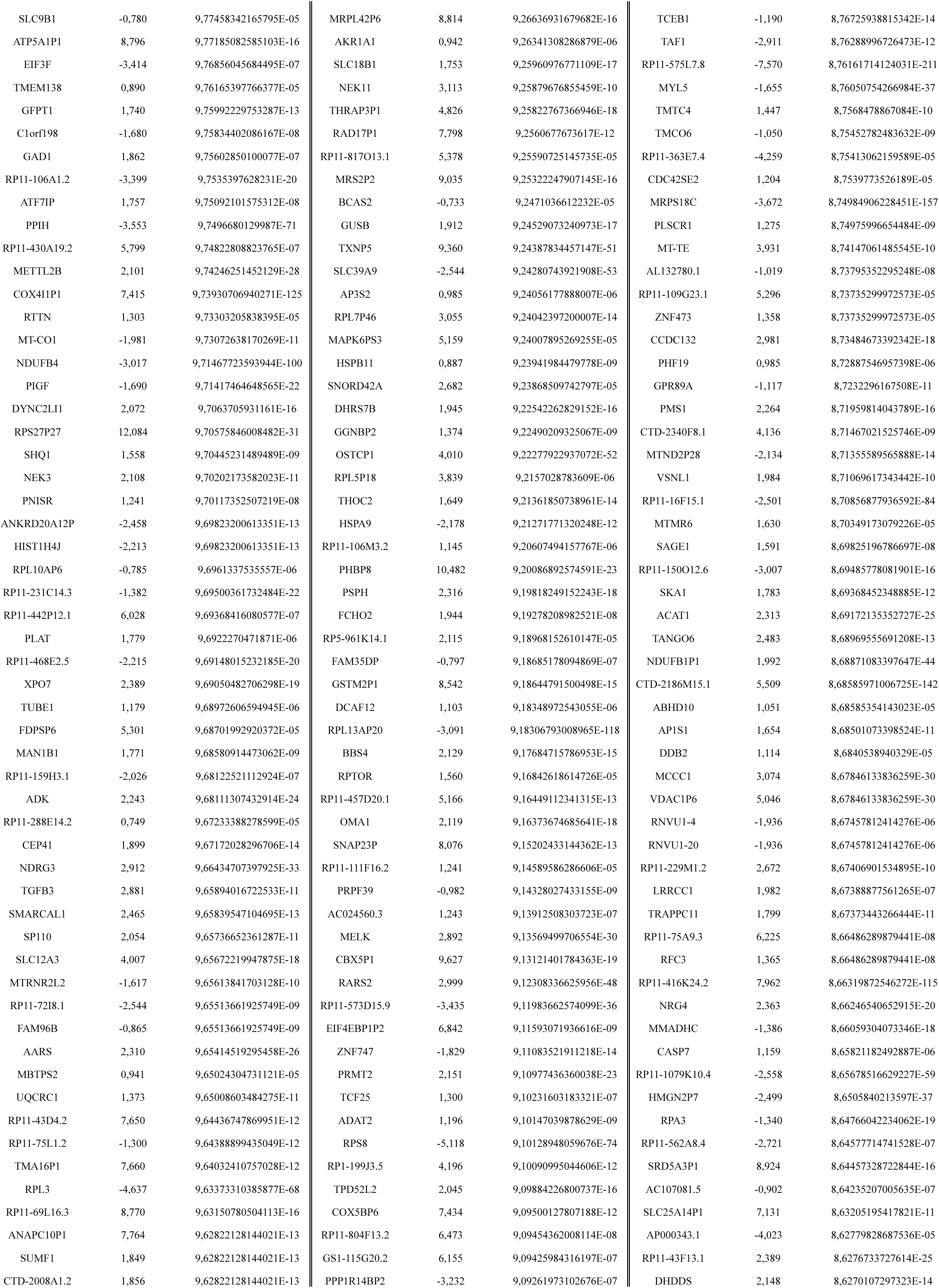
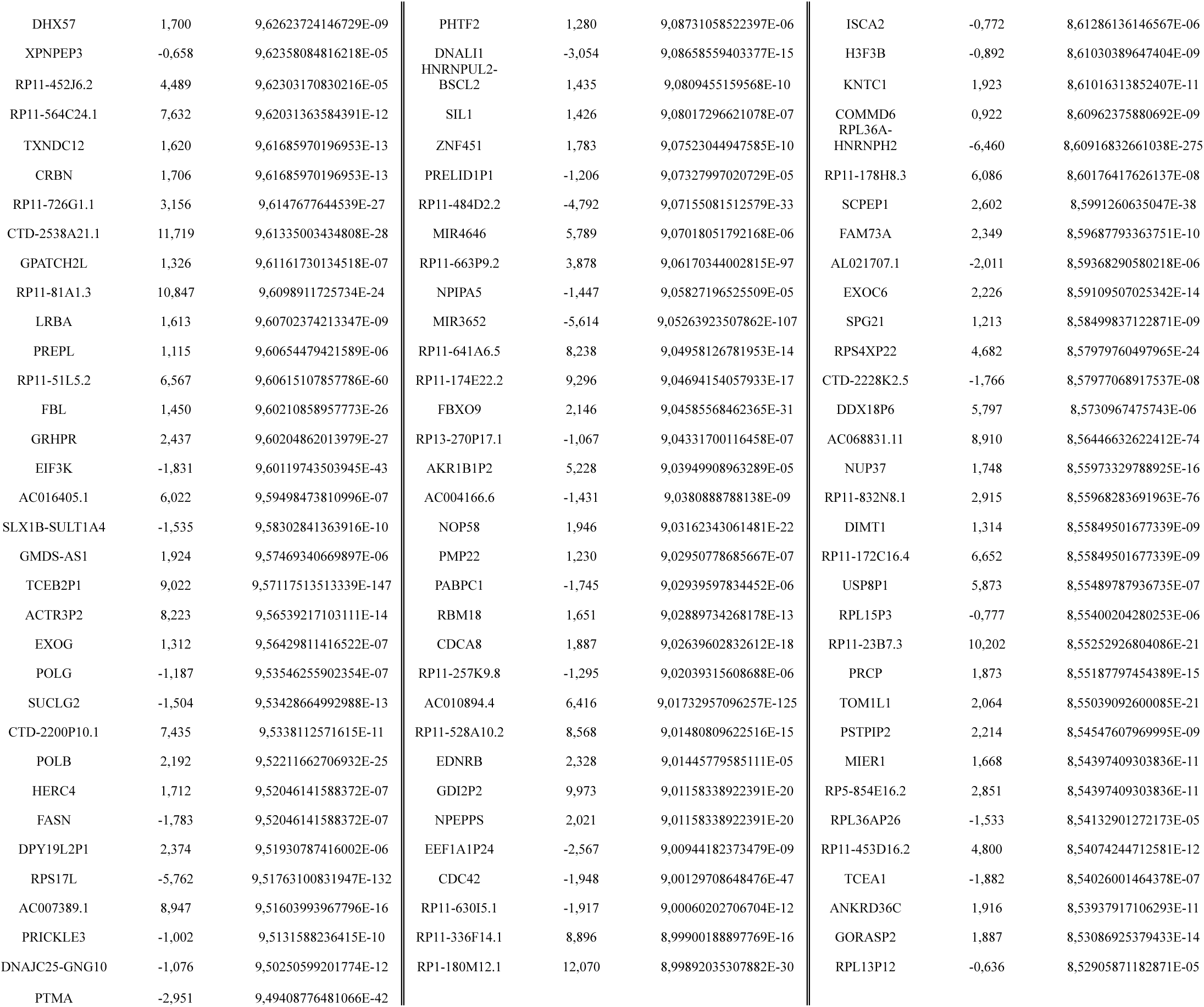
List of differentially expressed genes at 6h, 24h and 48h of Sumatriptan stimulation.

