## Supplementary information for "Triptans reprogram Schwann cells to drive medication-overuse headache via β-glycan/TGF-β3 signaling"

**Supplementary Materials**

**Supplementary Methods**

**AAV production and cell lysis**

AAVPro-HEK293T cells (#632273, RRID:CVCL_B0XW, Takara) were cultured in DMEM with heat-inactivated FBS (10 %), penicillin/streptomycin (1%), sodium pyruvate (1%), and L-glutamine (2%). Cells were plated in a CellBIND Polystyrene CellSTACK 2 Chamber (#3310, Corning) for 48 h to reach 80% confluence. Cells were washed with PBS, detached with trypsin-EDTA 0.05% (EuroClone). Cells (250 x $10^6$) were seeded in a CellBIND Polystyrene CellSTACK 5 Chamber (#3311; Corning) for 24 h, until 80% of confluence. Cells were transfected with 2.5 mg of DNA containing the three plasmids (packaging, helper and gene of interest (GOI)) in a 1:1:1 molar ratio. To produce rAAV that infects with high efficiency Schwann cells, Rep/Cap 2/rh10 was used (pAAV2/rh10 #112866, RRID:Addgene_112866 Addgene). To mainly infect primary sensor neurons, AAV2 rep-AAV-PHP.S were used (pUCmini-iCAP-PHP.S #103006, RRID:Addgene_103006, Addgene). Total DNA was diluted in 176 ml of OptiMEM (Thermo Fisher Scientific) and combined with PEI (1:3 DNA to PEI ratio). After 15 min of incubation, 350 mL of DMEM supplemented with FBS (2%) was added to the OptiMEM/DNA/PEI mix and used to replace the complete medium in each chamber. Cells were maintained at 37 °C in 5% $CO_2$ and 95% $O_2$ for 72 h, before AAV particles were started to be collected. Cells were harvested and transferred into 50 mL conical tubes and centrifuged at 1000 × g for 10 min at 4°C. The supernatant was filtered through 0.45-µm PES membranes and then 25 mL of PEG solution (400 g of 40% polyethylene glycol + 24 g of NaCl in ddH2O to a final volume of 1.000 mL) was added to every 100 mL of collected supernatant. The total solution was slowly stirred at 4 °C for 1 h and then, kept for 3 h without stirring at 4°C to allow full precipitation of particles. The solution was then centrifuged at 2.800 × g for 15 min at 4°C, the supernatant was discarded, and the virus was resuspended in a10 ml of PBS/pluronic F68 (0.001%)/NaCl (200 mM) solution. Virus producing cell pellet was directly resuspended in 10 mL of PBS/Pluronic F68 (0.001%)/NaCl (200 mM) solution and cells were lysed by 4 cycles of freezing/thaw. Each cycle included a 30min step at -80°C, followed by a 10 min thaw at 37°C, interspersed with vortexing. Sample was then centrifuged at 3.200 × g for 15 min at 4°C and supernatant, containing the AAV particles was collected, while cell debris were discarded. Samples obtained by medium treatment and cell lyses, containing the AAV particles, were finally mixed, incubated with benzonase (50 UI/mL) at 37°C for 45 min to digest residual plasmids and residual genomic DNA / cellular RNA. Then, sample was centrifuged at 2.400 × g for 10 min at 4°C. The clarified supernatant was transferred to new tubes and was kept overnight at 4°C before purification. Purification using a gradient of iodixanol was performed. Starting with a 60% iodixanol solution (OptiPrep; STEMCELL Technologies), a iodixanol gradient was prepared with a 15% solution [4.5 mL of iodixanol (60%) + 13.5 mL of NaCl/PBS-MK buffer (1M)], a 25% solution [5 mL of iodixanol (60%) + 7 mL of PBS-MK buffer (1×) + 30 μL of phenol red], a 40% solution [6.7 mL of iodixanol (60%) + 3.3 mL of PBS-MK buffer (1×)], and a 60% solution [10 mL of iodixanol (60%) + 45 μL of phenol red]. Each solution was added into a 39-mL Quick-Seal tube (Beckman Coulter) using 18 G needle syringe in the following order: 8 mL of the 15% iodixanol solution, 6 mL of the 25% iodixanol solution, 5 mL of the 40% iodixanol solution, and 5 mL of the 60% iodixanol solution. Finally, tubes were filled with the sample, sealed and centrifuged in a Type 70 Ti rotor (Beckman Coulter, California, USA) at 350.000 x g at 10°C for 90 min and then pierced with a 16 G needle on top and an 18 G needle at the interface between the 60% and 40% iodixanol gradients. Viral particles contained in the 40% iodixanol layer were fractioned in 1.5 mL microcentrifuge tubes and concentrated using Amicon Ultra-15 centrifugal filter units (molecular weight cut-off, 100 kDa). Before the concentration step, membranes were activated with 15 mL of 0.1% Pluronic F68 in PBS solution that were discarded and replaced with 15 mL of 0.01% Pluronic F68 in PBS solution. The tubes were centrifuged at 3.000 rpm for 5 min at 4 °C. The supernatant was discarded, and 15 mL of 0.001% Pluronic F68 in PBS + NaCl 200mM solution was added and centrifuged at 3.000 rpm for 5 min at 4°C. The sample was then added and centrifuged at 3.500 rpm for 8 min at 4°C and flowthrough was discarded. During concentration process, formulation buffer (0.001% Pluronic F68 in PBS) was also added to the sample after few centrifugation steps to replace iodixanol and to avoid toxicity in animals after AAV injection. The viral title was quantified using RT-qPCR.

**Immunofluorescence**

Trigeminal nerve tissues and DRG were collected mice deeply anesthetized and transcardially perfused with PBS, were fixed for 24-48 h in 10% formalin, paraffin-embedded and cut with microtome at 5 µm. Tissues were incubated with different primary antibodies: anti-5HT$_{1D}$ (#OSS00185W, Rabbit, 1:1000, Thermo Fisher), anti-5HT$_{1B}$ (#ASR-022, RRID:AB_10561260, Rabbit, 1:100, Alomone labs), anti-DNMT3A (#ab188470, RRID:AB_3073896, Rabbit, 1:500, Abcam), anti-DNMT3B (#ab2851, RRID:AB_303356, Rabbit, 1:500, Abcam), anti-TGF-β3 (#ab15537, RRID:AB_2202305, Rabbit, 1:100 Abcam), anti-TGF-βR1 (#30117-1-AP, RRID:AB_3086235,Rabbit, 1:200, Proteintech), anti-TGF-βR2 (#PA5-35076, RRID:AB_2552386, Rabbit, 1:100, Invitrogen), S100 (#MA1-26621, RRID:AB_2552386Mouse, 1:50, Thermo Fisher) diluted in fresh blocking solution (PBS1X pH 7.4, 5% normal goat serum (NGS) or normal donkey serum (NDS); or PBS1X pH 7.4, 3% Bovine Serum Albumin (BSA) overnight at 4°C. The primary antibody was used after antibody retrieval performed with tri-sodium citrate buffer (pH 6) for TGF-β3, TGF-βR1, TGF-βR2 and for DNMT3B; tris-EDTA buffer (pH 8) for 5HT$_{1B}$ and tris-EDTA (pH 9) for 5HT$_{1D}$ and DNMT3A and following a 1 h incubation with the specific blocking solution. Sections were washed in PBS and then incubated with the appropriate fluorescent polyclonal secondary antibodies Alexa Fluor® 488 (#A32731, goat polyclonal anti-rabbit, 1:600, Invitrogen,), Alexa Fluor® 488 (#A32790, RRID:AB_2633280, donkey polyclonal anti-rabbit, 1:600, Invitrogen), Alexa Fluor® 647 (#A31571, RRID:AB_162542, donkey polyclonal anti-mouse, 1:600, Invitrogen, Carlsbad, California), Alexa Fluor® 647 (#A21236, RRID:AB_2535805,goat polyclonal anti-mouse, 1:600, Invitrogen), Alexa Fluor® 594 (#A11005, RRID:AB_141372,goat polyclonal anti-mouse, 1:600, Invitrogen) for two hours RT. At the end tissues were covered using the mounting medium with DAPI (#ab104139, Abcam) and stored at 4°C utile imaging. Human sciatic nerve (#0062-HP-

261, Gentaur) were incubated with primary antibodies: anti-$5HT_{1D}$ (#OSS00185W, Rabbit, 1:1000, Thermo Fisher), anti-$5HT_{1B}$ (#ASR-022, RRID:AB_2756794, Rabbit, 1:100, Alomone labs,) followed by the same steps of mice tissues. All slides were visualized and analyzed using a Zeiss Axio Imager 2 microscope with Z-stacks in the Apotome mode (Zeiss) or Leica Stellaris 5 confocal microscope (Leica). Pearson correlation (Rcoloc) values in the colocalization analysis were calculated using the colocalization Plugin of the ImageJ (v.1.54 f; National Institutes of Health, Bethesda). We underline, as a limitation of the colocalization method, that detection of the biomarkers localized in two or more different intracellular compartments of the same cell may be affected by the slice orientation that may result in incomplete colocalization.

**Immunocytochemistry**

Human, primary and mouse Schwann cells were fixed in 10% neutral buffered formalin for 10 minutes, rinsed briefly in PBS and permeabilized with 0,5% Triton X-100 for 10 minutes. Cells were treated with a blocking solution 2% BSA in PBS for 1h and were incubated with different primary antibodies: anti-$5HT_{1D}$ (#OSS00185W, Rabbit, 1:100, Thermo Fisher), anti-$5HT_{1B}$ (#ASR-022, RRID:AB_10561260, Rabbit, 1:100, Alomone labs), anti-TGF-β3 (#ab15537, RRID:AB_2202305, Rabbit, 1:100, Abcam), and SOX10 (#AF-2864, RRID:AB_442208, Goat, 1:20, Biotechne) for 1h RT. All incubations were performed on a shaker to enhance the penetration of the solutions. Then cells were incubated with the fluorescent polyclonal secondary antibodies Alexa Fluor® 488 (#A32790, donkey polyclonal anti-rabbit, 1:600, Invitrogen), Alexa Fluor® 594 (#A11058, donkey polyclonal anti-goat, 1:600, Invitrogen) for two hours. At the end cells were covered using the mounting medium with DAPI (#ab104139, Abcam). Cells were imaged on a Leica Stellaris 5 confocal microscope (Leica), and images were processed with ImageJ.

**RNAScope fluorescent in situ hybridization and immunofluorescence**

Fluorescent *in situ* hybridization was performed using the RNAscope™ Multiplex Fluorescent V2 Assay (#323100, ACDbio), according to the manufacturer's protocol. Trigeminal nerves were collected from control and treated animals, fixed for 24–48 h in 10% formalin, paraffin-embedded, and sectioned at 7 µm using a microtome. Slides were pretreated by incubation at 60°C for 1 h on a Thermobrite system, followed by rinsing in xylene and 100% ethanol. RNAscope hydrogen peroxide (#322335; ACDbio) was then applied for 10 min RT to inactivate endogenous peroxidases. Sections were briefly rinsed in distilled water and subjected to antigen retrieval in an oil bath at 99°C for 15 min. After washing in distilled water and 100% ethanol, slides were left to dry overnight RT. The following day, slides were treated with 2–4 drops of Protease Plus (#322331; ACDbio) and incubated at 40°C in a HybEZ II Oven (ACDbio) for 30 min.

Sections were hybridized for 2 h with probes targeting *HTR1B, HTR1D*, and *BETAGLYCAN* in human (#449831, #1113191, #441031; Bio-Techne) and *Htr1b*, *Htr1d, Tgfb3*, and *Betaglycan* in mouse (#315861, #315871, #406211, #406221; Bio-Techne) trigeminal nerves. Signal amplification was performed by sequential incubation with AMP1, AMP2, and AMP3 (#323110; ACDbio) at 40°C. Samples were subsequently incubated with HRP-C1 and the fluorophore TSA-Vivid 650 (#7536; ACDbio) diluted 1:1500 in TSA Buffer (#322810; ACDbio) for 30 min at 40°C in the HybEZ II Oven, followed by incubation with HRP Blocker for 15 min. After the RNAscope procedure, immunohistochemistry was performed. Slides were incubated with 5% normal goat serum (NGS) in 1× PBS containing 0.1% Tween-20 for 1 h RT, followed by incubation with the primary antibody S100 (#MA1-26621, RRID:AB_795376, mouse; 1:50 Thermo Fisher) or NeuN (#MAB377, RRID:AB_2298772 mouse; 1:250) diluted in blocking solution overnight at 4°C. Sections were washed in PBS and incubated with Alexa Fluor® 488 (#A32723, RRID:AB_2633275, goat polyclonal anti-mouse, 1:600, Invitrogen) for 2 h RT. Slides were mounted with DAPI-containing mounting medium (#ab104139, Abcam). Fluorescent images were acquired using a Leica Stellaris 5 (Leica).

### Supplementary Figures

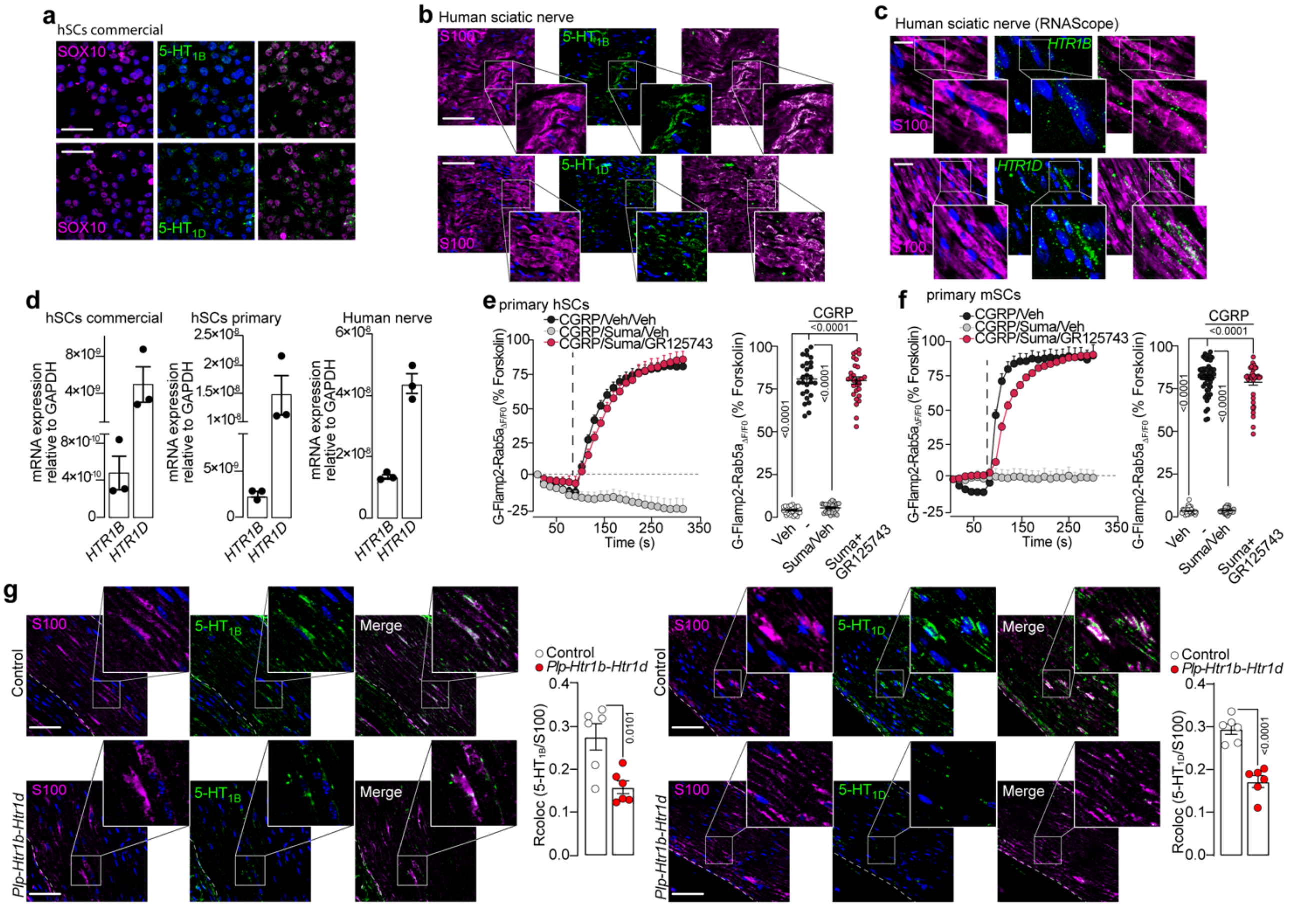


### Figure S1

**(a)** Representative images of 5-HT$_{1B}$ e 5-HT$_{1D}$ expression in human Schwann cells (hSCs, scale bar: 10 µm, n=6) and **(b)** in human sciatic nerve tissues (scale bar: 20 µm, n=6 ). SOX10 and S100 are markers of SCs. **(c)** Representative RNAscope images of *HTR1B* and *HTR1D* mRNA and S100 protein expression in human sciatic nerve (scale bar: 20 µm, n=6). **(d)** RT-qPCR for *HTR1B* and *HTR1D* mRNA in commercial and primary (isolated from lingual, sublingual and inferior alveolar nerves biopsies) hSCs and in human nerve tissues. **(e)** Typical traces and cumulative data of the effects of sumatriptan (Suma, 30 µM) on CGRP (3 µM)-evoked Rab5a cAMP formation in the presence of GR125743 (1 µM) or vehicle (Veh) in primary isolated hSCs (cells number: Veh/Veh/Veh=29, CGRP/Veh/Veh =26, CGRP/Suma/Veh =26, CGRP/Suma/GR125743 =29) and in **(f)** primary mouse SCs (mSCs) (cells number: Veh/Veh/Veh=40, CGRP/Veh/Veh =50, CGRP/Suma/Veh =37,

CGRP/Suma/GR125743 =41). **(g)** Representative images colocalization data (Rcoloc) of 5-$HT_{1B}$ and 5-$HT_{1D}$ silencing in SCs in *Plp-Htr1b-Htr1d* and Control mice. Data are mean ± s.e.m. Student's t-test and 1-way ANOVA, Bonferroni correction.

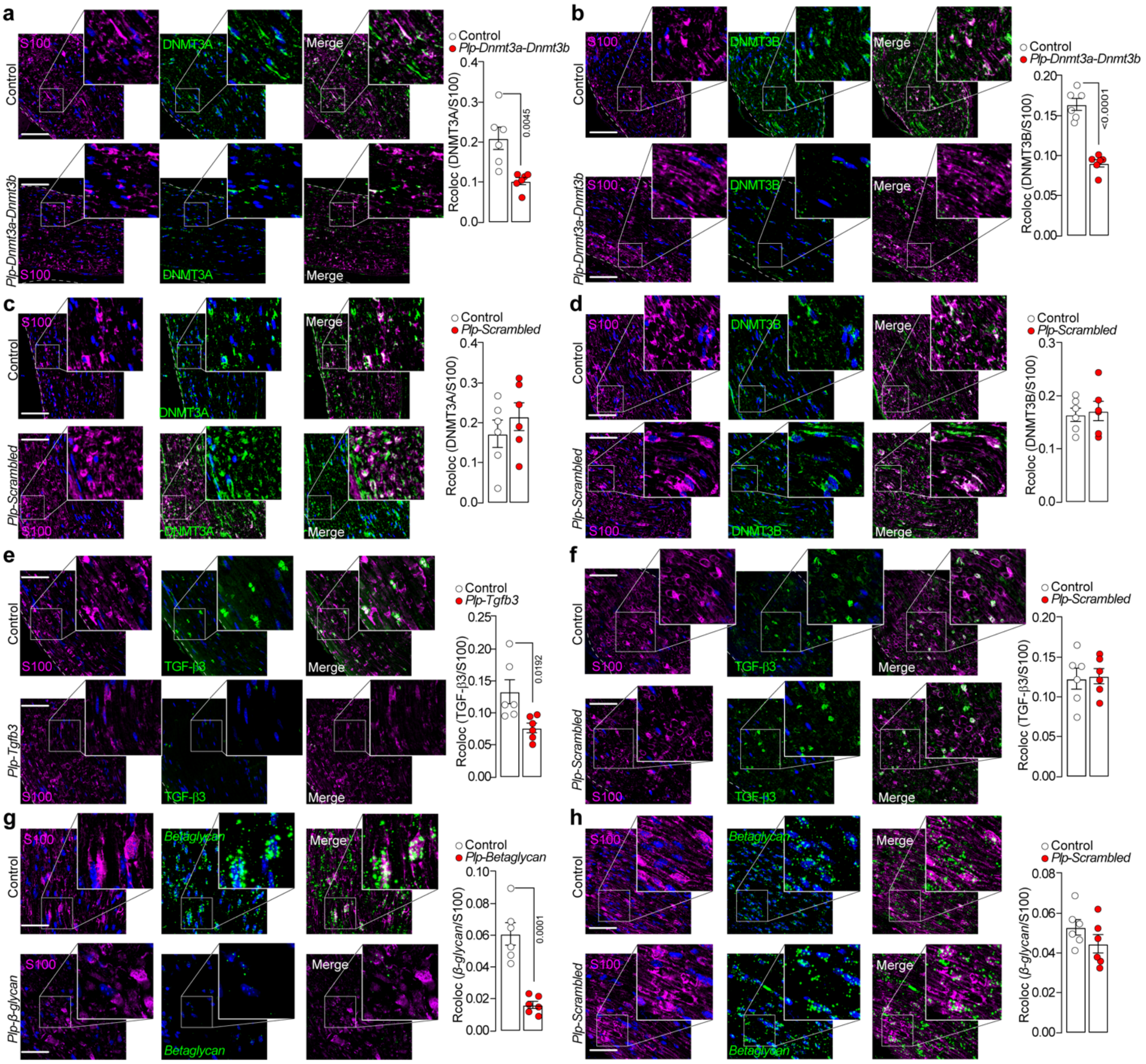


### Figure S2

**(a-d)** Representative images of DNMT3A or DNMT3B and S100 protein expression and colocalization data (Rcoloc) in trigeminal nerve from *Plp-Dnmt3a-Dnmt3b*, *Plp-Scrambled* or Control mice (scale bar: 20 µm, n=6). **(e,f)** Representative images of TGF-β3 and S100 and Rcoloc in trigeminal nerve from *Plp-Tgfb3*, in *Plp-Scrambled* or Control mice (scale bar: 20 µm, n=6). **(g,h)** RNAscope images of *Betaglycan* and S100 and Rcoloc in trigeminal nerve from *Plp-Betaglycan, Plp-Scrambled* or control mice (scale bar: 20 µm, n=6). Data are mean ± s.e.m. (n=6 mice/group). Student's t-test.

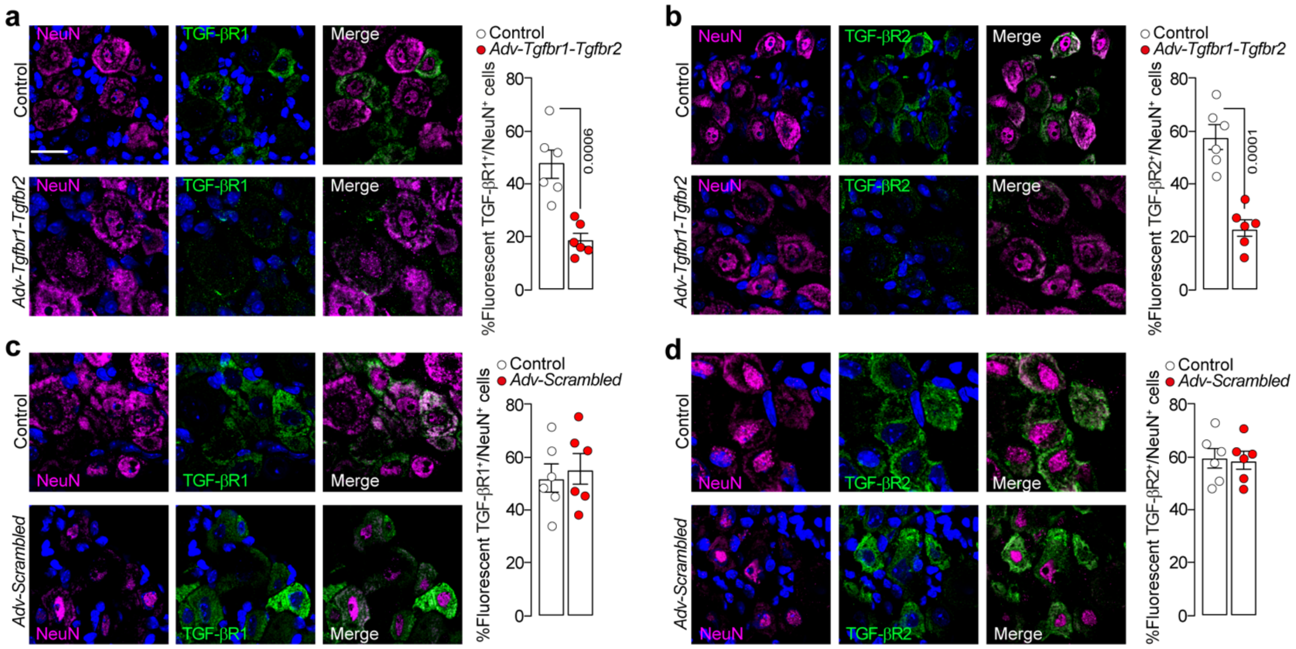


**Figure S3**

Representative images of TGF-βR1 or TGF-βR2 and NeuN protein expression in dorsal root ganglia (DRG) from Control mice and **(a,b)** *Adv-Tgfbr1-Tgfbr2* and **(c,d)** *Adv-Scrambled* mice (scale bar: 20 μm, n=6). Data are mean ± s.e.m. (n=6 mice/group). Student's t-test.

### Supplementary Tables

Table S1: Primers for plasmids construction

| PRIMER | SEQUENCE 5’-3’ |
|---|---|
| P1 | ggcaACCAAGACCCTAGGGATCATTTTACATCTGTGGCTTCACTAAAATGATCCCTAGGGTCTTGGG |
| P2 | agcgCCCAAGACCCTAGGGATCATTTTAGTGAAGCCACAGATGTAAAATGATCCCTAGGGTCTTGGT |
| P3 | ggcaCCGGGTTGATTTGTGCTATCTTTACATCTGTGGCTTCACTAAAGATAGCACAAATCAACCCGT |
| P4 | agcgACGGGTTGATTTGTGCTATCTTTAGTGAAGCCACAGATGTAAAGATAGCACAAATCAACCCGG |
| P5 | ggcaAGCACCTTCCAGGTGGATATAATACATCTGTGGCTTCACTATTATATCCACCTGGAAGGTGCG |
| P6 | agcgCGCACCTTCCAGGTGGATATAATAGTGAAGCCACAGATGTATTATATCCACCTGGAAGGTGCT |
| P7 | ggcaTCCACGAACCTAAGGGTTACTATACATCTGTGGCTTCACTATAGTAACCCTTAGGTTCGTGGG |
| P8 | agcgCCCACGAACCTAAGGGTTACTATAGTGAAGCCACAGATGTATAGTAACCCTTAGGTTCGTGGA |
| P9 | ggcaTCGCTCCGCTGAAGGAATATTTACATCTGTGGCTTCACTAAATATTCCTTCAGCGGAGCGG |
| P10 | agcgCCGCTCCGCTGAAGGAATATTTAGTGAAGCCACAGATGTAAATATTCCTTCAGCGGAGCGA |
| P11 | ggcaTGATGGCTTCAAAGAATGATAATACATCTGTGGCTTCACTATTATCATTCTTTGAAGCCATCG |
| P12 | agcgCGATGGCTTCAAAGAATGATAATAGTGAAGCCACAGATGTATTATCATTCTTTGAAGCCATCA |
| P13 | ggcaGGCTGACAGCTTTGCGAATTAATACATCTGTGGCTTCACTATTAATTCGCAAAGCTGTCAGCT |
| P14 | agcgAGCTGACAGCTTTGCGAATTAATAGTGAAGCCACAGATGTATTAATTCGCAAAGCTGTCAGCC |
| P15 | ggcaCCTCAGGAAATGAGATTGATTTTACATCTGTGGCTTCACTAAAATCAATCTCATTTCCTGAGT |
| P16 | agcgACTCAGGAAATGAGATTGATTTTAGTGAAGCCACAGATGTAAAATCAATCTCATTTCCTGAGG |
| P17 | ggcaGGTCCCTCAAATGCGAAATCTATACATCTGTGGCTTCACTATAGATTTCGCATTTGAGGGACT |
| P18 | agcgAGTCCCTCAAATGCGAAATCTATAGTGAAGCCACAGATGTATAGATTTCGCATTTGAGGGACC |
| P19 | ggcaGGTAATGGCTTCGTCGTTCTTTTACATCTGTGGCTTCACTAAAAGAACGACGAAGCCATTACT |
| P20 | agcgAGTAATGGCTTCGTCGTTCTTTTAGTGAAGCCACAGATGTAAAAGAACGACGAAGCCATTACC |
| P21 | ggcaGTGGATCCTACAACATCAGAGTTACATCTGTGGCTTCACTAACTCTGATGTTGTAGGATCCAT |
| P22 | agcgATGGATCCTACAACATCAGAGTTAGTGAAGCCACAGATGTAACTCTGATGTTGTAGGATCCAC |
| P23 | ggcaGTAATGACCCTGAACACGTACTTACATCTGTGGCTTCACTAAGTACGTGTTCAGGGTCATTAT |
| P24 | agcgATAATGACCCTGAACACGTACTTAGTGAAGCCACAGATGTAAGTACGTGTTCAGGGTCATTAC |
| P25 | ggcaGCTTTCGCGATTCGTCAAATGATACATCTGTGGCTTCACTATCATTTGACGAATCGCGAAAGT |
| P26 | agcgACTTTCGCGATTCGTCAAATGATAGTGAAGCCACAGATGTATCATTTGACGAATCGCGAAAGC |
| P27 | ggcaGGCTTGAAATAGTTACGATAAATACATCTGTGGCTTCACTATTTATCGTAACTATTTCAAGCT |
| P28 | agcgAGCTTGAAATAGTTACGATAAATAGTGAAGCCACAGATGTATTTATCGTAACTATTTCAAGCC |
| P29 | ggcaGTACCTCGTGTCAGAAGATGTATACATCTGTGGCTTCACTATACATCTTCTGACACGAGGTAT |
| P30 | agcgATACCTCGTGTCAGAAGATGTATAGTGAAGCCACAGATGTATACATCTTCTGACACGAGGTAC |
| P31 | ggcaGCAAATAAGTTCTAGCGTTAGTTACATCTGTGGCTTCACTAACTAACGCTAGAACTTATTTGT |
| P32 | agcgACAAATAAGTTCTAGCGTTAGTTAGTGAAGCCACAGATGTAACTAACGCTAGAACTTATTTGC |
| P33 | ATACGAATTCGCCACCATGGGCTCCCACCAC |
| P34 | ACTATCTAGATCATTAGGCTGAGGCAGCTGC |
| P35 | ATACGAATTCGCCACCATGGGCTCCCACCAC |
| P36 | CTTCTGCTAGTGGAGGCTGAGGCAGCTGCTCCTCGT |
| P37 | GCAGCTGCCTCAGCCTCCACTAGCAGAAGCACAGC |
| P38 | TACGGTCGACTCAGTTGCTACAACACTGGC |
| P39 | AAAGAATTCAATGGAGGAACCGGGT |

| | |
|---|---|
| P40 | AAAGGTACCTTAACTTGTGCACTTAAAACG |
| P41 | AAAGAATTCATGTCCCCACTGAAC |
| P42 | AAAGGTACCTTAGGAGGCCTTCC |
| P43 | ATCTGCTAGCATGGAGGAACCGGGTGC |
| P44 | ACCGCTCGATTTACTTGTGCACTTAAAACGTATCAGTTTATGGAATGC |
| P45 | CAAGTAAATCGAGCGGTGGTGGC |
| P45 | GTTCCTCCATGCTAGCAGATCTGGTGGCTTTACC |
| P47 | TGCTAGCATGTCCCCACTGAACCAGT |
| P48 | CCGCTCGATTTGGAGGCCTTCCGGAAAG |
| P49 | CTCCAAATCGAGCGGTGGTGGC |
| P50 | GGGGACATGCTAGCAGATCTGGTGGCT |
| P51 | AGCGAATTCATCCACTAGCAGAAGCAC |
| P52 | AAAGCTAGCTCAGTTGCTACAACACTG |

Table S2: Primer for RT-qPCR

| Gene | Sequence Primer |
|---|---|
| Human GAPDH (NM_002046) | 5'-ACATCGCTCAGACACCATG-3'<br>5'-TGTAGTTGAGGTCAATGAAGGG-3' |
| Human HTR1B (NM_000863) | 5’-TCTTCTCCATCTCTATCTCGCT-3’<br>5’-TGTAGAGGATGTGGTCGGT-3’ |
| Human HTR1D (NM_000864) | 5’-TCCTGCATCTCTGTGTCATTG-3’<br>5’-GTCCTGCGTTTACTGTATTCCA-3’ |
| Human TGFB3 (NM_003239) | 5'-GAGTGGCTGTCCTTTGATGT-3'<br>5'-AGTGAATGCTGATTTCTAGACCT-3' |
| Human TGFBR3 (NM_036634) | 5'-ATCACCTTCAACATGGAGCTAT-3'<br>5'-CCCAGTTCTTGTTCAGCCTTA-3' |
| Mouse Gapdh (NM_008084) | 5'-AATGGTGAAGGTCGGTGTG-3'<br>5'-GTGGAGTCATACTGGAACATGTAG-3' |
| Mouse Htr1b (NM_010482) | 5'-ACCCTAGGGATCATTTAGGAGC-3'<br>5'-ATGAGGGAGTTAAGATAGCCTAACC-3' |
| Mouse Htr1d (NM_008309) | 5'-GGATGCTGGTGATAACAAGACA-3'<br>5'-GTGACCAAGACTCAAAGAATGC-3' |
| Mouse Tgfb3 (NM_009368) | 5'-GCCAAAGAGATCCATAAATTCGAC-3'<br>5'-ACTGAGGACACATTGAAACGA-3' |
| Mouse Tgfbr3 (NM_011578) | 5'-GCATGTGAAGACAGAGAGACT-3'<br>5'-CTCCTTTCCTCTGTTTCTGCT-3' |
| Mouse Sox11 (NM_009234) | 5’-CAATGAGATACGAGTGGAGAGG-3’<br>5’-CCATCAAAGAAACCAAAGCCAT-3’ |
| Mouse Fosl2 (NM_008037) | 5’-CAGCCAAGTGTCGGAACC-3’<br>5'-CTGCAGCTCAGCAATCTCTT-3’ |
| Mouse Klf6 (NM_011803) | 5’-TCCCAATGTGTAGCATCTTCC-3’<br>5’-AAGATAGCGTTCCAACTCCAG-3’ |

| | |
|---|---|
| Mouse Arid5a (NR_033310) | 5'-CACCCAGCTCCAGTTAACC-3'<br>5'-GTTCATCATACACGTTCTTCCAG-3' |
| Mouse Atf4 (NM_009716) | 5'-CGTATTAGAGGCAGCAGTGC-3'<br>5'-AGGTATCTTTGTCCGTTACAGC-3' |
| Mouse Atf3 (NM_007498) | 5'-AGCTGAGATTCGCCATCCAGAA-3'<br>5'-CTCGCCGCCTCCTTTTCCT-3' |
| Mouse Onecut2 (NM_194268) | 5'-GAGTTCCAGCGCATGTCT-3'<br>5'-TTCTTCTGCGAGTTGTTCCTG-3' |

Table S3: Heatmap of differentially expressed genes DEGs in hSCs.

| ENST_ID | Gene_Symbol | ENST_ID | Gene_Symbol |
|---|---|---|---|
| ENST00000607092 | CUX1 | ENST00000464238 | MIR9-1HG |
| ENST00000287771 | RBPMS | ENST00000560549 | TMOD3 |
| ENST00000509007 | THAP9-AS1 | ENST00000292530 | ZNF333 |
| ENST00000504869 | THAP9-AS1 | ENST00000604886 | ZNF559-ZNF177 |
| ENST00000221554 | YJU2B | ENST00000382905 | ZMYM5 |
| ENST00000593174 | YJU2B | ENST00000467542 | ZMYM5 |
| ENST00000569540 | TMEM170A | ENST00000562413 | |
| ENST00000518808 | XPO7 | ENST00000524224 | CCNE2 |
| ENST00000508529 | TUBBP5 | ENST00000567632 | NA |
| ENST00000543687 | ATP6V0A2 | ENST00000517465 | UBR5 |
| ENST00000354638 | OCA2 | ENST00000482921 | ASB13 |
| ENST00000580288 | STRADA | ENST00000338523 | SNX10 |
| ENST00000591921 | MAP4K1 | ENST00000601142 | ZNF30 |
| ENST00000591748 | IGFLR1 | ENST00000254250 | THAP1 |
| ENST00000397512 | TMEM80 | ENST00000545654 | FAM76B |
| ENST00000269724 | SAMD1 | ENST00000524469 | AHI1 |
| ENST00000367433 | KCNT2 | ENST00000601648 | MUC20-OT1 |
| ENST00000237536 | MTCL2 | ENST00000553234 | COQ10A |
| ENST00000531108 | | ENST00000579124 | RBBP8 |
| ENST00000565951 | CPNE2 | ENST00000454300 | NA |
| ENST00000511022 | PPIP5K2 | ENST00000518914 | FAM114A2 |
| ENST00000543863 | LTO1 | ENST00000479608 | ARID2 |
| ENST00000534562 | SWAP70 | ENST00000594075 | TUBB4A |
| ENST00000395002 | FAM13A | ENST00000369559 | HIPK1 |
| ENST00000435940 | CENPPP1 | ENST00000361786 | CIPC |
| ENST00000593228 | HEATR6 | ENST00000461646 | DLG3 |
| ENST00000572757 | NAA60 | ENST00000397578 | RESF1 |
| ENST00000576468 | OR1F1 | ENST00000459986 | MYH10 |
| ENST00000509560 | WDR19 | ENST00000235150 | RNF19B |
| ENST00000311417 | ZMAT3 | ENST00000507667 | ELMOD2 |
| ENST00000398578 | ZC3H11C | ENST00000533112 | MYO18A |
| ENST00000306481 | CEP120 | ENST00000477719 | ABCB8 |
| ENST00000541318 | KDM2B | ENST00000523569 | CHRAC1 |
| ENST00000556663 | GPATCH2L | ENST00000514821 | CIR1P2 |
| ENST00000383926 | NA | ENST00000475981 | MLF1-DT |
| ENST00000286181 | HYCC2 | ENST00000453131 | APOD |

| | | | |
|---|---|---|---|
| ENST00000535007 | STK25 | ENST00000521620 | WRN |
| ENST00000427980 | ATXN1L | ENST00000497267 | ARHGEF3 |
| ENST00000413791 | PCBP1-AS1 | ENST00000453074 | TRMT6 |
| ENST00000589740 | HEATR6-DT | ENST00000574954 | ARRB2 |
| ENST00000608174 | TMEM80 | ENST00000420891 | ZNF620 |
| ENST00000591517 | | ENST00000542012 | NA |
| ENST00000396857 | MAP4K1 | ENST00000560185 | AKAP13 |
| ENST00000561911 | SPG7 | ENST00000356929 | ZNF708 |
| ENST00000577978 | DDX42 | ENST00000384627 | RNU6-2 |
| ENST00000331758 | SS18L1 | ENST00000424010 | CT75 |
| ENST00000592969 | NA | ENST00000429475 | CT75 |
| ENST00000461444 | STXBP4 | ENST00000446532 | CT75 |
| ENST00000496626 | CREM | ENST00000372172 | HECTD3 |
| ENST00000512741 | AP1AR | ENST00000273580 | NA |
| ENST00000458537 | SKIL | ENST00000554938 | LIN52 |
| ENST00000421413 | EYA4 | ENST00000394520 | RNF14 |
| ENST00000588267 | ZNF846 | ENST00000464425 | VPS52 |
| ENST00000341491 | ATF3 | ENST00000492047 | WBP1 |
| ENST00000409009 | NA | ENST00000438374 | LRRFIP2 |
| ENST00000476559 | RPGR | ENST00000566042 | CPNE2 |
| ENST00000463509 | RBFOX2 | ENST00000584754 | PPP4R1 |
| ENST00000456735 | NF1 | ENST00000595324 | TUBB4A |
| ENST00000509707 | SPINK2 | ENST00000601152 | TUBB4A |
| ENST00000588061 | TNRC6C | ENST00000585155 | CLUL1 |
| ENST00000527374 | | ENST00000399100 | SEC24B |
| ENST00000380692 | MSL3 | ENST00000594208 | KPTN |
| ENST00000603726 | DUSP22 | ENST00000536339 | NA |
| ENST00000554660 | VPS33B | ENST00000579843 | TRIM16 |
| ENST00000254488 | SLC6A11 | ENST00000494872 | TRIM2 |
| ENST00000550223 | LINC01234 | ENST00000519553 | PTPN12 |
| ENST00000590806 | DNM2 | ENST00000496969 | SF1 |
| ENST00000523642 | NDRG1 | ENST00000376956 | ZFAND5 |
| ENST00000586018 | CILP2 | ENST00000570092 | RHOT2 |
| ENST00000291495 | CILP2 | ENST00000505634 | DCUN1D4 |
| ENST00000378883 | MCM8 | ENST00000478966 | SERPINE2 |
| ENST00000352210 | PPOX | ENST00000446263 | MTND4LP1 |
| ENST00000367999 | PPOX | ENST00000428103 | NA |
| ENST00000331426 | RBM43 | ENST00000458525 | |
| ENST00000475249 | PPP1R1C | ENST00000541054 | NA |
| ENST00000453212 | MATCAP2 | ENST00000590706 | LIN37 |
| ENST00000541339 | GALNT8 | ENST00000519264 | ASPH |
| ENST00000477754 | ARHGEF2 | ENST00000303586 | ZNF30 |
| ENST00000441123 | BRPF3 | ENST00000439785 | ZNF30 |
| ENST00000381975 | TMEM44 | ENST00000464934 | FANCD2 |
| ENST00000347147 | TMEM44 | ENST00000471564 | STRBP |
| ENST00000270583 | KLHDC4 | ENST00000372427 | HYI |
| ENST00000433861 | BRPF1 | ENST00000325089 | SLITRK5 |
| ENST00000369158 | H3C14 | ENST00000460912 | SYNE1 |
| ENST00000498534 | CNOT4 | ENST00000538413 | LRRC51 |
| ENST00000560928 | CD276 | ENST00000559040 | GALK2 |
| ENST00000343267 | APOD | ENST00000433855 | NA |
| ENST00000458447 | APOD | ENST00000465307 | PGAP2 |

| | | | |
|---|---|---|---|
| ENST00000254262 | FAAP24 | ENST00000585953 | RPRD1A |
| ENST00000483649 | ASAH2B | ENST00000566662 | WWOX |
| ENST00000543840 | NA | ENST00000474353 | SORBS1 |
| ENST00000056233 | NFE2L3 | ENST00000502931 | STARD4 |
| ENST00000536790 | ZNF140 | ENST00000324794 | PIAS2 |
| ENST00000462549 | RABGGTBP1 | ENST00000315249 | RFFL |
| ENST00000554055 | | ENST00000466189 | PHC3 |
| ENST00000527788 | QSER1 | ENST00000526526 | KIAA1191P2 |
| ENST00000392432 | TMEM44 | ENST00000439000 | MYADM |
| ENST00000594276 | TUBB4A | ENST00000451655 | BACH1 |
| ENST00000519301 | NRG1 | ENST00000338754 | TPST2 |
| ENST00000496838 | TSC22D1 | ENST00000520407 | NRG1 |
| ENST00000461013 | SLC22A5 | | |

Table S4: List of differentially expressed genes at 6h, 24h and 48h of Sumatriptan stimulation

| 6 hours stimulation | | |
|---|---|---|
| Gene | log2FoldChange | padj |
| PDIA3P1 | 0,593 | 0,00709658623068178 |
| RP11-364L4.1 | 0,635 | 0,000393460243944849 |
| FTH1P8 | 0,598 | 0,000161148093271818 |
| MRPL41 | 0,701 | 0,000949435374337061 |
| COX5A | 0,884 | 3,38746287369936E-07 |
| GCSH | 0,625 | 0,000161148093271818 |
| ENO3 | 0,626 | 0,000161148093271818 |

| 24 hours stimulation | | | | | | | | |
|---|---|---|---|---|---|---|---|---|
| Gene | log2FoldChange | padj | Gene | log2FoldChange | padj | Gene | log2FoldChange | padj |
| DDX11L1 | -1,930 | 1,74902823756886E-06 | RP3-426I6.2 | 9,274 | 2,93673942200055E-33 | ANAPC10P1 | 8,061 | 6,15194034328835E-14 |
| AL627309.1 | -2,979 | 0,000383481263806135 | STX12 | 1,067 | 2,24582647630993E-08 | PLA2G12AP1 | 4,236 | 5,742917843209E-08 |
| RP11-34P13.15 | -3,501 | 0,00212671786805353 | PPP1R8 | -0,747 | 0,00103826528250255 | ORC1 | 1,302 | 1,54549049622084E-06 |
| AP006222.2 | 2,008 | 9,54090012359026E-20 | SMPDL3B | 2,324 | 5,40947931565244E-22 | PRPF38A | -1,775 | 2,88663639599155E-28 |
| AP006222.1 | 2,198 | 3,1186760297904E-22 | RP11-460I13.2 | 0,780 | 0,00455135309342556 | COA7 | 1,012 | 1,08889125731074E-10 |
| RP4-669L17.10 | -1,844 | 0,000422450421090022 | RP5-1053E7.3 | 6,585 | 4,15127008474614E-09 | NDUFS5P3 | 6,338 | 7,0685520844312E-227 |
| RP4-669L17.8 | -1,969 | 0,0072640557042167 | DNAJC8 | -0,674 | 9,44295787387074E-08 | RP4-631H13.6 | -1,560 | 3,36357513144842E-27 |
| MTND1P23 | -2,149 | 1,8734977034992E-15 | AL353354.1 | -0,612 | 1,75106061458481E-06 | RP4-631H13.4 | 5,631 | 0,000102805723651797 |
| MTND2P28 | -2,791 | 1,2283922790887E-22 | AL353354.2 | -0,745 | 2,2101516989741E-09 | SCP2 | 1,589 | 2,13064518079349E-43 |
| hsa-mir-6723 | -2,190 | 2,32173995159995E-19 | ATPIF1 | 1,043 | 4,12400919695532E-11 | CPT2 | -1,126 | 5,3185877736841E-10 |
| RP5-857K21.7 | -2,177 | 6,15833348074723E-20 | RP1-308E4.1 | 4,348 | 7,66565619275592E-07 | RP5-1024G6.2 | -1,714 | 0,000890959096642097 |
| MTATP8P1 | -1,782 | 5,65944078043701E-09 | SNHG3 | -0,806 | 0,000600373331749548 | NDC1 | 1,970 | 7,70739258786802E-25 |
| MTATP6P1 | -1,984 | 2,74849798950211E-12 | PRDX3P2 | 9,452 | 1,79201282176127E-45 | SNORA58 | -2,913 | 2,10284752147255E-37 |
| RP5-857K21.11 | -1,993 | 9,77163350581376E-13 | TRNAU1AP | 1,191 | 5,76721050589722E-18 | YIPF1 | 2,500 | 2,25376337906298E-30 |
| RP11-206L10.1 | -2,228 | 0,00489620461989903 | TAF12 | 1,622 | 7,52773485047438E-14 | HSPB11 | 1,287 | 3,91189645605204E-19 |
| RP11-206L10.8 | 2,623 | 5,11824137388815E-06 | GMEB1 | 2,575 | 6,51482990128008E-18 | LRRC42 | 0,593 | 0,000563318743777319 |
| LINC01128 | 0,795 | 0,00572099216125424 | YTHDF2 | 1,657 | 2,06100343642365E-20 | TMEM59 | 2,829 | 1,48756583081889E-130 |
| SAMD11 | -1,286 | 0,000481105070667523 | RP4-604A21.1 | 5,474 | 1,92513875040935E-170 | TCEANC2 | 1,134 | 3,64272516098903E-05 |
| NOC2L | -1,280 | 0,000170102606130789 | TMEM200B | -2,700 | 1,15595459076613E-05 | RP4-758J24.4 | 8,018 | 1,92464350439667E-184 |
| RP11-54O7.17 | -4,042 | 0,00447505772864628 | SRSF4 | 1,806 | 1,73979631010117E-08 | RP11-446E24.4 | 1,013 | 0,00453337573753484 |
| ISG15 | 2,262 | 0,00776272528765196 | MECR | 2,434 | 1,98783592086298E-72 | AL357673.1 | -3,603 | 0,00528689697593198 |
| C1orf159 | 2,635 | 0,00990208378623062 | PTPRU | -4,316 | 2,09678486129124E-05 | CYB5RL | 1,621 | 3,40455319666928E-08 |
| SDF4 | 0,744 | 0,000866177962634915 | SDC3 | -4,643 | 0,000242184017258713 | MRPL37 | 1,814 | 1,26350106657308E-49 |

| | | | | | | | | |
|---|---|---|---|---|---|---|---|---|
| B3GALT6 | -2,454 | 3,86559249111093E-05 | SEPW1P | 6,208 | 2,19934548568985E-06 | SSBP3 | -1,746 | 0,000113927870119946 |
| UBE2J2 | 2,476 | 7,66344714136968E-12 | SNRNP40 | -2,489 | 1,27635406860483E-38 | MROH7-TTC4 | -0,683 | 0,000815336964760056 |
| PUSL1 | 0,760 | 0,00860962790012483 | RP11-490K7.4 | 9,255 | 1,30782202482341E-17 | TTC4 | -0,597 | 0,00424735059808877 |
| CPSF3L | 1,824 | 1,33273661549703E-19 | ZCCHC17 | 1,860 | 1,24118874741505E-30 | GYG1P3 | 9,310 | 6,28780895490831E-18 |
| TAS1R3 | -2,359 | 0,00245056371999418 | SERINC2 | 2,518 | 0,00011556666515294 | RP11-377K22.2 | -2,839 | 1,70576742982677E-32 |
| DVL1 | -1,937 | 0,0049432689377459 | HCRTR1 | -1,337 | 6,48163448771321E-10 | DAB1 | 0,994 | 0,000955744550841443 |
| RP4-758J18.2 | -0,732 | 0,000150712999628393 | PEF1 | 1,100 | 3,54372636923761E-12 | RPS26P15 | -1,093 | 1,32521629255584E-09 |
| MRPL20 | 0,678 | 0,00063055246101383 | RP11-73M7.6 | -3,251 | 1,1615221884768E-34 | OMA1 | 2,611 | 6,71383984160353E-31 |
| SSU72 | 1,071 | 3,64584303736447E-11 | RP11-73M7.9 | -3,266 | 5,01340620808207E-34 | RP4-592A1.2 | 2,193 | 3,84599180894164E-36 |
| C1orf233 | -3,826 | 0,000185329703035807 | PTP4A2 | -1,704 | 2,43010460706526E-12 | MYSM1 | 1,034 | 0,00141776045544811 |
| RP11-345P4.10 | -2,054 | 0,000108853567403373 | KHDRBS1 | -1,875 | 1,35173709766585E-17 | JUN | -3,207 | 7,08091779953037E-19 |
| SLC35E2B | -1,069 | 0,00098593254899608 | TMEM39B | 1,555 | 1,56565742195151E-13 | PHBP3 | 9,769 | 3,33104081185821E-118 |
| GNB1 | -1,385 | 3,60455328168947E-13 | RP4-622L5.2 | -1,904 | 3,17962355247373E-10 | FGGY | 3,388 | 8,35977718175299E-17 |
| PRKCZ | 2,936 | 0,00176749179103229 | TXLNA | -0,952 | 0,00405485205691397 | CYP2J2 | 2,398 | 8,91672199315301E-17 |
| SKI | -2,441 | 7,40354334513958E-06 | CCDC28B | 1,195 | 4,86757221671024E-18 | TM2D1 | 1,951 | 2,71194008395211E-44 |
| RER1 | 1,342 | 2,11227531631146E-21 | RP4-622L5.7 | -2,008 | 2,69216611173893E-30 | LAMTOR5P1 | 9,716 | 2,5761289653979E-38 |
| PANK4 | 2,917 | 8,71294208822522E-13 | TMEM234 | 1,814 | 1,48256334796911E-16 | USP1 | -1,649 | 1,7221448362769E-09 |
| WRAP73 | 3,559 | 5,64528623951383E-58 | EIF3I | 0,956 | 3,65715332007497E-12 | ATG4C | 2,567 | 4,87007492500376E-49 |
| TP73-AS1 | -2,407 | 3,39832157568161E-05 | HDAC1 | 1,601 | 5,9289927413484E-83 | RP4-792G4.3 | 10,008 | 7,51371570929137E-21 |
| LRRC47 | -2,789 | 1,23104329306843E-07 | MARCKSL1 | -1,974 | 4,06775394670709E-22 | ALG6 | 2,809 | 4,37168546562409E-94 |
| DFFB | 1,294 | 0,00769494746917844 | BSDC1 | 1,236 | 2,78790389030462E-09 | ITGB3BP | 1,581 | 1,10608489473086E-36 |
| C1orf174 | 0,769 | 1,41830750350164E-06 | ZBTB8OS | -1,002 | 2,75363008753577E-09 | EFCAB7 | 2,635 | 1,6179734928063E-23 |
| RPL22 | -4,013 | 1,8976918912102E-159 | RBBP4 | -3,935 | 1,00956387523473E-104 | DLEU2L | 2,842 | 0,000429498219934734 |
| RNF207 | -2,869 | 2,14409139350651E-17 | SYNC | -4,080 | 2,95556377092826E-91 | PGM1 | 2,546 | 5,59204466426148E-16 |
| ICMT | -1,401 | 1,78191358444374E-07 | KIAA1522 | -2,292 | 8,39841633414774E-06 | RAVER2 | -1,912 | 1,11220037551737E-07 |
| KLHL21 | -1,890 | 0,00152803301012107 | YARS | 1,456 | 8,32584875607254E-32 | RP4-535B20.2 | 4,010 | 3,50219170156465E-05 |
| PHF13 | -2,091 | 0,00810948459361248 | S100PBP | 1,043 | 0,00726617809653138 | MRPS21P1 | 7,630 | 7,08053310269565E-11 |
| THAP3 | 2,010 | 1,1613255748394E-17 | RNF19B | -4,533 | 0,000120239824022521 | AK4 | -1,989 | 1,34905501361716E-11 |
| DNAJC11 | 2,410 | 9,2667293621578E-36 | RP1-117O3.2 | -3,016 | 0,00379770214452853 | RP4-700A9.1 | 3,914 | 3,26895111993394E-13 |
| snoU13 | -1,104 | 1,40606310114185E-05 | ZNF362 | -2,763 | 0,00404551977559472 | DNAJC6 | -3,028 | 0,00742738908883974 |
| CAMTA1 | 1,157 | 5,26503631457044E-23 | AL513327.1 | -2,666 | 0,00628884279615162 | RP11-430H12.2 | 5,773 | 4,191284552324E-06 |
| PARK7 | 1,175 | 1,51076183305777E-17 | PHC2 | -1,381 | 0,000196778985979793 | MIER1 | 1,479 | 1,0481296194808E-10 |
| RERE | -2,352 | 5,88657230897045E-06 | RP4-580O19.2 | 12,743 | 1,18227727765966E-33 | SERBP1 | -1,677 | 4,09195690200978E-43 |
| RPL7P7 | -4,811 | 0,00040269356069592 | RP11-244H3.1 | -0,593 | 0,00236197209904385 | GNG12-AS1 | -1,313 | 0,000230919520525492 |
| ENO1 | -0,951 | 5,69880822365113E-09 | ZMYM6NB | 1,008 | 1,99493571087113E-10 | RP4-609E1.2 | 6,312 | 1,93020161402461E-06 |
| HMGN2P17 | -0,936 | 3,8476311376477E-06 | SFPQ | -1,012 | 4,44330686789113E-05 | WLS | 1,902 | 2,29065860605751E-62 |
| SLC25A33 | -2,904 | 6,15613653097387E-05 | RPL5P4 | 3,888 | 8,11131752233816E-118 | CTBP2P8 | 2,599 | 2,76110457354722E-06 |
| PIK3CD | -3,006 | 3,01526781680313E-08 | NCDN | -1,209 | 0,00466152270502122 | RPS7P4 | -1,454 | 6,11510647726733E-21 |
| CLSTN1 | -2,150 | 0,000164230751294359 | RP4-728D4.2 | 2,163 | 0,00627039662846302 | TCEB1P18 | 5,541 | 0,000154838880620694 |
| CTNNBIP1 | -2,633 | 0,000310388982241247 | PSMB2 | 1,956 | 1,91962888842086E-44 | RP5-1033K19.2 | 8,856 | 9,17611921816271E-16 |
| RP11-84A14.5 | -3,198 | 0,000181440506169627 | AGO4 | -2,352 | 0,00778176165868827 | DEPDC1 | -1,785 | 4,17050378504749E-11 |
| LZIC | 2,037 | 8,46507766148868E-20 | AGO1 | -1,980 | 1,71684102471346E-07 | RP4-694A7.2 | -5,598 | 4,41826107906515E-07 |
| KIF1B | -2,558 | 2,0294914382995E-07 | TRAPPC3 | 1,147 | 3,12523948266649E-16 | RP4-694A7.4 | -4,160 | 0,00789018003102611 |
| PGD | -2,198 | 6,8363895639433E-33 | MAP7D1 | -2,561 | 1,43284153742111E-07 | PIN1P1 | 6,618 | 4,9647044669736E-09 |
| APITD1 | 1,535 | 2,63443713578455E-20 | THRAP3 | -1,038 | 0,000478904219723899 | LRRC40 | 2,257 | 3,17707698436343E-23 |
| APITD1-CORT | 1,584 | 8,21826502353943E-22 | LSM10 | 0,866 | 3,0444925627048E-12 | CTH | 1,751 | 5,29906368245695E-09 |
| DFFA | 1,460 | 4,26488187661364E-25 | OSCP1 | 1,123 | 0,000391947081611556 | RP11-42O15.2 | 4,509 | 3,99249544843334E-28 |
| SRM | -0,954 | 0,000402378900743588 | SNORA63 | -2,143 | 0,00934272677491001 | CASP3P1 | 6,987 | 9,23595686319922E-09 |
| EXOSC10 | 1,966 | 4,51563081135253E-46 | MRPS15 | 1,738 | 3,22833171800773E-52 | ZRANB2-AS1 | -1,202 | 2,37969477862164E-10 |
| RP4-635E18.6 | -1,011 | 0,000230637857878506 | MEAF6 | 1,956 | 1,58335772811198E-56 | ZRANB2 | 1,468 | 2,95479084195857E-27 |
| MTOR | -1,636 | 0,000229604761032807 | FTH1P1 | -2,721 | 0,0065070004230249 | GDI2P2 | 9,710 | 1,27516472575083E-19 |
| UBE2V2P3 | 5,075 | 0,000222657855884109 | RP3-423B22.5 | -1,430 | 2,94937857244246E-25 | RPL31P12 | -1,725 | 5,16945391968217E-11 |
| FBXO44 | 1,183 | 0,000118515181618774 | DNALI1 | -3,537 | 1,10040076113054E-56 | LRRIQ3 | 0,939 | 0,00118114785960026 |
| FBXO6 | 1,311 | 3,89002627248997E-05 | C1orf109 | 0,691 | 0,00405680705869803 | CRYZ | 0,860 | 2,18138479617803E-08 |
| MAD2L2 | 1,477 | 2,58418543652708E-23 | CDCA8 | 1,417 | 2,29644326901921E-13 | TYW3 | 1,439 | 8,58845998574376E-24 |
| AGTRAP | 1,553 | 4,16601414881988E-31 | MANEAL | -1,804 | 0,0030054746903095 | ACADM | 0,664 | 1,11804694130239E-08 |

| | | | | | | | | |
|---|---|---|---|---|---|---|---|---|
| KIAA2013 | -2,608 | 8,32932512915574E-07 | RP11-109P14.9 | -2,400 | 0,000312884856888379 | RABGGTB | 1,421 | 2,16745565455358E-39 |
| PLOD1 | 2,846 | 3,54892559485005E-26 | YRDC | -1,973 | 2,7694014749621E-08 | PIGK | 2,105 | 2,40278857255881E-47 |
| RP5-1077B9.5 | 6,713 | 9,72014105581867E-08 | C1orf122 | -1,430 | 7,94719459982986E-08 | RP11-363H12.1 | 5,892 | 4,48060570377277E-22 |
| MIIP | 2,982 | 4,56256277923753E-10 | RP11-109P14.10 | -2,051 | 0,00182251336859949 | USP33 | 1,332 | 0,000990580160681228 |
| SNORA70 | -1,668 | 9,04119774469546E-05 | INPP5B | 1,466 | 0,00299167405852646 | FAM73A | 1,926 | 0,00708556041324074 |
| VPS13D | -2,687 | 2,16435173133257E-05 | SF3A3 | -2,820 | 6,8810209579811E-59 | NSRP1P1 | 2,620 | 2,00502026889965E-07 |
| DHRS3 | 1,518 | 0,000565694015107408 | UTP11L | 1,623 | 3,79013019557859E-33 | NEXN | 1,616 | 8,12538309526753E-07 |
| KAZN | -3,523 | 0,00643263076243768 | RP5-884C9.3 | 8,035 | 6,72771246176402E-12 | FUBP1 | 0,585 | 0,000117013133435027 |
| CASP9 | 1,298 | 1,80189546346153E-06 | RRAGC | -2,193 | 5,86774810307058E-06 | RP11-386I14.2 | 11,037 | 4,44361951115381E-25 |
| CHCHD2P6 | 0,991 | 3,66822577163694E-16 | MACF1 | -1,966 | 0,000201043859585583 | RP11-386I14.3 | 8,176 | 7,36731391155347E-68 |
| DDI2 | -1,712 | 0,00260753972870842 | PABPC4 | -1,824 | 7,79932514365128E-15 | RNFT1P2 | 5,257 | 0,000673663360267607 |
| PLEKHM2 | -2,286 | 0,00502860059015895 | RP11-69E11.8 | -3,777 | 4,5522833525672E-20 | RP11-472F19.1 | 3,279 | 0,000328005595688243 |
| RP11-169K16.7 | -0,599 | 6,40588502662516E-05 | PPIE | 2,253 | 2,79133541912137E-83 | RP4-641G12.4 | 10,138 | 3,96925547904642E-21 |
| RP11-169K16.8 | 4,091 | 2,34981647213297E-185 | RP1-144F13.4 | 4,274 | 1,25980061178505E-09 | PSAT1P3 | 7,004 | 8,29061669740568E-33 |
| RP11-169K16.9 | -1,674 | 7,6293259344777E-07 | BMP8B | -1,335 | 0,000602960845966412 | RP3-445O10.1 | -2,447 | 2,52824612784877E-09 |
| SPEN | -2,209 | 2,42511709128979E-12 | TRIT1 | 1,546 | 5,06320161010177E-18 | RP5-837I24.5 | -3,908 | 1,6821524858763E-11 |
| MT1XP1 | 2,256 | 4,31531472192038E-29 | CAP1 | -0,788 | 2,76681882406189E-05 | RP11-486G15.1 | 8,928 | 4,32429660506871E-16 |
| SZRD1 | 0,704 | 8,47833282689936E-06 | PPT1 | 1,860 | 4,44164370722172E-53 | RP11-82H13.2 | 4,939 | 5,84150838322091E-113 |
| RP11-430L17.1 | 3,927 | 1,494190280549E-14 | RP11-115D7.3 | 8,194 | 6,34081083944672E-13 | SAMD13 | 2,145 | 0,00459637725780951 |
| NECAP2 | 2,245 | 4,69823268477509E-27 | RP1-39G22.7 | -1,568 | 1,32020864795624E-05 | RPF1 | 1,315 | 1,66031907666932E-19 |
| RNU1-1 | -2,298 | 7,21170453300372E-10 | ZMPSTE24 | 1,593 | 2,32483571190541E-13 | GNG5 | -1,172 | 1,08301130815268E-20 |
| NBPF1 | -1,288 | 0,00153332774563552 | RP1-39G22.4 | 3,689 | 3,83124246029287E-06 | MCOLN3 | 1,189 | 0,000944311145021819 |
| RNU1-3 | -2,354 | 9,93219530381162E-11 | RP1-228H13.1 | 2,009 | 1,27324216838623E-12 | C1orf52 | 1,067 | 5,22774314784165E-10 |
| RNU1-4 | -2,339 | 1,38035422586573E-09 | ZNF684 | 0,679 | 0,00840342031991496 | ODF2L | 0,954 | 0,00328983028058372 |
| CROCCP4 | -2,794 | 0,00590575284357184 | NFYC-AS1 | -1,323 | 0,000590923142061698 | SH3GLB1 | 1,743 | 1,30992884343912E-28 |
| RNU1-2 | -2,490 | 8,05385105445559E-11 | NFYC | 0,790 | 0,00140081273595909 | SEP15 | 1,131 | 7,73226450277027E-24 |
| MFAP2 | 3,238 | 8,55764190245589E-27 | CTPS1 | 1,951 | 3,24595684862547E-30 | RP5-1052I5.2 | 1,468 | 0,000494872403190196 |
| SDHB | 2,697 | 4,1386724081903E-102 | SCMH1 | 1,273 | 8,70627513829877E-05 | RP11-384B12.2 | 6,270 | 2,34434358365909E-10 |
| RP1-20B21.4 | -2,848 | 2,27108228242338E-34 | TMSB4XP1 | -4,125 | 5,70094433861475E-12 | RP11-384B12.3 | 7,781 | 7,80707285377649E-12 |
| RCC2 | -2,579 | 2,48369386763901E-35 | PPIH | -3,672 | 7,08553523153267E-99 | GTF2B | 1,818 | 6,17372015062082E-25 |
| AC004824.1 | -3,749 | 0,00757542235715795 | YBX1 | -3,001 | 4,80005306620025E-57 | CCBL2 | 1,951 | 5,0054295638863E-32 |
| RP1-8B22.1 | 10,891 | 1,24983470680231E-24 | C1orf50 | 0,687 | 0,00151825698617437 | PTGES3P1 | 0,919 | 7,42163512461959E-23 |
| RP13-279N23.2 | 2,970 | 0,000281158171258873 | CCDC23 | 0,887 | 1,36823562492196E-09 | RP4-620F22.3 | 11,387 | 4,90177607229541E-27 |
| ALDH4A1 | 2,708 | 0,000373315156142096 | ERMAP | -0,977 | 6,2862355488706E-07 | GEMIN8P4 | 1,281 | 0,000255512948011381 |
| RP5-1126H10.2 | -3,178 | 4,98796183588303E-08 | ATP6V1E1P1 | 7,122 | 1,08320082920021E-40 | ZNF326 | 1,790 | 2,15026746755947E-13 |
| RP1-43E13.2 | -1,223 | 0,00663985796314327 | EBNA1BP2 | 1,761 | 7,38771313361438E-41 | RPL5P6 | 6,568 | 1,00389123176133E-160 |
| CAPZB | 2,271 | 9,71423783931502E-85 | RP1-92O14.6 | -4,404 | 3,41162744451514E-39 | CDC7 | 0,918 | 1,42421633723715E-06 |
| MINOS1-NBL1 | -1,257 | 4,30866041152169E-20 | PTPRF | -1,964 | 5,48689463435607E-13 | HSP90B3P | 2,421 | 3,57715813365871E-05 |
| MINOS1 | -1,263 | 2,53910633501865E-20 | ST3GAL3 | 1,385 | 0,00289315083958795 | EPHX4 | 2,288 | 7,59292814041606E-07 |
| CAMK2N1 | -2,412 | 0,0096435330409362 | SHMT1P1 | 6,523 | 8,44693349778617E-09 | PRKAR1AP | 8,894 | 8,03592622571893E-16 |
| PINK1 | -2,948 | 1,49090705750552E-11 | IPO13 | -1,329 | 0,000464234241193166 | GLMN | 2,446 | 5,39356263108907E-50 |
| PINK1-AS | -1,241 | 4,61304712607242E-23 | DPH2 | -1,066 | 0,000174947571410807 | RPAP2 | 1,154 | 1,63614252195842E-08 |
| AL391357.1 | -3,586 | 1,12439128770141E-06 | B4GALT2 | -1,305 | 2,30828745216518E-07 | GFI1 | -3,384 | 0,00988344796850662 |
| DDOST | 0,646 | 7,03027996658148E-08 | RP11-570P14.1 | 1,165 | 2,5621135161219E-10 | EVI5 | -2,883 | 6,97913764631533E-05 |
| RP5-930J4.4 | -2,100 | 1,39558381711789E-08 | DMAP1 | 1,513 | 7,58095639464497E-11 | RP11-330C7.3 | 4,574 | 0,000842850735431535 |
| ECE1 | 1,591 | 0,000261954823524422 | ERI3 | 1,756 | 9,87656783001626E-37 | RPL5 | -5,988 | 6,95314204797785E-279 |
| PPP1R11P1 | 5,399 | 0,000699274635165124 | TMEM53 | 1,073 | 4,81244514898271E-06 | FAM69A | 1,322 | 7,37878050660911E-05 |
| RAP1GAP | 2,212 | 0,00980665165316651 | KIF2C | 1,757 | 3,50501349189403E-21 | RP11-386I23.1 | 7,431 | 2,66939395441855E-16 |
| USP48 | 1,692 | 8,35863091001225E-05 | RP11-269F19.2 | -8,746 | 0 | MTF2 | 0,976 | 0,00633266709323089 |
| CDC42 | -2,253 | 1,72246482259323E-85 | RPS8 | -5,891 | 0 | TMED5 | 1,078 | 1,47500521607628E-17 |
| RP1-224A6.8 | 7,387 | 3,15196792298176E-10 | SNORD38A | -1,943 | 0,00133443385258669 | DR1 | -0,716 | 0,00436462221044719 |
| KDM1A | -1,048 | 1,94453991715143E-06 | RPS15AP11 | -1,974 | 1,11129791064127E-44 | DNTTIP2 | 1,194 | 1,83733495817888E-14 |
| TCEA3 | 2,860 | 0,000156858579836695 | RP5-882O7.1 | -1,942 | 5,95283593739482E-14 | GCLM | -1,847 | 5,51954277302952E-06 |
| RP4-654C18.1 | 11,410 | 4,54108830776627E-27 | EIF2B3 | 2,748 | 4,51563081135253E-46 | ABCD3 | 1,292 | 2,68066657290548E-21 |
| ID3 | 0,874 | 2,50706818245733E-06 | CCNB1IP1P1 | 10,278 | 1,82813744424086E-21 | SLC44A3 | 3,068 | 0,0015354300929012 |
| LYPLA2 | 1,850 | 8,59217163131992E-18 | MRPS17P1 | 6,176 | 1,54186096155339E-50 | CNN3 | 0,795 | 7,26977334933568E-08 |

| | | | | | | | | |
|---|---|---|---|---|---|---|---|---|
| HMGCL | 2,022 | 7,44053483267472E-19 | ZSWIM5 | -2,355 | 0,002335515113634 | ALG14 | 1,055 | 4,40410595781346E-15 |
| FUCA1 | 2,031 | 2,30205131023778E-11 | OSTCP5 | 7,823 | 3,48065402615338E-13 | TMEM56-RWDD3 | -0,783 | 0,000119162798252621 |
| PNRC2 | -1,254 | 6,60999207194997E-11 | TESK2 | -0,813 | 0,000160406773411041 | RP4-736I12.1 | 7,812 | 2,41985146959689E-10 |
| RP11-4M23.7 | 4,906 | 8,26073513951221E-113 | MMACHC | -1,029 | 1,08386681830762E-10 | UBE2WP1 | 6,796 | 1,05267184503061E-07 |
| STPG1 | 1,095 | 3,95160242804717E-06 | PRDX1 | -0,982 | 1,02855059842032E-09 | EEF1A1P11 | -2,180 | 1,86088223279477E-50 |
| NIPAL3 | 3,504 | 9,10572830554631E-41 | AKR1A1 | 0,856 | 5,14641959080086E-06 | DPYD | 1,569 | 9,67082500158421E-05 |
| SRRM1 | -2,569 | 8,23745881383437E-16 | NASP | 0,716 | 0,000604888642364079 | SNX7 | 0,902 | 0,000298257879295761 |
| CLIC4 | -1,474 | 2,07506255082484E-05 | GPBP1L1 | 0,762 | 0,00641778923054913 | RP11-234N17.1 | -0,961 | 1,96963116146486E-05 |
| SYF2 | 2,686 | 6,73826236711061E-82 | RP11-767N6.7 | -1,811 | 0,00615906958123846 | RP4-735N21.1 | 8,576 | 2,07146250098873E-14 |
| TMEM50A | 2,196 | 3,35288948328596E-109 | MAST2 | -2,417 | 6,5193989173466E-05 | HIAT1 | 1,858 | 2,21976146015437E-08 |
| SEPN1 | -3,558 | 9,82753938745095E-14 | TMA16P2 | 7,365 | 1,81489596580484E-16 | RP4-714D9.2 | -2,234 | 0,00217944007451532 |
| RP1-317E23.3 | -3,467 | 8,16363487354865E-08 | RP4-533D7.3 | 5,981 | 3,22031009848375E-10 | TRMT13 | 1,269 | 2,70536121386799E-05 |
| MTFR1L | 1,259 | 2,09325507364713E-11 | UQCRH | -4,518 | 1,52609469817803E-303 | RP11-305E17.7 | -4,512 | 0,0020900207275366 |
| AL020996.1 | -1,770 | 6,47163330770738E-06 | NSUN4 | 1,213 | 0,000756825240895426 | BRI3P1 | -1,562 | 0,00776566071339968 |
| STMN1 | 0,660 | 5,83971527255857E-05 | MKNK1 | 1,345 | 4,01747145000569E-14 | RTCA | 2,400 | 2,39044747403544E-71 |
| SNRPFP2 | 9,381 | 2,41733350909143E-18 | RP11-184J23.2 | 4,587 | 0,0032321161331459 | RP5-837M10.1 | 7,702 | 6,66215338665728E-12 |
| MIR3917 | -5,792 | 2,77399936389499E-124 | CMPK1 | 0,885 | 1,02230230629512E-08 | BCAS2P2 | 7,814 | 1,2756369363013E-33 |
| PAFAH2 | 1,609 | 0,00326206375849546 | ATP6V0E1P4 | 5,330 | 0,000886503028266585 | SLC30A7 | 1,129 | 2,53171866444803E-06 |
| SH3BGRL3 | 0,695 | 2,31234065742044E-05 | SPATA6 | 1,979 | 7,46861431493859E-05 | DPH5 | 1,579 | 8,91750494899465E-27 |
| AIM1L | 3,562 | 0,000706535062681768 | PPP1R8P1 | 8,292 | 1,32388560202998E-13 | AC093157.1 | -0,664 | 0,000211899924544176 |
| DHDDS | 2,369 | 1,57100780659892E-17 | FCF1P6 | 8,115 | 5,29656607970535E-13 | RP4-575N6.2 | 5,794 | 5,34452913586887E-05 |
| HMGN2 | -1,385 | 1,1119322964001E-31 | PHBP12 | 5,263 | 5,05914246721678E-05 | RP11-556K13.1 | -0,711 | 1,40478050750228E-05 |
| ARID1A | -2,269 | 2,27797616154994E-08 | RP11-296A18.3 | 4,099 | 2,87891289669183E-09 | RNPC3 | -1,724 | 0,000197144956343178 |
| ZDHHC18 | -2,406 | 7,57594816575853E-05 | RNF11 | -1,058 | 0,00149011372550514 | AMY2B | -2,012 | 0,000435625294687376 |
| NUDC | -1,456 | 3,75419369752517E-12 | EPS15 | 0,852 | 0,0038262158121156 | SEPT2P1 | 9,741 | 9,78922967563965E-20 |
| OSTCP2 | 11,474 | 1,81780875848327E-27 | NRD1 | 2,044 | 1,43379446446874E-09 | RP11-90H3.2 | 6,400 | 3,48247846390833E-16 |
| CHCHD3P3 | 0,916 | 2,10692960891768E-05 | RP4-657D16.3 | -1,767 | 1,12562050981731E-07 | RP11-90H3.1 | 6,557 | 1,1177416507097E-17 |
| NPM1P39 | -2,608 | 6,95398087880292E-76 | TXNDC12 | 1,646 | 6,35007984729081E-33 | PRMT6 | -1,632 | 1,44701927662103E-05 |
| TMEM222 | 0,715 | 5,12416275321084E-06 | KTI12 | -1,674 | 0,000275377469910943 | FAM102B | -2,827 | 0,000214218524500727 |
| WASF2 | -1,782 | 1,93474502857908E-06 | BTF3L4 | -2,955 | 1,46204676188976E-100 | STXBP3 | 2,038 | 1,50996527423109E-13 |
| RP1-159A19.3 | -1,028 | 7,92292500771062E-17 | RP4-800M22.2 | 9,792 | 6,56304325579787E-20 | CLCC1 | 1,781 | 2,69961233684189E-21 |
| IFI6 | 1,203 | 0,00927472794858125 | | | | | | |

### 48 hours stimulation

| Gene | log2FoldChange | padj | Gene | log2FoldChange | padj | Gene | log2FoldChange | padj |
|---|---|---|---|---|---|---|---|---|
| CNTN1 | 2,569 | 9,99104999366897E-11 | FOCAD | 1,594 | 9,49123503403451E-17 | EIF1AX | -2,449 | 8,99395721305158E-21 |
| GPN1 | 2,403 | 9,99053300843396E-22 | ARNTL | 1,738 | 9,49022882675831E-06 | ATF1 | 1,465 | 8,99292642909882E-09 |
| TMEM185AP1 | 2,159 | 9,98111120585312E-05 | BTF3L4 | -2,542 | 9,48485265378265E-56 | TCAIM | 1,962 | 8,99207599281752E-14 |
| SUPV3L1 | 2,457 | 9,978555673598E-32 | AP1M2 | 1,372 | 9,484653041055E-07 | NDUFA5 | -0,795 | 8,9899176126702E-05 |
| RP4-814D15.2 | -2,702 | 9,97809419217563E-52 | SOAT1 | 1,949 | 9,47720349378363E-13 | ZBTB25 | -2,691 | 8,98649150060667E-34 |
| SFPQP1 | 7,745 | 9,97490296216159E-12 | CTD-2616J11.14 | 3,919 | 9,47538418761008E-10 | APOPT1 | 1,039 | 8,98438838205269E-07 |
| SPINT2 | 3,009 | 9,96714284031678E-25 | PCNPP1 | 7,108 | 9,46626194936375E-11 | SLC12A8 | 1,324 | 8,98025275182784E-06 |
| RFC5 | 2,167 | 9,96428844778841E-18 | SUGT1P2 | 3,043 | 9,45834952828872E-27 | NIP7 | -1,359 | 8,97471990267071E-14 |
| VAMP3 | 0,907 | 9,96169701629963E-06 | RP11-222O23.1 | 7,180 | 9,45222789468369E-10 | NIFKP8 | 7,392 | 8,97411601858324E-11 |
| C4orf33 | 1,009 | 9,95774994318844E-05 | TRIM38 | 1,450 | 9,45044773578504E-05 | RASGEF1B | 2,013 | 8,97398476889227E-06 |
| MAGEA3 | -4,593 | 9,95676748301443E-22 | MIR4700 | -1,827 | 9,44919062996712E-05 | AC079250.1 | -0,812 | 8,97280995239448E-06 |
| COX6CP4 | 9,004 | 9,94450059765681E-17 | AC092106.1 | 10,761 | 9,44881384335533E-24 | BCRP6 | 2,572 | 8,9672993845956E-18 |
| EXT2 | 2,150 | 9,93836254840388E-20 | RP11-365O16.6 | -1,092 | 9,44866694303467E-07 | MEIS1 | 1,729 | 8,96527621459173E-08 |
| ACTR10 | 2,367 | 9,93770468666504E-30 | ARPC4-TTLL3 | 1,656 | 9,44586666126476E-10 | PRPF40A | 1,666 | 8,96415531341183E-18 |
| EEF1B2P1 | -3,768 | 9,93770468666504E-30 | PVT1 | 1,578 | 9,44060854787745E-11 | SPG20 | 1,675 | 8,96359063311635E-13 |
| RP11-475I24.3 | -1,912 | 9,93655176183688E-19 | CTA-276O3.4 | 2,621 | 9,4385321881518E-51 | SMNDC1 | 1,285 | 8,96126016710915E-06 |
| SSB | -1,242 | 9,93529090063621E-20 | RAD1 | -2,216 | 9,43558809903619E-38 | PSMD12P | 6,042 | 8,95773900141324E-34 |
| NECAP1P1 | 5,258 | 9,9349779005146E-05 | PIK3R4 | 1,051 | 9,43476387572631E-05 | TRMT112 | -0,946 | 8,95413572252286E-11 |
| DYNLT3P1 | 2,034 | 9,93448556394341E-10 | POLRMT | -1,711 | 9,43476387572631E-05 | AC125238.2 | 7,454 | 8,95250539613693E-41 |
| SEPT7P6 | 5,279 | 9,93092070014331E-28 | RP11-382A20.1 | -4,789 | 9,42864070362974E-19 | TOP1 | 1,325 | 8,95169422921065E-13 |
| HS1BP3 | 1,178 | 9,92600363556472E-05 | PHBP10 | 5,520 | 9,42677816092061E-49 | YWHAEP1 | 7,587 | 8,95168480736109E-54 |

| | | | | | | | | |
|---|---|---|---|---|---|---|---|---|
| LAMTOR3P1 | 5,698 | 9,92587260621431E-06 | RP11-345F18.1 | 1,301 | 9,42072664958198E-05 | RP13-820C6.2 | -1,744 | 8,94874739712929E-05 |
| PDHA2 | 8,804 | 9,92441922868433E-16 | RPL35P2 | 0,744 | 9,4203087787573E-07 | ZNF638 | 1,121 | 8,94738442810468E-10 |
| PRPS2 | 1,747 | 9,90995546987684E-09 | MAGOH2 | 10,786 | 9,41174032244569E-25 | RP11-140K17.3 | -1,128 | 8,94709025160793E-05 |
| PUS7 | 1,953 | 9,90952251495217E-16 | RP11-195F19.9 | -1,698 | 9,40385563083855E-05 | HYPK | -1,315 | 8,9450115443392E-19 |
| CASP3P1 | 6,104 | 9,90926206822209E-07 | TBC1D7 | 2,104 | 9,39886711045411E-22 | STAU1 | 1,379 | 8,93687748093368E-10 |
| FAM86C2P | 1,104 | 9,90926206822209E-07 | AC002310.12 | -1,014 | 9,39730517094363E-05 | RP11-397K18.1 | 7,046 | 8,92353594353357E-10 |
| LL22NC03-80A10.6 | 0,752 | 9,90748597601953E-05 | RP11-460N20.4 | 5,375 | 9,39319771406697E-06 | TCTN2 | 1,880 | 8,92193215717372E-08 |
| RAD18 | 2,140 | 9,90575590769501E-17 | AC005534.6 | 2,441 | 9,39276453468121E-08 | WI2-89927D4.1 | 5,209 | 8,92140630616345E-05 |
| RER1 | 1,299 | 9,90487413291955E-16 | RINT1 | 1,244 | 9,39151244788815E-07 | SUCLA2 | -1,202 | 8,91023585039722E-10 |
| NACAP2 | 5,507 | 9,90487115485203E-06 | CALM1P1 | 8,631 | 9,38547700389382E-16 | UBL5P2 | 2,541 | 8,90997811437776E-64 |
| AC009963.6 | 9,555 | 9,90423191384172E-19 | CARS | 3,448 | 9,37693435401502E-41 | RP11-129B9.2 | 9,136 | 8,90671955202481E-17 |
| RP11-601I15.1 | 9,485 | 9,90402757222612E-111 | SPPL2A | -2,325 | 9,37326778375564E-06 | ZNF771 | -1,411 | 8,90247834460256E-18 |
| RP11-151F5.2 | 1,887 | 9,90311691435436E-11 | EMC2 | 1,896 | 9,36661589374129E-20 | CTD-2230M5.1 | 5,619 | 8,89945668423762E-06 |
| RPL34P33 | 5,478 | 9,90118814699539E-135 | RNA5S2 | -2,023 | 9,36618816689945E-06 | DDX28 | -1,267 | 8,89844133860288E-11 |
| RPS15 | -1,645 | 9,90084151892275E-40 | MOXD1 | 1,079 | 9,36618816689945E-06 | RP11-241I20.5 | 7,451 | 8,89472717601003E-11 |
| PIK3R2 | 0,676 | 9,90011052541634E-05 | SEC23IP | 1,748 | 9,36600861461708E-13 | RP11-764K9.1 | -5,062 | 8,88843655497146E-28 |
| HEXA | 2,063 | 9,89442956316107E-18 | RP11-963H1.1 | 3,213 | 9,36277028441986E-07 | UBE2CP4 | 8,465 | 8,88783168977946E-15 |
| TSC1 | 1,419 | 9,8941530951906E-10 | RP11-311H10.7 | 8,076 | 9,3616255628735E-13 | KLHL41 | 2,733 | 8,88030170736569E-09 |
| RP11-467H10.2 | 2,444 | 9,89342378869059E-09 | MKLN1 | 2,970 | 9,35626557511291E-24 | FAM126A | 1,108 | 8,87894943442602E-05 |
| ZC3HC1 | 2,104 | 9,89332871063647E-20 | EEF1GP5 | -2,862 | 9,35503462780423E-50 | RBM5 | 2,679 | 8,87157935782942E-26 |
| USP44 | 1,688 | 9,88800415243541E-05 | TEKT2 | -1,689 | 9,35356302873424E-05 | AP2M1 | 0,666 | 8,87042925310167E-05 |
| CDC45 | 2,847 | 9,88574601991247E-38 | ERMARD | 2,593 | 9,34973031942582E-17 | DTYMK | 1,213 | 8,86846381365421E-09 |
| AC008753.3 | 2,031 | 9,88341958214837E-05 | FAM86FP | 2,267 | 9,34617326443233E-16 | LRRC37A7P | 8,932 | 8,86698128913606E-16 |
| HOXB6 | -1,251 | 9,88088745500902E-08 | ARL17B | 1,126 | 9,3441979745165E-08 | RP11-549B18.3 | 8,932 | 8,86698128913606E-16 |
| MTND1P23 | -1,622 | 9,88024801567521E-08 | CCDC82 | 1,501 | 9,34193227538088E-05 | NPM1P18 | 7,443 | 8,86641101669922E-17 |
| SPATS2L | 1,434 | 9,87858830261559E-06 | CTD-2576F9.2 | 2,677 | 9,33961483777211E-07 | TSG101 | 2,294 | 8,86597473614292E-21 |
| VAMP4 | 1,453 | 9,8775531306768E-09 | GAPDHP40 | 2,352 | 9,33926757503834E-05 | RP11-382B18.3 | 3,409 | 8,8654575385228E-08 |
| PRPS1P2 | 7,789 | 9,86651911295209E-18 | RP11-378J18.6 | 11,441 | 9,33862367502995E-27 | RP11-478B9.3 | 8,424 | 8,8621711720362E-14 |
| DMD | 2,192 | 9,86039630394431E-14 | AC011737.2 | -0,773 | 9,33223728170825E-06 | RP11-113I24.1 | 6,355 | 8,85530005688892E-08 |
| CHRFAM7A | 1,404 | 9,86012958113128E-05 | RP11-93O17.1 | 7,819 | 9,33202064607782E-12 | AP3B1 | 2,685 | 8,85529451243548E-20 |
| EEF1GP1 | 2,692 | 9,85386346592221E-06 | SOX6 | -3,255 | 9,33163399910849E-26 | PTPRK | 2,060 | 8,85529451243548E-20 |
| CTB-55O6.10 | -0,836 | 9,85386346592221E-06 | DYX1C1-CCPG1 | 1,266 | 9,33126519908869E-05 | CTD-2314B22.2 | -1,416 | 8,85494564593847E-13 |
| INTS4 | 2,415 | 9,84903121268051E-16 | TTC12 | 3,226 | 9,32363793162771E-25 | RTCA | 2,193 | 8,8526617909503E-27 |
| CTD-2336O2.2 | 5,236 | 9,84675318995704E-05 | ABCF2 | -1,371 | 9,32330628792922E-09 | CCNDBP1 | 2,085 | 8,85166541755599E-17 |
| TRAPPC5 | -1,800 | 9,8465768987626E-07 | SYAP1 | 1,307 | 9,32319706841473E-14 | HNRNPA1P6 | 5,767 | 8,84838208279668E-06 |
| RP11-274E7.2 | -0,769 | 9,84182093456644E-06 | TBC1D19 | 3,186 | 9,32008970613892E-26 | COX17P1 | -1,108 | 8,84567545676854E-15 |
| COQ10B | 1,251 | 9,83634576635552E-07 | TUBG1P | 6,936 | 9,31925317581869E-30 | STARD5 | 2,019 | 8,84346486807408E-08 |
| ORC1 | 2,549 | 9,83601181873374E-27 | AMPH | 2,907 | 9,31644583467492E-12 | SPDYE9P | -1,996 | 8,84227160180165E-07 |
| FAM134A | -1,496 | 9,83426726786644E-05 | FASTKD2 | 1,775 | 9,31509223124271E-10 | GPR89B | -1,080 | 8,84061259427472E-10 |
| RPL34P34 | 5,102 | 9,816303522032E-180 | SEC61A2 | -0,727 | 9,31496081695843E-05 | RP11-587D21.1 | -1,206 | 8,83115948158616E-14 |
| UBA52P3 | 8,254 | 9,81428572415023E-50 | PSMD10 | -0,673 | 9,31452062206106E-05 | CETN2 | -1,039 | 8,82829467059521E-10 |
| RP11-34P1.2 | 5,170 | 9,80888371711373E-117 | HDAC8 | 1,717 | 9,31224453285355E-14 | RP11-573M3.4 | 6,915 | 8,82815797199073E-10 |
| RP5-1165K10.2 | 1,344 | 9,80151447083456E-10 | RP11-369K17.2 | 5,784 | 9,31197058658014E-06 | COG8 | -1,141 | 8,82637844925779E-05 |
| PTMAP2 | -0,994 | 9,79697471811215E-07 | EEF1B2 | -3,931 | 9,29819250726121E-108 | DSTNP4 | 5,655 | 8,82514432863456E-06 |
| RP11-430L17.1 | 4,161 | 9,79355444788456E-20 | PLRG1 | 1,834 | 9,29282787059052E-15 | CTD-2194M22.1 | 5,265 | 8,81744954883557E-05 |
| RPL5P1 | 1,960 | 9,79355444788456E-20 | CANX | 1,550 | 9,29282787059052E-15 | REEP5 | 1,161 | 8,81117029484242E-12 |
| BSCL2 | 1,564 | 9,79250506848749E-11 | RPL30P13 | 5,582 | 9,29237952484496E-06 | AC097711.1 | 11,056 | 8,80452170407513E-25 |
| TMEM14B | -1,038 | 9,79182144355883E-10 | F8 | -5,060 | 9,28911579719677E-51 | RNFT1 | 2,024 | 8,80029819724411E-16 |
| CTC-428G20.6 | -1,894 | 9,79114897799633E-07 | PSMD12 | -1,459 | 9,28707831238385E-16 | CEP72 | 1,732 | 8,79983194607218E-05 |
| NAA11 | 11,313 | 9,79114219348328E-26 | MRPL24 | -0,798 | 9,28588308598509E-07 | CCS | 0,941 | 8,79749866314626E-05 |
| THOC3 | 0,993 | 9,7879467908992E-08 | RPL37AP1 | 2,208 | 9,28327301849697E-54 | CCT3 | 1,863 | 8,79470247460705E-17 |
| ZNF215 | 1,405 | 9,78439949632344E-06 | RP11-430C7.2 | 7,437 | 9,28257822110036E-11 | NFX1 | 1,925 | 8,7940271871403E-12 |
| OSBPL9P3 | 7,497 | 9,77686380605483E-12 | TEAD2 | 1,473 | 9,28036114253431E-07 | TIMM13 | -0,751 | 8,78212579723056E-07 |
| PTDSS1 | 2,213 | 9,77629975138E-18 | CTD-2287O16.5 | -1,556 | 9,27559187612699E-08 | AC007899.3 | 1,774 | 8,78041298583127E-07 |
| TBC1D3 | 2,023 | 9,7756575517457E-14 | GBE1 | 2,254 | 9,27191974836012E-11 | RP11-192M23.1 | 5,638 | 8,77978304252558E-06 |
| PAFAH1B1 | -1,144 | 9,77513194032924E-05 | RMDN1 | 1,806 | 9,27130126900154E-18 | R3HCC1 | 1,179 | 8,76890121883768E-06 |

| | | | | | | | | |
|---|---|---|---|---|---|---|---|---|
| SLC9B1 | -0,780 | 9,77458342165795E-05 | MRPL42P6 | 8,814 | 9,26636931679682E-16 | TCEB1 | -1,190 | 8,76725938815342E-14 |
| ATP5A1P1 | 8,796 | 9,77185082585103E-16 | AKR1A1 | 0,942 | 9,26341308286879E-06 | TAF1 | -2,911 | 8,76288996726473E-12 |
| EIF3F | -3,414 | 9,76856045684495E-07 | SLC18B1 | 1,753 | 9,25960976771109E-17 | RP11-575L7.8 | -7,570 | 8,76161714124031E-211 |
| TMEM138 | 0,890 | 9,76165397766377E-05 | NEK11 | 3,113 | 9,25879676855459E-10 | MYL5 | -1,655 | 8,76050754266984E-37 |
| GFPT1 | 1,740 | 9,75992229753287E-13 | THRAP3P1 | 4,826 | 9,25822767366946E-18 | TMTC4 | 1,447 | 8,7568478867084E-10 |
| C1orf198 | -1,680 | 9,75834402086167E-08 | RAD17P1 | 7,798 | 9,2560677673617E-12 | TMCO6 | -1,050 | 8,75452782483632E-09 |
| GAD1 | 1,862 | 9,75602850100077E-07 | RP11-817O13.1 | 5,378 | 9,25590725145735E-05 | RP11-363E7.4 | -4,259 | 8,75413062159589E-05 |
| RP11-106A1.2 | -3,399 | 9,7535397628231E-20 | MRS2P2 | 9,035 | 9,25322247907145E-16 | CDC42SE2 | 1,204 | 8,7539773526189E-05 |
| ATF7IP | 1,757 | 9,75092101575312E-08 | BCAS2 | -0,733 | 9,2471036612232E-05 | MRPS18C | -3,672 | 8,74984906228451E-157 |
| PPIH | -3,553 | 9,7496680129987E-71 | GUSB | 1,912 | 9,24529073240973E-17 | PLSCR1 | 1,275 | 8,74975996654484E-09 |
| RP11-430A19.2 | 5,799 | 9,74822808823765E-07 | TXNP5 | 9,360 | 9,24387834457147E-51 | MT-TE | 3,931 | 8,74147061485545E-10 |
| METTL2B | 2,101 | 9,74246251452129E-28 | SLC39A9 | -2,544 | 9,24280743921908E-53 | AL132780.1 | -1,019 | 8,73795352295248E-08 |
| COX4I1P1 | 7,415 | 9,73930706940271E-125 | AP3S2 | 0,985 | 9,24056177888007E-06 | RP11-109G23.1 | 5,296 | 8,73735299972573E-05 |
| RTTN | 1,303 | 9,73303205838395E-05 | RPL7P46 | 3,055 | 9,24042397200007E-14 | ZNF473 | 1,358 | 8,73735299972573E-05 |
| MT-CO1 | -1,981 | 9,73072638170269E-11 | MAPK6PS3 | 5,159 | 9,24007895269255E-05 | CCDC132 | 2,981 | 8,73484673392342E-18 |
| NDUFB4 | -3,017 | 9,71467723593944E-100 | HSPB11 | 0,887 | 9,23941984479778E-09 | PHF19 | 0,985 | 8,72887546957398E-06 |
| PIGF | -1,690 | 9,71417464648565E-22 | SNORD42A | 2,682 | 9,23868509742797E-05 | GPR89A | -1,117 | 8,7232296167508E-11 |
| DYNC2LI1 | 2,072 | 9,7063705931161E-16 | DHRS7B | 1,945 | 9,22542262829152E-16 | PMS1 | 2,264 | 8,71959814043789E-16 |
| RPS27P27 | 12,084 | 9,70575846008482E-31 | GGNBP2 | 1,374 | 9,22490209325067E-09 | CTD-2340F8.1 | 4,136 | 8,71467021525746E-09 |
| SHQ1 | 1,558 | 9,70445231489489E-09 | OSTCP1 | 4,010 | 9,22277922937072E-52 | MTND2P28 | -2,134 | 8,71355589565888E-14 |
| NEK3 | 2,108 | 9,70202173582023E-11 | RPL5P18 | 3,839 | 9,2157028783609E-06 | VSNL1 | 1,984 | 8,71069617343442E-10 |
| PNISR | 1,241 | 9,70117352507219E-08 | THOC2 | 1,649 | 9,21361850738961E-14 | RP11-16F15.1 | -2,501 | 8,70856877936592E-84 |
| ANKRD20A12P | -2,458 | 9,69823200613351E-13 | HSPA9 | -2,178 | 9,21271771320248E-12 | MTMR6 | 1,630 | 8,70349173079226E-05 |
| HIST1H4J | -2,213 | 9,69823200613351E-13 | RP11-106M3.2 | 1,145 | 9,20607494157767E-06 | SAGE1 | 1,591 | 8,69825196786697E-08 |
| RPL10AP6 | -0,785 | 9,69613375355557E-06 | PHBP8 | 10,482 | 9,20086892574591E-23 | RP11-150O12.6 | -3,007 | 8,69485778081901E-16 |
| RP11-231C14.3 | -1,382 | 9,69500361732484E-22 | PSPH | 2,316 | 9,19818249152243E-18 | SKA1 | 1,783 | 8,69368452348885E-12 |
| RP11-442P12.1 | 6,028 | 9,69368416080577E-07 | FCHO2 | 1,944 | 9,19278208982521E-08 | ACAT1 | 2,313 | 8,69172135352727E-25 |
| PLAT | 1,779 | 9,6922270471871E-06 | RP5-961K14.1 | 2,115 | 9,18968152610147E-05 | TANGO6 | 2,483 | 8,68969555691208E-13 |
| RP11-468E2.5 | -2,215 | 9,69148015232185E-20 | FAM35DP | -0,797 | 9,18685178094869E-07 | NDUFB1P1 | 1,992 | 8,68871083397647E-44 |
| XPO7 | 2,389 | 9,69050482706298E-19 | GSTM2P1 | 8,542 | 9,18644791500498E-15 | CTD-2186M15.1 | 5,509 | 8,68585971006725E-142 |
| TUBE1 | 1,179 | 9,68972606594945E-06 | DCAF12 | 1,103 | 9,18348972543055E-06 | ABHD10 | 1,051 | 8,68585354143023E-05 |
| FDPSP6 | 5,301 | 9,68701992920372E-05 | RPL13AP20 | -3,091 | 9,18306793008965E-118 | AP1S1 | 1,654 | 8,68501073398524E-11 |
| MAN1B1 | 1,771 | 9,68580914473062E-09 | BBS4 | 2,129 | 9,17684715786953E-15 | DDB2 | 1,114 | 8,6840538940329E-05 |
| RP11-159H3.1 | -2,026 | 9,68122521112924E-07 | RPTOR | 1,560 | 9,16842618614726E-05 | MCCC1 | 3,074 | 8,67846133836259E-30 |
| ADK | 2,243 | 9,68111307432914E-24 | RP11-457D20.1 | 5,166 | 9,16449112341315E-13 | VDAC1P6 | 5,046 | 8,67846133836259E-30 |
| RP11-288E14.2 | 0,749 | 9,67233388278599E-05 | OMA1 | 2,119 | 9,16373674685641E-18 | RNVU1-4 | -1,936 | 8,67457812414276E-06 |
| CEP41 | 1,899 | 9,67172028296706E-14 | SNAP23P | 8,076 | 9,15202433144362E-13 | RNVU1-20 | -1,936 | 8,67457812414276E-06 |
| NDRG3 | 2,912 | 9,66434707397925E-33 | RP11-111F16.2 | 1,241 | 9,14589586286606E-05 | RP11-229M1.2 | 2,672 | 8,67406901534895E-10 |
| TGFB3 | 2,881 | 9,65894016722533E-11 | PRPF39 | -0,982 | 9,14328027433155E-09 | LRRCC1 | 1,982 | 8,67388877561265E-07 |
| SMARCAL1 | 2,465 | 9,65839547104695E-13 | AC024560.3 | 1,243 | 9,13912508303723E-07 | TRAPPC11 | 1,799 | 8,67373443266444E-11 |
| SP110 | 2,054 | 9,65736652361287E-11 | MELK | 2,892 | 9,13569499706554E-30 | RP11-75A9.3 | 6,225 | 8,66486289879441E-08 |
| SLC12A3 | 4,007 | 9,65672219947875E-18 | CBX5P1 | 9,627 | 9,13121401784363E-19 | RFC3 | 1,365 | 8,66486289879441E-08 |
| MTRNR2L2 | -1,617 | 9,65613841703128E-10 | RARS2 | 2,999 | 9,12308336625956E-48 | RP11-416K24.2 | 7,962 | 8,66319872546272E-115 |
| RP11-72I8.1 | -2,544 | 9,65513661925749E-09 | RP11-573D15.9 | -3,435 | 9,11983662574099E-36 | NRG4 | 2,363 | 8,66246540652915E-20 |
| FAM96B | -0,865 | 9,65513661925749E-09 | EIF4EBP1P2 | 6,842 | 9,11593071936616E-09 | MMADHC | -1,386 | 8,66059304073346E-18 |
| AARS | 2,310 | 9,65414519295458E-26 | ZNF747 | -1,829 | 9,11083521911218E-14 | CASP7 | 1,159 | 8,65821182492887E-06 |
| MBTPS2 | 0,941 | 9,65024304731121E-05 | PRMT2 | 2,151 | 9,10977436360038E-23 | RP11-1079K10.4 | -2,558 | 8,65678516629227E-59 |
| UQCRC1 | 1,373 | 9,65008603484275E-11 | TCF25 | 1,300 | 9,10231603183321E-07 | HMGN2P7 | -2,499 | 8,6505840213597E-37 |
| RP11-43D4.2 | 7,650 | 9,64436747869951E-12 | ADAT2 | 1,196 | 9,10147039878629E-09 | RPA3 | -1,340 | 8,64766042234062E-19 |
| RP11-75L1.2 | -1,300 | 9,64388899435049E-12 | RPS8 | -5,118 | 9,10128948059676E-74 | RP11-562A8.4 | -2,721 | 8,64577714741528E-07 |
| TMA16P1 | 7,660 | 9,64032410757028E-12 | RP1-199J3.5 | 4,196 | 9,10090995044606E-12 | SRD5A3P1 | 8,924 | 8,64457328722844E-16 |
| RPL3 | -4,637 | 9,63373310385877E-68 | TPD52L2 | 2,045 | 9,09884226800737E-16 | AC107081.5 | -0,902 | 8,64235207005635E-07 |
| RP11-69L16.3 | 8,770 | 9,63150780504113E-16 | COX5BP6 | 7,434 | 9,09500127807188E-12 | SLC25A14P1 | 7,131 | 8,63205195417821E-11 |
| ANAPC10P1 | 7,764 | 9,62822128144021E-13 | RP11-804F13.2 | 6,473 | 9,09454362008114E-08 | AP000343.1 | -4,023 | 8,62779828687536E-05 |
| SUMF1 | 1,849 | 9,62822128144021E-13 | GS1-115G20.2 | 6,155 | 9,09425984316197E-07 | RP11-43F13.1 | 2,389 | 8,6276733727614E-25 |
| CTD-2008A1.2 | 1,856 | 9,62822128144021E-13 | PPP1R14BP2 | -3,232 | 9,09261973102676E-07 | DHDDS | 2,148 | 8,6270107297323E-14 |

| | | | | | | | | |
|---|---|---|---|---|---|---|---|---|
| DHX57 | 1,700 | 9,62623724146729E-09 | PHTF2 | 1,280 | 9,08731058522397E-06 | ISCA2 | -0,772 | 8,61286136146567E-06 |
| XPNPEP3 | -0,658 | 9,62358084816218E-05 | DNALI1 | -3,054 | 9,08658559403377E-15 | H3F3B | -0,892 | 8,61030389647404E-09 |
| RP11-452J6.2 | 4,489 | 9,62303170830216E-05 | HNRNPUL2-BSCL2 | 1,435 | 9,0809455159568E-10 | KNTC1 | 1,923 | 8,61016313852407E-11 |
| RP11-564C24.1 | 7,632 | 9,62031363584391E-12 | SIL1 | 1,426 | 9,08017296621078E-07 | COMMD6 | 0,922 | 8,60962375880692E-09 |
| TXNDC12 | 1,620 | 9,61685970196953E-13 | ZNF451 | 1,783 | 9,07523044947585E-10 | RPL36A-HNRNPH2 | -6,460 | 8,60916832661038E-275 |
| CRBN | 1,706 | 9,61685970196953E-13 | PRELID1P1 | -1,206 | 9,07327997020729E-05 | RP11-178H8.3 | 6,086 | 8,60176417626137E-08 |
| RP11-726G1.1 | 3,156 | 9,6147677644539E-27 | RP11-484D2.2 | -4,792 | 9,07155081512579E-33 | SCPEP1 | 2,602 | 8,5991260635047E-38 |
| CTD-2538A21.1 | 11,719 | 9,61335003434808E-28 | MIR4646 | 5,789 | 9,07018051792168E-06 | FAM73A | 2,349 | 8,59687793363751E-10 |
| GPATCH2L | 1,326 | 9,61161730134518E-07 | RP11-663P9.2 | 3,878 | 9,06170344002815E-97 | AL021707.1 | -2,011 | 8,59368290580218E-06 |
| RP11-81A1.3 | 10,847 | 9,6098911725734E-24 | NPIPA5 | -1,447 | 9,05827196525509E-05 | EXOC6 | 2,226 | 8,59109507025342E-14 |
| LRBA | 1,613 | 9,60702374213347E-09 | MIR3652 | -5,614 | 9,05263923507862E-107 | SPG21 | 1,213 | 8,58499837122871E-09 |
| PREPL | 1,115 | 9,60654479421589E-06 | RP11-641A6.5 | 8,238 | 9,04958126781953E-14 | RPS4XP22 | 4,682 | 8,57979760497965E-24 |
| RP11-51L5.2 | 6,567 | 9,60615107857786E-60 | RP11-174E22.2 | 9,296 | 9,04694154057933E-17 | CTD-2228K2.5 | -1,766 | 8,57977068917537E-08 |
| FBL | 1,450 | 9,60210858957773E-26 | FBXO9 | 2,146 | 9,04585568462365E-31 | DDX18P6 | 5,797 | 8,5730967475743E-06 |
| GRHPR | 2,437 | 9,60204862013979E-27 | RP13-270P17.1 | -1,067 | 9,04331700116458E-07 | AC068831.11 | 8,910 | 8,56446632622412E-74 |
| EIF3K | -1,831 | 9,60119743503945E-43 | AKR1B1P2 | 5,228 | 9,03949908963289E-05 | NUP37 | 1,748 | 8,55973329788925E-16 |
| AC016405.1 | 6,022 | 9,59498473810996E-07 | AC004166.6 | -1,431 | 9,0380888788138E-09 | RP11-832N8.1 | 2,915 | 8,55968283691963E-76 |
| SLX1B-SULT1A4 | -1,535 | 9,58302841363916E-10 | NOP58 | 1,946 | 9,03162343061481E-22 | DIMT1 | 1,314 | 8,55849501677339E-09 |
| GMDS-AS1 | 1,924 | 9,57469340669897E-06 | PMP22 | 1,230 | 9,02950778685667E-07 | RP11-172C16.4 | 6,652 | 8,55849501677339E-09 |
| TCEB2P1 | 9,022 | 9,57117513513339E-147 | PABPC1 | -1,745 | 9,02939597834452E-06 | USP8P1 | 5,873 | 8,55489787936735E-07 |
| ACTR3P2 | 8,223 | 9,56539217103111E-14 | RBM18 | 1,651 | 9,02889734268178E-13 | RPL15P3 | -0,777 | 8,55400204280253E-06 |
| EXOG | 1,312 | 9,56429811416522E-07 | CDCA8 | 1,887 | 9,02639602832612E-18 | RP11-23B7.3 | 10,202 | 8,55252926804086E-21 |
| POLG | -1,187 | 9,53546255902354E-07 | RP11-257K9.8 | -1,295 | 9,02039315608688E-06 | PRCP | 1,873 | 8,55187797454389E-15 |
| SUCLG2 | -1,504 | 9,53428664992988E-13 | AC010894.4 | 6,416 | 9,01732957096257E-125 | TOM1L1 | 2,064 | 8,55039092600085E-21 |
| CTD-2200P10.1 | 7,435 | 9,5338112571615E-11 | RP11-528A10.2 | 8,568 | 9,01480809622516E-15 | PSTPIP2 | 2,214 | 8,54547607969995E-09 |
| POLB | 2,192 | 9,52211662706932E-25 | EDNRB | 2,328 | 9,01445779585111E-05 | MIER1 | 1,668 | 8,54397409303836E-11 |
| HERC4 | 1,712 | 9,52046141588372E-07 | GDI2P2 | 9,973 | 9,01158338922391E-20 | RP5-854E16.2 | 2,851 | 8,54397409303836E-11 |
| FASN | -1,783 | 9,52046141588372E-07 | NPEPPS | 2,021 | 9,01158338922391E-20 | RPL36AP26 | -1,533 | 8,54132901272173E-05 |
| DPY19L2P1 | 2,374 | 9,51930787416002E-06 | EEF1A1P24 | -2,567 | 9,00944182373479E-09 | RP11-453D16.2 | 4,800 | 8,54074244712581E-12 |
| RPS17L | -5,762 | 9,51763100831947E-132 | CDC42 | -1,948 | 9,00129708648476E-47 | TCEA1 | -1,882 | 8,54026001464378E-07 |
| AC007389.1 | 8,947 | 9,51603993967796E-16 | RP11-630I5.1 | -1,917 | 9,00060202706704E-12 | ANKRD36C | 1,916 | 8,53937917106293E-11 |
| PRICKLE3 | -1,002 | 9,5131588236415E-10 | RP11-336F14.1 | 8,896 | 8,99900188897769E-16 | GORASP2 | 1,887 | 8,53086925379433E-14 |
| DNAJC25-GNG10 | -1,076 | 9,50250599201774E-12 | RP1-180M12.1 | 12,070 | 8,99892035307882E-30 | RPL13P12 | -0,636 | 8,52905871182871E-05 |
| PTMA | -2,951 | 9,49408776481066E-42 | | | | | | |
